# Sex differences and immune correlates of Long COVID development, persistence, and resolution

**DOI:** 10.1101/2024.06.18.599612

**Authors:** Rebecca E. Hamlin, Shaun M. Pienkos, Leslie Chan, Mikayla A. Stabile, Kassandra Pinedo, Mallika Rao, Philip Grant, Hector Bonilla, Marisa Holubar, Upinder Singh, Karen B. Jacobson, Prasanna Jagannathan, Yvonne Maldonado, Susan P. Holmes, Aruna Subramanian, Catherine A. Blish

**Author notes:** These authors contributed equally to this work (Rebecca E. Hamlin, Shaun M. Pienkos).

## Abstract

Sex differences have been observed in acute COVID-19 and Long COVID (LC) outcomes, with greater disease severity and mortality during acute infection in males and a greater proportion of females developing LC. We hypothesized that sex-specific immune dysregulation contributes to the pathogenesis of LC. To investigate the immunologic underpinnings of LC development and persistence, we used single-cell transcriptomics, single-cell proteomics, and plasma proteomics on blood samples obtained during acute SARS-CoV-2 infection and at 3 and 12 months post-infection in a cohort of 45 patients who either developed LC or recovered. Several sex-specific immune pathways were associated with LC. Specifically, males who would develop LC at 3 months had widespread increases in *TGF-β* signaling during acute infection in proliferating NK cells. Females who would develop LC demonstrated increased expression of *XIST*, an RNA gene implicated in autoimmunity, and increased *IL1* signaling in monocytes at 12 months post infection. Several immune features of LC were also conserved across sexes. Both males and females with LC had reduced co-stimulatory signaling from monocytes and broad upregulation of *NF-κB* transcription factors. In both sexes, those with persistent LC demonstrated increased LAG3, a marker of T cell exhaustion, reduced *ETS1* transcription factor expression across lymphocyte subsets, and elevated intracellular IL-4 levels in T cell subsets, suggesting that ETS1 alterations may drive an aberrantly elevated Th2-like response in LC. Altogether, this study describes multiple innate and adaptive immune correlates of LC, some of which differ by sex, and offers insights toward the pursuit of tailored therapeutics.

**One Sentence Summary:** This multi-omic analysis of Long COVID reveals sex differences and immune correlates of Long COVID development, persistence, and resolution.

## INTRODUCTION

SARS-CoV-2, the virus that causes Coronavirus Disease 19 (COVID-19), has thus far led to over 770,000,000 reported COVID-19 cases and over 7,000,000 deaths globally since its first reported case in December, 2019 (*1*). In addition to the morbidity and mortality caused by acute disease, a significant proportion of patients (approximately 10-20%) continue to suffer from symptoms for months-to-years after SARS-CoV-2 infection (*2*). This condition carries multiple names (i.e. Long COVID, Post-COVID Condition(s), and Post-acute sequelae of SARS-CoV-2 infection (PASC)) and has been defined as the persistence or development of clinical signs or symptoms attributable to SARS-CoV-2, either 4- or 12-weeks post infection (*2–4*).

For this study, we use the term Long COVID (LC) and the World Health Organization (WHO) definition for symptoms persisting beyond 12-weeks post infection (*2*). LC is a heterogeneous condition with potential clinical signs and symptoms spanning nearly every organ system, which commonly include: fatigue, myalgias, dyspnea, cough, chest pain, neurocognitive dysfunction, dysosmia, and/or dysgeusia (*2, 5, 6*). Given the heterogeneity of clinical presentations, LC may encompass potential subgroups of patients afflicted by different symptomatology and pathophysiology, although this continues to be an area of active research investigation without clear consensus (*7–11*).

There are a number of proposed, not mutually exclusive, pathophysiologic mechanisms for LC. These include immune dysregulation (*11–30*), SARS-CoV-2 persistence (*31–47*), SARS-CoV-2-specific (*11, 12, 24, 31, 48–51*) and auto-reactive antibody responses (*11, 52–58*), human herpesvirus (e.g. Epstein-Barr Virus (EBV)) reactivation (*11, 12, 59–62*), microvascular and clotting dysfunction (*63–67*), thromboinflammation (*62*), amyloid deposition (*63, 68–70*), mitochondrial dysfunction (*68, 71, 72*), dysbiosis (*73–76*), and hormonal or metabolic abnormalities, such as cortisol (*11, 12*) or serotonin (*77*) dysregulation. Prior literature on immune dysregulation in LC clinical cohorts have primarily investigated immune cell frequencies, single-cell cytokine production, plasma proteomics, and SARS-CoV-2 antibody responses, with variable results across studies and cohorts (*11–15, 17–19, 21–27, 29, 31, 48, 50, 78, 79*). Several studies have implicated different immune cell populations in the pathophysiology of LC, but a unifying etiology across different studies has not been discovered (*11, 12, 14, 22, 24, 51*).

Intriguingly, sex-based differences have underpinned outcomes for both acute COVID-19 and LC. Within the first year of the pandemic, males were observed to have higher rates of critical illness and death compared to females from acute COVID-19 (*80–86*). Conversely, female sex has been epidemiologically associated with increased risk of developing LC (*17, 87–94*). While the exact mechanisms underlying these sex differences remain to be fully elucidated, several innate and adaptive immunologic differences have been noted between the sexes (*95, 96*). Following symptomatic COVID-19, females were found to have higher percentages of CD19^+^ B cells and CD4^+^ T-cells compared to males, while males were noted to have higher inflammatory markers (i.e. C-reactive protein and IL-6) compared to females (*97*). Males were also found to have slower SARS-CoV-2 viral clearance than females, which may contribute to differences in acute COVID-19 outcomes (*98*). While there is a similarity between the higher prevalence of females with LC and autoimmunity (*95, 96, 99*), the mechanisms underlying sex differences in LC require further research investigation.

We hypothesized that sex-specific, dysregulated immune cell communication during both acute COVID-19 and convalescence contribute to the pathogenesis of LC. To investigate this hypothesis, we performed comprehensive multi-omic profiling in 45 participants from a cohort of patients who were recruited at 3 months post infection to study recovery from SARS-CoV-2 infection. Many subjects had persistent LC symptoms at 3 and 12 months post infection, affording us the opportunity to evaluate predictors of LC versus recovery. We performed single cell RNA-sequencing (scRNA-seq) and proteomic analyses, focusing our analyses on networks of gene activation and inferred cell-cell communication pathways to gain novel insights into the pathways associated with LC.

## RESULTS

### Cohort description

Multiple assays were performed on samples from 45 participants from the prospective, observational patient cohort called “Infection Recovery in SARS-CoV-2” (IRIS) at Stanford University to evaluate immune function during recovery from COVID-19. Demographics of the cohort as assessed at the enrollment visit at 3 months after infection are presented in Table S1.

At 3 months post infection, 9 participants had fully recovered and 36 had persistent symptoms indicative of LC. Of the 9 participants who recovered, 44.4% were female, the mean age was 41.7 years, and 1 participant (11.1%) was previously hospitalized for acute SARS-CoV-2 infection (Table S1). Of the 36 participants who developed LC at 3 months post infection, 55.6% were female, the mean age was 43.0 years, and 8 participants (22.2%) were previously hospitalized for acute SARS-CoV-2 infection (Table S1). Participant demographics stratified by sex and LC status are also shown (Table S1). Symptom prevalence for these subjects is shown in fig. S1A, and demonstrated that a variety of symptoms were present among both male and female participants, with constitutional, cognitive, and pulmonary symptoms the most prevalent.

Banked peripheral blood mononuclear cell (PBMC) samples from these subjects during acute infection, at 3 months post infection, and at 12 months post infection were used for scRNA-seq and cytometry by time-of-flight (CyTOF) (Fig. 1A). Olink proteomics was performed on plasma samples (Fig. 1A). For scRNA-seq data, cell types were annotated with Seurat v4 PBMC multimodal reference mapping (*100*) to identify multiple innate and adaptive cell types. Within individual cell types, we assessed expression changes in gene modules and inferred ligand-receptor interaction analyses with target gene predictions (Fig. 1B). CyTOF and plasma proteomic data were used to validate findings from transcriptional analyses.

**Fig. 1.**
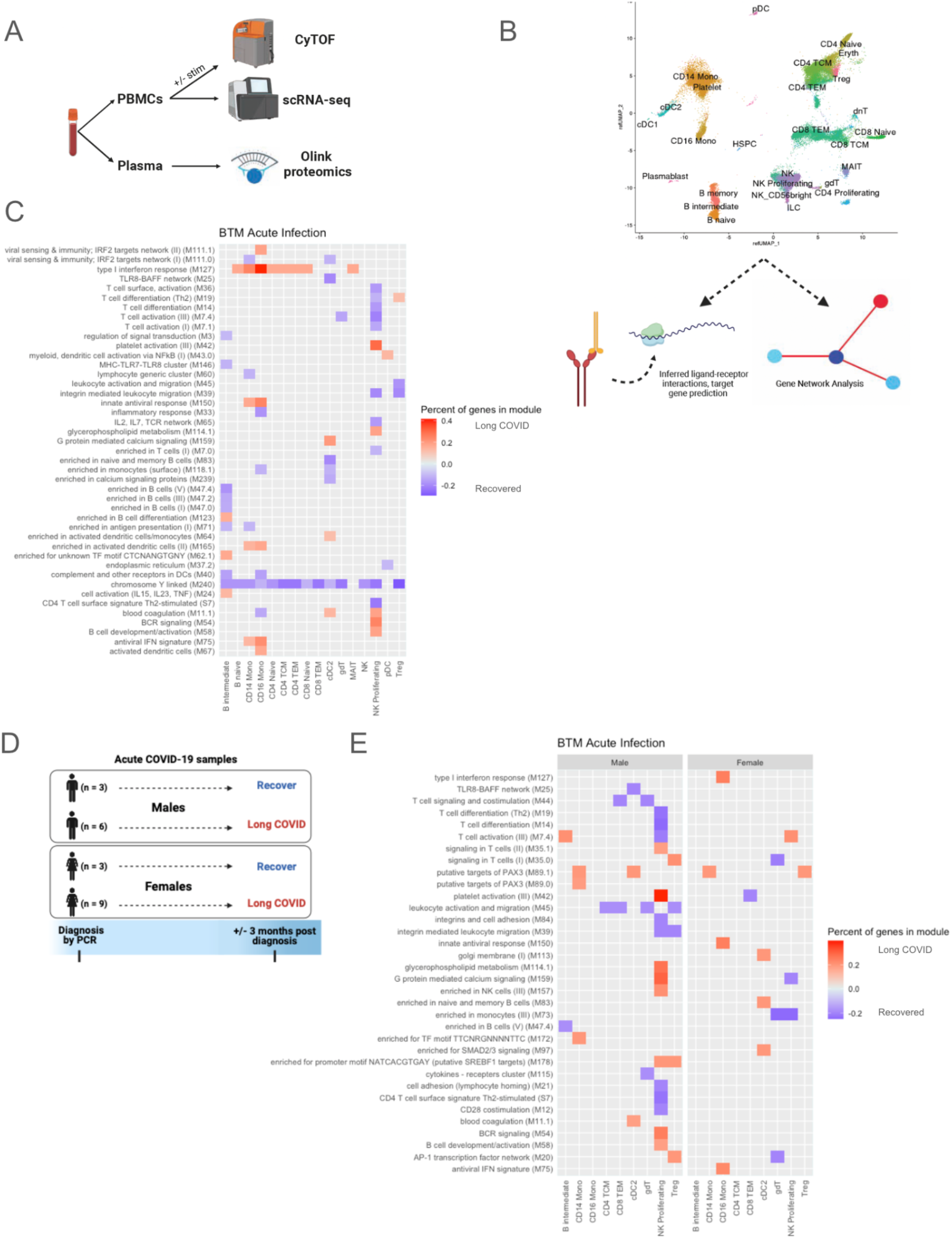
Experimental design, cell clustering and blood transcription modules (BTM) differences during acute infection. **(A)** Experimental design demonstrating use of CyTOF and scRNA-seq for PBMC samples and Olink proteomics for plasma samples. **(B)** UMAP of scRNA-seq data (subsampled to 100,000 cells to reduce overplotting). Individual cell types were further analyzed with MultiNicheNet for cell-cell communication inference and BioNet for gene network analysis. **(C)** BTMs of samples during acute infection that differentiated subjects with LC versus those who recovered by 3 months post infection for males and females combined. Only modules in which >10% of the constituent genes had an absolute log2 fold-change (log2FC) >0.25 are plotted. Red tiles indicate higher expression in subjects who developed LC, and blue tiles indicate lower expression. **(D)** Subjects included in acute SARS-CoV-2 infection analysis, separated by sex and LC versus recovered status 3 months post infection. **(E)** BTMs during acute infection that differentiated those with LC versus those who recovered by 3 months post infection. Males and females are plotted separately. The top 50 modules, by percent of genes, in which >10% of the constituent genes had an absolute log2 fold-change (log2FC) >0.25, are plotted. Red tiles indicate higher expression in subjects who developed LC, and blue tiles indicate lower expression.

### Sex differences in blood transcription modules during acute SARS-CoV-2 infection

To identify immune dysregulation during acute infection that might predict LC, we performed transcriptional and proteomic analyses on the 21 subjects (of the original 45) who had samples available during acute infection (Table S2). Among these 21 subjects, 15 patients went on to develop LC at 3 months post infection, and 6 patients fully recovered by 3 months post infection (Table S2). Of the 6 participants who would recover, 50% were female, the mean age was 41.8 years, and none were hospitalized for acute SARS-CoV-2 infection (Table S2). Of the 15 participants who would develop LC, 60% were female, the mean age was 45.8 years, and 1 participant (6.7%) was hospitalized for acute SARS-CoV-2 infection (Table S2). Participant demographics stratified by sex and LC status are also shown (Table S2).

We first sought to identify transcriptional predictors of the development of LC using blood transcription modules (BTMs) sourced from a reference gene set database dedicated to studying blood transcriptional immune responses (*104*). Numerous BTMs were different during acute infection in patients who would later develop LC versus those who would recover at 3 months post infection. In particular, the “type I interferon response” module, which is critical for antiviral immune responses (*102*), was increased in acute samples of patients who would develop LC in various monocyte, B cell, and T cell subsets (Fig. 1C). It was also noteworthy that the BTM “chromosome Y linked” was decreased in patients who would develop LC across many cell types (Fig. 1C, Fig S2A). The genes driving this BTM difference were *XIST*, *DDX3Y*, *UTY/KDM6C*, *KDM5D*, *PRKY,* and *USP9Y*, which are sex-linked genes thought to have important roles in immune responses (*103–108*) (Table S3). A greater proportion of females than males in the cohort progressed to LC after acute infection (75% vs 67%, respectively), accounting for a portion of these sex-linked gene expression differences when the sexes are analyzed together.

When gene expression differences were separated by sex (Fig. 1D), expression of *DDX3Y, UTY, PRKY,* and *USP9Y* differed in males who progressed to LC compared to those who recovered (fig. S2B). *DDX3Y*, the Y-linked homolog of *DDX3X* with known roles in the interferon (IFN) response to pathogens (*104*), was notably downregulated across most cell subsets at 12 months in males with LC. In females, the RNA gene *XIST*, which has previously been implicated in autoimmunity (*103*), had higher expression in several innate and adaptive immune cell subsets in those who developed LC (fig. S2C). At 3 and 12 months after acute infection, increases in *XIST* in females with LC were still present but less prominent (fig. S2C).

Based on these findings and the evidence for sex-related differences in COVID-19 outcomes (*17, 80–94, 97, 98*), we next analyzed the sexes separately (Fig. 1D). This analysis revealed stark differences between males and females in the acute infection gene profiles of those who went on to develop LC versus those who recovered, with prominent gene expression differences in multiple BTMs in proliferating NK cells and reduced expression in the “leukocyte activation and migration” module for several T cell subsets in males who developed LC (Fig. 1E). In contrast, females who developed LC had expression changes which were less restricted to particular cell types or modules, affecting multiple lymphoid and myeloid subsets (Fig. 1E). Increased expression of BTMs in CD14^+^ and CD16^+^ monocytes was found in both sexes of those who developed LC (Fig. 1E). Given these sex-specific differences found in BTM analyses of acutely infected samples, subsequent analyses of these samples focused on proliferating NK cells, monocytes, and sex-specific differences in several key regulators of inflammation.

### Acute TGFβ1 signaling in male proliferating NK cells predict LC

To understand the drivers of the transcriptional changes observed in BTM analysis (Fig. 1E), we used MultiNicheNet to infer communication between proliferating NK cells and other immune cells during acute infection (*109*). The majority of the top ligand-receptor interactions with proliferating NK cells as senders of signal in acutely infected male samples was dominated by Transforming Growth Factor-β1 (*TGFβ1*) signaling, as well as IFNɣ signaling, in those who would develop LC compared to those who would recover (Fig. 2A).

**Fig. 2.**
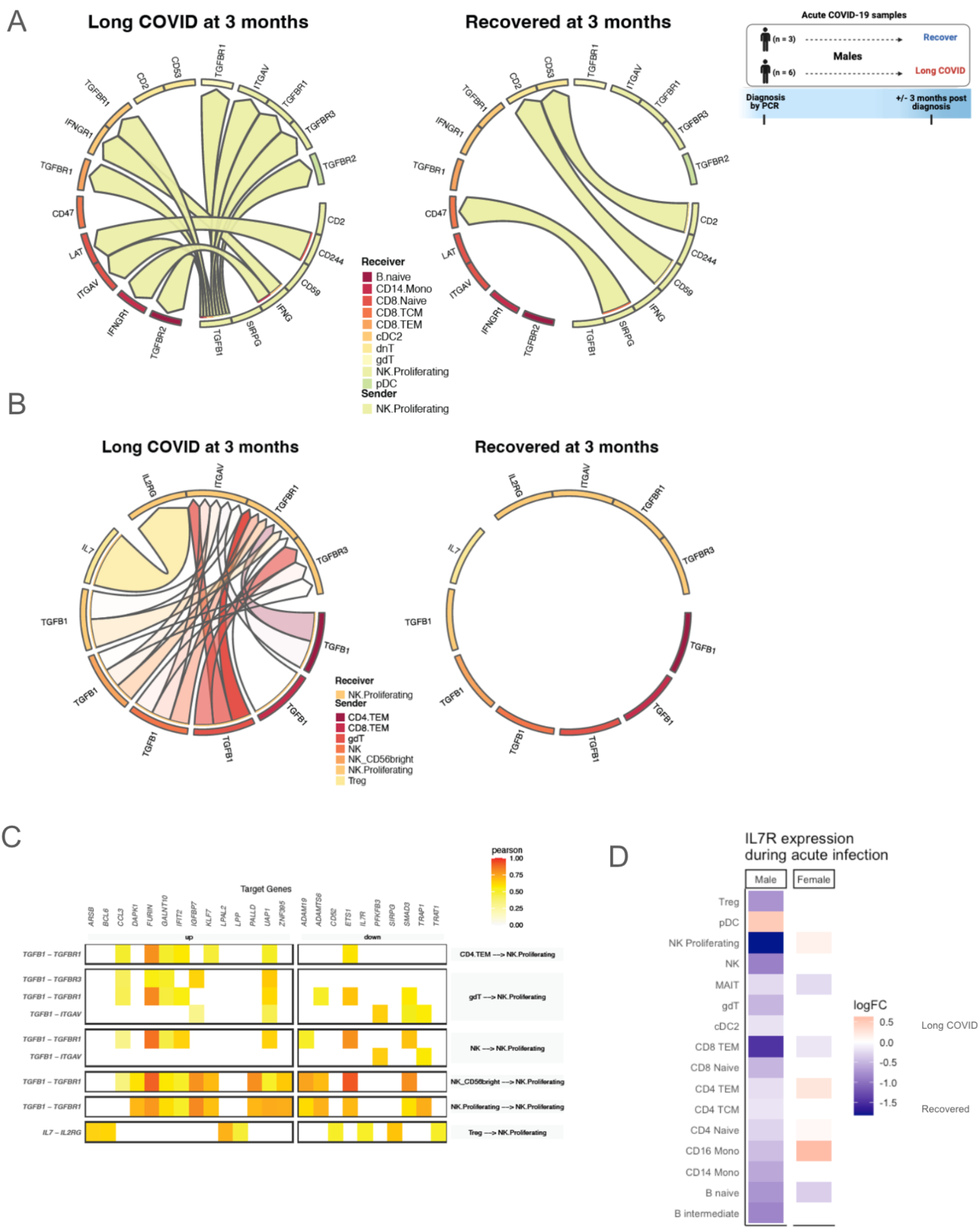
Proliferating NK cells in males during acute infection. **(A-B)** Predicted ligand-receptor interactions involving proliferating NK cells, separated by development of LC symptoms 3 months after acute infection versus resolution of symptoms by 3 months, with **(A)** showing signaling sent from proliferating NK cells, and **(B)** showing signaling received by proliferating NK cells. **(C)** Target genes with altered expression in proliferating NK cells of acutely infected male subjects who will develop LC at 3 months, with associated ligand-receptor interactions. Tile color in the heatmap indicates the Pearson correlation coefficient of expression of the ligand-receptor pair and the target gene. **(D)** Pseudobulk interleukin-7 receptor (*IL7R*) expression by cell type during acute infection, separated by sex. Tile color in the heatmap indicates log2FC, with higher expression in those who will develop LC at 3 months post infection indicated by red and lower expression indicated by blue.

Similarly, most of the top interactions received by proliferating NK cells were dominated by *TGFβ1* signaling from several NK cell and T cell populations in acute samples from males who would develop LC (Fig. 2B). Such signaling was predicted to result in the upregulation of: the chemokine *CCL3*, which can augment NK cell cytolytic activity and can be produced by activated NK cells to recruit other effector cells (*110*); *DAPK1*, which is involved in IFNɣ-induced apoptosis and may promote NK cell killing (*111, 112*); interferon-induced *IFIT2*, which can promote apoptosis (*113*); *ZNF395*, which can activate interferon-stimulated genes (*114*); the protease *FURIN*, which is also a TGFβ1-converting enzyme (*115*); *IGFBP7*, which can upregulate TGFβ1 signaling (*116*); and *GALNT10*, which may be associated with an immunosuppressive phenotype (*117*); among others (Fig. 2C). This TGFβ1 signaling was also predicted to downregulate: *ADAMTS6*, which can increase TGFβ1 activation (*118*); *SMAD3*, which is an intracellular signaling component of the TGFβ pathway but has previously been shown to be down-regulated in response to TGFβ in certain contexts (*119*); *TRAP1*, which is suggested to have cytoprotective effects against oxidative stress and reactive oxygen species (*120, 121*); and *ETS1*, which regulates NK cell development and terminal differentiation (*122*); among others (Fig. 2C). Though *TGFβ1* is best described as an inhibitory signal in NK cells, these transcriptional differences also suggest that cytolytic functions and antiviral responses may be altered by *TGFβ1 s*ignaling in this context.

In addition to *TGFβ1* signaling, MultiNicheNet also showed upregulation of IL-7 signaling in acutely infected males who will develop LC, which is important for NK cell survival and cytotoxicity (*123*) (Fig. 2B). This was predicted to result in downregulation of *IL7R*, suggesting a negative regulatory mechanism (Fig. 2C) (*124*). IL7R expression during acute infection was indeed downregulated in males who will develop LC in the majority of cell types, including proliferating NK cells, and this pattern was not observed in females (Fig. 2D). The predicted effects on downstream target genes in proliferating NK cells were also apparent and consistent across individual patients (fig. S3).

Unlike in males, cell-cell communication analyses of proliferating NK cells in acutely infected females who would develop LC demonstrated signaling predominantly involving *CCL5, CD99*, *ADAM10*, *IL15*, and *NRG1*, among others, but did not demonstrate predominant *TGFβ1* signaling (fig. S4).

Given the prominent *TGFβ1* signaling in proliferating NK cells of the acutely infected males who subsequently developed LC, we next examined whether *TGFβ1* was important across different cell types and time points.

### TGFβ1 expression and protein secretion differs in the development and persistence of LC by sex

During acute infection, *TGFβ1* expression was increased in males who would develop LC not only in proliferating NK cells, but also in the majority of innate and adaptive cell types (Fig. 3A). On the other hand, *TGFβ1* expression was decreased during acute infection in females who would develop LC in the majority of examined cell types (Fig. 3A). Corroborating these findings at the protein level, latency-associated peptide (LAP) TGFβ1 was significantly increased in the plasma of acutely infected males who would develop LC compared to males who would recover; this difference was not observed in females (Fig. 3B).

**Fig 3.**
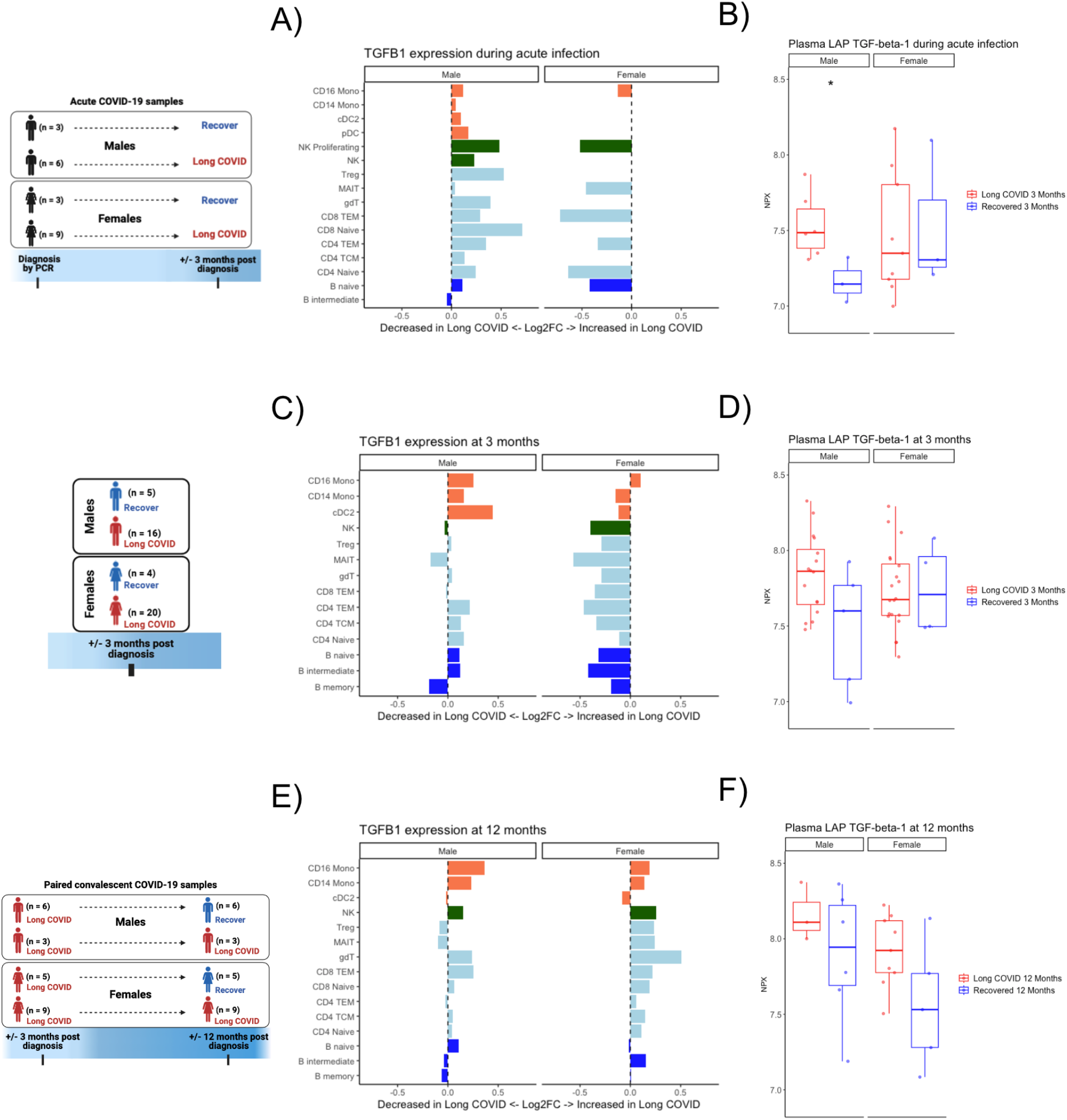
TGF-β in transcriptomic and plasma proteomics analyses differs by sex in those with LC. **(A)** Log2FC of pseudobulk expression of *TGFβ1* gene during acute infection, separated by sex. Bars moving to the right of the dashed line indicate increased expression and bars moving to the left indicate reduced expression in those who will develop LC at 3 months post infection. Bar colors correspond to major cell types. **(B)** Plasma proteomic measurement of LAP TGFβ1 during acute infection, separated by sex. * indicates unadjusted p<0.05 between those who will develop LC vs. recover at 3 months post infection. **(C)** Log2FC of pseudobulk expression of *TGFβ1* gene 3 months after acute infection, separated by sex. Bars moving to the right of the dashed line indicate increased expression and bars moving to the left indicate reduced expression in those with LC at 3 months post infection. Bar colors correspond to major cell types. **(D)** Plasma proteomic measurement of LAP TGFβ1 in those with LC vs. those who recovered at 3 months post infection, separated by sex. **(E)** Log2FC of pseudobulk expression of *TGFβ1* gene 12 months after acute infection, separated by sex. Bars moving to the right of the dashed line indicate increased expression and bars moving to the left indicate reduced expression in those with persistent LC at 12 months. Bar colors correspond to major cell types. **(F)** Plasma proteomic measurement of LAP TGFβ1 in those with persistent LC vs. recovery at 12 months post infection, separated by sex.

We next evaluated transcriptional and proteomic differences in all study participants at 3 months post infection between 36 individuals who developed LC versus 9 individuals who recovered (Table S1), with the analysis divided by sex (Fig. 3C). Similar to the findings during acute infection, *TGFβ1* expression was increased in males with LC in the majority of examined cell types, while *TGFβ1* expression was mostly decreased in the different cell types in females with LC (Fig. 3C). At the plasma protein level, LAP TGFβ1 appeared to be increased in males with LC compared to recovered males and similar in females with LC compared to recovered females, but these differences were not statistically significant (Fig. 3D).

Additionally, a subgroup of the total 45 participants had samples available at both 3 and 12 months post infection to enable evaluation for LC persistence (Table S4). This analysis involved 12 participants who had persistent LC at both 3 and 12 months post infection and 11 participants who had LC at 3 months but recovered at 12 months post infection. Of the 12 participants with persistent LC at 3 and 12 months, 75% were female, the mean age was 44.8 years, and 1 participant (8.3%) was previously hospitalized for acute SARS-CoV-2 infection (Table S4). Of the 11 participants who recovered at 12 months, 45.5% were female, the mean age was 37.3 years, and 3 participants (27.3%) were previously hospitalized for acute SARS-CoV-2 infection (Table S4). Participant demographics stratified by sex and LC status are also shown (Table S4). LC symptoms for this group of patients at 12 months are shown in fig. S1B and were notable for increased prevalence of cognitive, constitutional, neurologic, and pulmonary symptoms in those with persistent symptoms. We evaluated the LC versus recovered groups separately by sex.

*TGFβ1* expression was again found to be increased in males with persistent LC at 12 months post infection in the majority of cell types (Fig. 3E). Interestingly, unlike earlier time points, *TGFβ1* expression was also increased in females with persistent LC at 12 months post infection in the majority of cell types (Fig. 3E). Secreted LAP TGFβ1 in plasma also appeared to be increased in both males and females with persistent LC at 12 months compared to the recovered males and recovered females, respectively, but these differences did not reach statistical significance (Fig. 3F). Altogether, these data suggest that increased transcriptional and proteomic TGFβ1 are predictive of LC development and markers of persistent LC in males, but only at 12 months post infection are there hints of increased TGFβ1 playing a role in LC in females.

### Differential signaling by sex in monocytes during acute infection predicts LC

Due to the differences of BTMs in monocytes during acute infection (Fig 1C, 1E), we further investigated these cells in both sexes with cell-cell communication and gene network analyses. In males, inference of ligand-receptor signaling toward monocytes showed that CD40 ligand signaling was most different during acute infection in those who went on to develop LC versus those who recovered (Fig. 4A). CD14^+^ monocytes and CD16^+^ monocytes of subjects who would recover received CD40 ligand signaling from T cells, targeting multiple integrins that act to facilitate CD40 signaling (*125–127*), an important stimulus for inflammatory responses in infection and expression of co-stimulatory signals (*128, 129*). CD14^+^ monocytes in males who developed LC received proinflammatory *IFNG* and *IL15* signaling (*130, 131*) as well as pro-apoptotic *TNFSF10* (aka *TRAIL*) signaling (*132*) (Fig. 4A). CD16^+^ monocytes in males who developed LC received CD226 signaling from NK and T cells directed to stimulate *PVR* and *Nectin2* (Fig. 4B). *PVR* encodes CD155, an inhibitory protein which has previously been associated with poor antiviral responses to SARS-CoV-2 (*133*). Additionally, *PVR* and *Nectin2* both promote tissue infiltration of monocytes (*134*), which has been associated with deleterious local inflammation (*135*). Thus, during acute infection, monocytes in males who will recover received CD40 ligand signaling that may promote co-stimulatory signals, while monocytes in males who will develop LC may be more prone to tissue infiltration.

**Fig 4.**
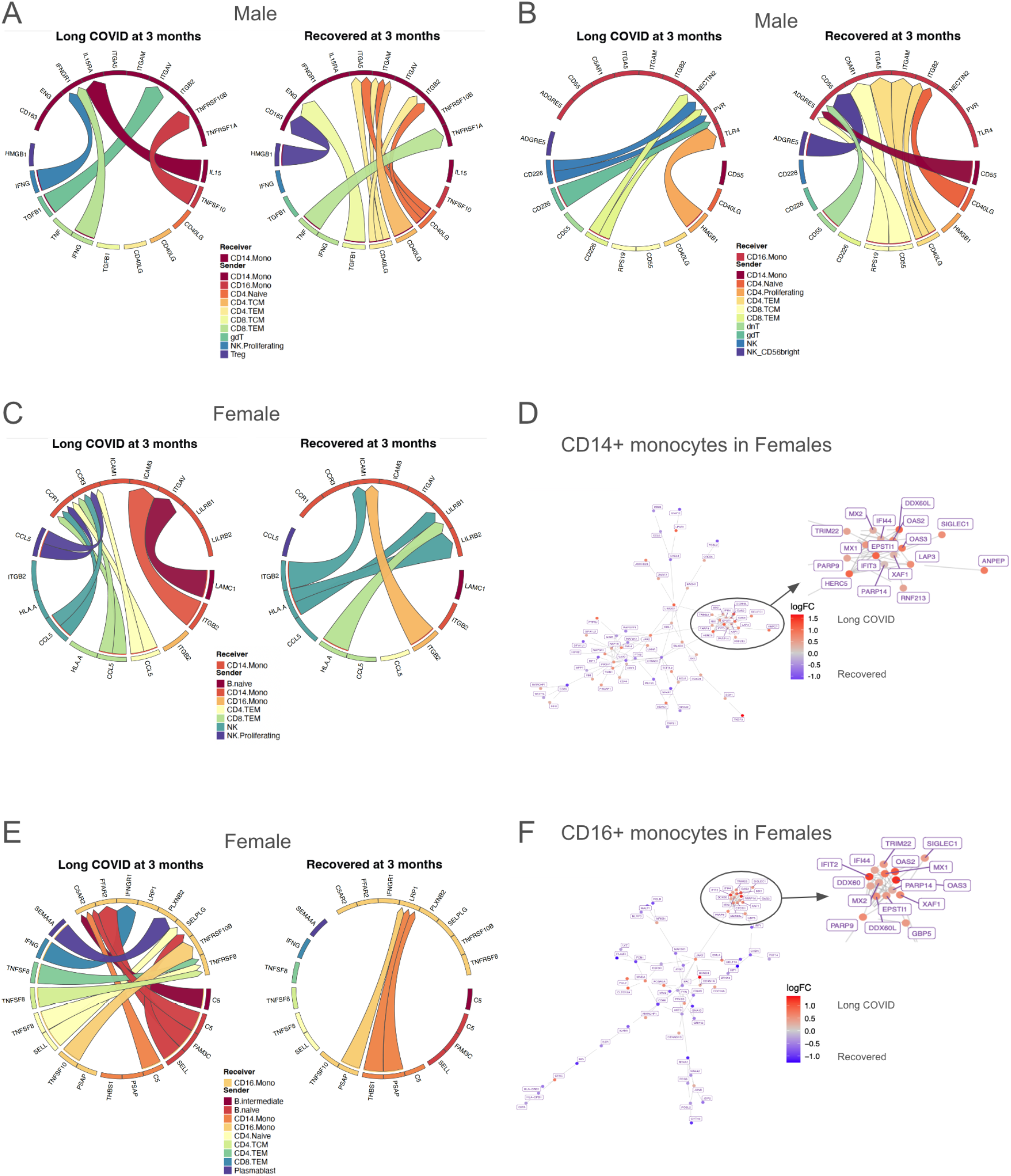
Sex-specific differences in inflammatory monocytes during acute infection. **(A-B)** Inferred cell-cell communication, separated by development of LC symptoms 3 months after acute infection versus resolution of symptoms by 3 months, toward **(A)** CD14^+^ monocytes and **(B)** CD16^+^ monocytes in males. **(C)** Cell-cell communication inference, separated by development of LC symptoms 3 months after acute infection versus resolution of symptoms by 3 months, toward CD14^+^ monocytes in females. **(D)** Differential gene network in CD14^+^ monocytes in females during acute infection. Red nodes indicate higher expression (log2FC) in those who will develop LC, and blue nodes indicate lower expression. **(E)** Cell-cell communication inference, separated by development of LC symptoms 3 months after acute infection versus resolution of symptoms by 3 months, toward CD16^+^ monocytes in females. **(F)** Differential gene network in CD16^+^ monocytes in females during acute infection. Red nodes indicate higher expression (log2FC) in those who will develop LC, and blue nodes indicate lower expression.

In females, CD14^+^ monocytes during acute infection in those who went on to develop LC received increased CCL5 signaling from NK and T cell subsets, and the monocytes themselves show increased expression of chemokine receptors *CCR1* and *CCR3* (Fig. 4C), a combination of which would be expected to promote chemotaxis. Interestingly, higher levels of *CCR* genes in monocytes and macrophages has been associated with increased acute disease severity in COVID-19 patients (*136*). Gene network analysis for CD14^+^ monocytes in females shows a subnetwork of interferon-stimulated and antiviral genes such as *MX1, MX2, TRIM22, OAS2, OAS3, IFI44, DDX60L*, *EPSTI1*, *PARP14*, and *IFIT3* (*137–144*) (Fig. 4D), which are upregulated in those who developed LC.

CD16^+^ monocytes in females who went on to develop LC received increased TNF signaling compared to those who recovered, including *TNFSF8* (aka *CD30L*) (Fig. 4E), which induces NF-κB activation and is implicated in autoimmunity (*145*). Additionally, these monocytes received complement C5 signaling, and complement dysregulation has previously been implicated in LC pathogenesis (*62, 146*). Gene network analysis showed upregulated interferon-stimulated and antiviral genes including *PARP14*, *EPSTI1*, *OAS2, OAS3*, *MX1*, *DDX60*, *TRIM22*, *IFIT2*, and *IFI44* (*137, 139–144, 147*) (Fig. 4F). Interestingly, *IFI44*, which is upregulated in gene networks for both CD14^+^ and CD16^+^ monocytes in females, has been observed to reduce interferon responses and has been associated with immune evasion in SARS-CoV-2 (*148, 149*).

In sum, transcriptional analyses of monocytes during acute infection revealed signatures suggestive of increased potential to infiltrate target tissues in subjects with LC, though the mechanisms differed by sex with males showing *CD226* to *NECTIN2/PVR* signaling and females showing *CCL5* to *CCR1/CCR3* signaling. Additionally, there was upregulation of interferon-stimulated genes in females who developed LC, suggesting a more activated monocyte phenotype which may promote local tissue inflammation.

### Pro-inflammatory monocytes with reduced co-stimulatory function in LC

We next turned our attention to the immune status at 3 and 12 months post infection. BTM analysis at 3 and 12 months in both males and females with persistent LC revealed ongoing differences in monocytes between subjects with persistent LC compared to those who recovered by 12 months (Fig. 5A-B). Of particular interest, modules associated with chemokines and inflammatory molecules as well as chemokine receptor signaling were upregulated in CD14^+^ monocytes at 3 months in those with persistent LC (Fig. 5A). At 12 months post infection, males with persistent LC showed upregulation of numerous BTMs in CD14^+^ monocytes, including modules associated with proinflammatory myeloid cell responses, chemokines, NF-κB activation, and antigen presentation, among others (Fig. 5B). BTM analysis of CD16^+^ monocytes in females also showed upregulation in BTMs associated with proinflammatory myeloid cell responses, NF-κB activation, and chemokines in those with persistent LC at 12 months (Fig. 5B). CD14^+^ monocytes in females with LC at 12 months also showed upregulation of the module associated with chemokines and inflammatory molecules (Fig. 5B).

**Fig. 5.**
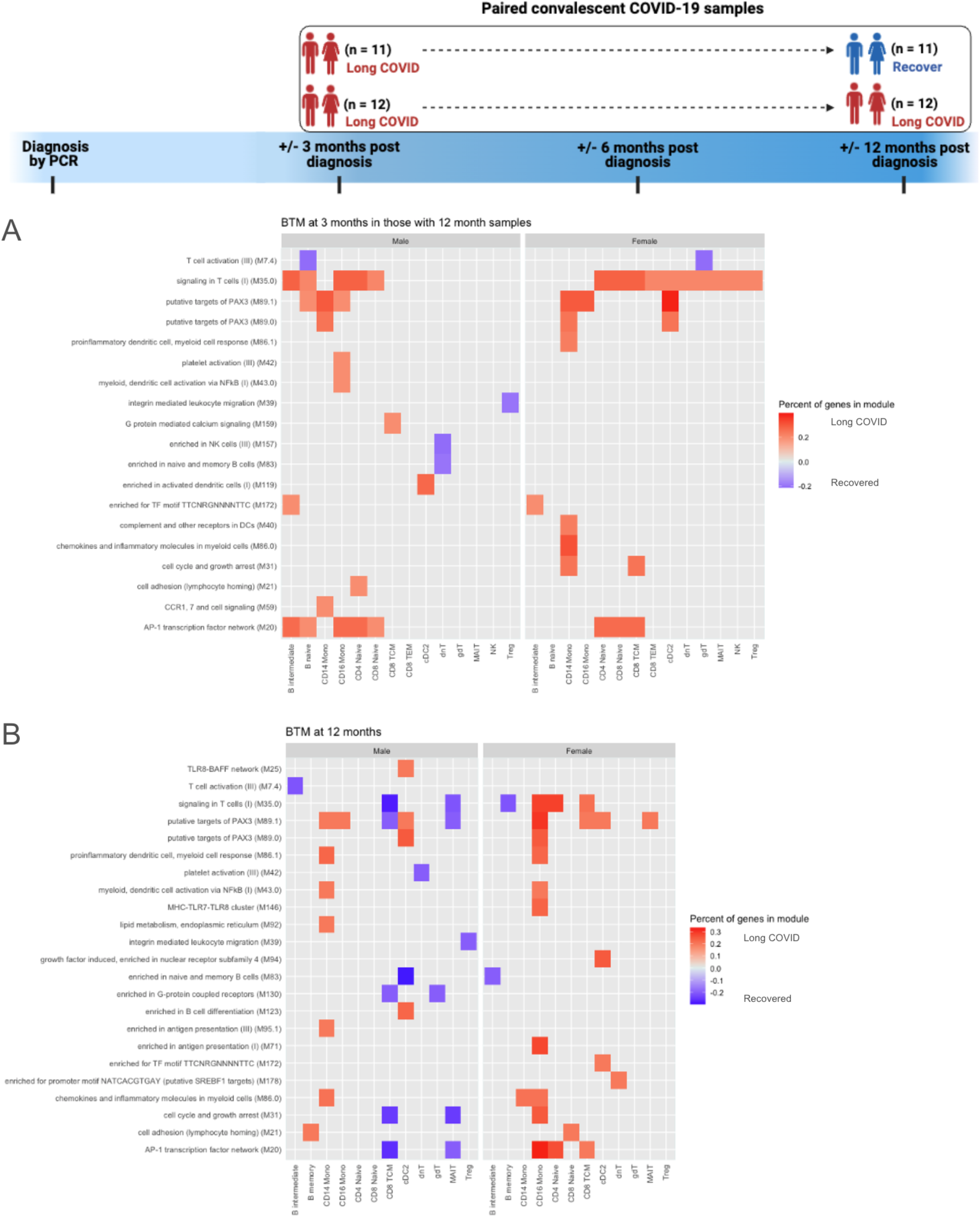
BTMs at 3 and 12 months after acute SARS-CoV-2 infection. **(A)** BTMs 3 months after acute infection, separated by sex, which differentiate those with persistent LC at 12 months post infection versus those who will recover by 12 months. **(B)** BTMs 12 months after acute infection, separated by sex, which differentiate those with persistent LC at 12 months post infection versus those who recovered between 3 and 12 months. The top 50 modules, by percent of genes, in which >10% of the constituent genes had an absolute log2 fold-change (log2FC) >0.25, are plotted. Red tiles indicate higher expression in subjects with LC, and blue tiles indicate lower expression.

Cell-cell communication analysis of CD14^+^ monocytes in males revealed much greater activity in subjects with LC at 12 months. As senders of signal, activating and proinflammatory signals such as *CD58* and *IL-1𝛂* toward regulatory T cells and *S100A12* toward CD16^+^ monocytes and conventional dendritic cells 2 (cDC2) are prioritized in those who have LC (Fig 6A). Conversely, *IL1RN (*the anti-inflammatory antagonist of IL1) (*150*) signaling toward cDC2 was also upregulated in those with LC (Fig 6A). As receivers, CD14^+^ monocytes in those with LC are targets of *TGFβ1* signaling from NK cells and multiple T cell subsets (Fig 6B), consistent with our finding that TGFβ1 transcription is upregulated across all time points in male subjects who developed LC (Fig 3E).

**Fig. 6.**
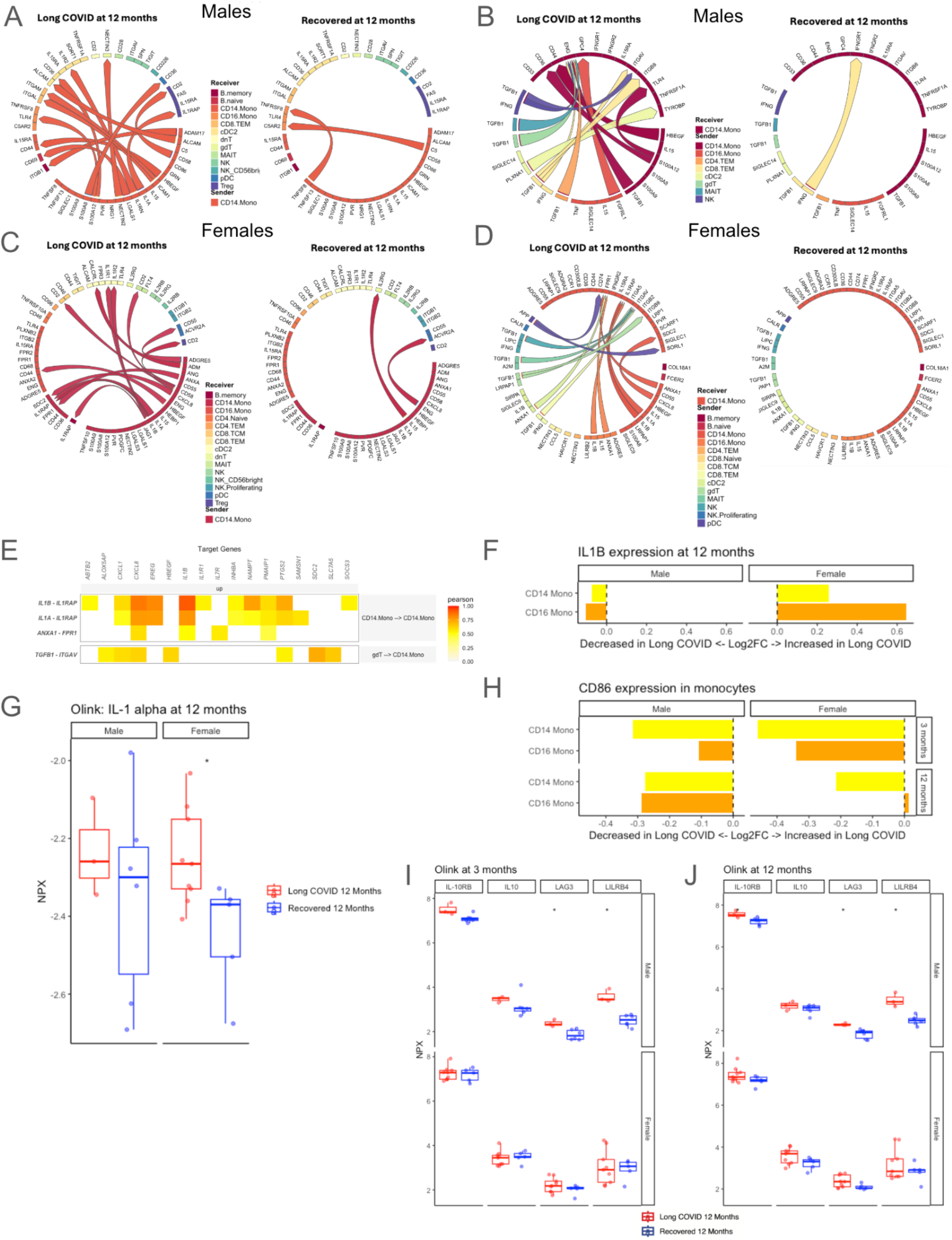
Cell-cell communication of CD14^+^ monocytes, co-stimulatory potential, and markers of T cell exhaustion in persistent LC. **(A-B)** Inferred communication **(A)** originating from, and **(B)** going toward CD14^+^ monocytes in males at 12 months post infection, comparing those with persistent LC versus those who recovered by 12 months. **(C-D)** Inferred cell-cell communication **(C)** originating from, and **(D)** going toward CD14^+^ monocytes in females at 12 months post infection, comparing those with persistent LC versus those who recovered by 12 months. **(E)** Target genes with increased expression in CD14^+^ monocytes of females with LC at 12 months, with associated ligand-receptor interactions. Tile color in the heatmap indicates the Pearson correlation coefficient of expression of the ligand-receptor pair and the target gene. **(F)** Log2FC of pseudobulk *IL-1β* expression in CD14^+^ and CD16^+^ monocytes at 12 months post infection, comparing those with persistent LC and those who recovered by 12 months, separated by sex. Bars moving to the right of the dashed line indicate increased expression and bars moving to the left indicate reduced expression in those with LC. **(G)** Plasma proteomic measurement of IL-1𝛂 at 12 months post infection, separated by sex. * indicates unadjusted p<0.05 between LC versus recovered subjects. **(H)** Log2FC of pseudobulk *CD86* expression in CD14^+^ and CD16^+^ monocytes in sexes combined at 3 and 12 months post infection. Bars moving to the left of the dashed line indicate reduced expression in those with LC. **(I-J)** Plasma proteomic correlates of T cell exhaustion at **(I)** 3 months and **(J)** 12 months in those with persistent LC to 12 months compared to those with LC at 3 months but recovery by 12 months post infection, separated by sex. * indicates unadjusted p<0.05.

In females, different cell-cell communication patterns emerged. CD14^+^ monocytes in those with LC at 12 months showed IL1 signaling toward memory B cells, cDC2, and other CD14^+^ monocytes (Fig. 6C). These cells were also predicted to receive IL1β from CD16^+^ monocytes, as well as TGFβ1 signaling from NK cells, Mucosal Associated Invariant T (MAIT) cells, gamma delta T cells (ɣ𝛿T cells), and CD8+ T cells (Fig. 6D). These ligand-receptor interaction were predicted to increase expression of several genes which encode proteins involved in inflammation, such as chemokines *CXCL1* and *CXCL8*, cytokine *IL1β*, as well as *EREG* and *PTGS2*, in addition to pro-apoptotic *PMAIP1* (Fig. 6E).

Given the prevalence of *IL1β* signaling in females, we compared pseudobulk expression of this gene in monocytes in males and females, finding that both CD14^+^ and CD16^+^ monocytes in those with LC had increased *IL1β* expression in females; however, this was not the case in males (Fig. 6F). Consistent with these findings, plasma proteomic analysis demonstrated upregulation of IL-1𝛂 in females with persistent LC at 12 months, but not in males (Fig. 6G). When we evaluated single-cell proteomics, we found no differences in intracellular IL-1β between LC and recovered groups in either sex at 3 and 12 months (fig. S5), which may indicate that some of this protein had been secreted. However, in subjects with LC persisting to 12 months, intracellular IL-6 and IP-10 (aka CXCL10) were higher in CD14^+^ and CD16^+^ monocytes at 3 and 12 months in males, and IP-10 was higher in CD14^+^ monocytes at 3 months in females, indicating a pro-inflammatory state (fig. S5).

Given the role of monocytes in providing co-stimulatory signals to T cells, we next evaluated *CD86*, which is an important co-stimulatory molecule expressed on antigen-presenting cells (124) that interacts with the CD28 receptor on T cells to help induce proliferation and cytotoxicity (125). Interestingly, we also found reduced expression of *CD86* at 3 and 12 months after acute infection in CD14^+^ and CD16^+^ monocytes for those with LC in both sexes (Fig 6H). Plasma proteomics also supported reduced co-stimulatory function of monocytes in males, as LILRB4, which has been shown to induce immune tolerance in antigen-presenting cells and an immunosuppressive T cell phenotype (126, 127), was higher in male subjects with LC at 12 months (Fig. 6I-J). Additionally, several proteomic markers of T cell anergy/exhaustion, including IL-10, IL-10RB, and LAG3, were higher in those with persistent LC at 3 and 12 months, although this was only statistically significant for LAG3 in males (Fig. 6I-J). If sexes were considered together to increase power, the findings were similar for these markers (fig. S6). Altogether, monocytes of subjects with LC persistence demonstrated both sex-specific differences, such as prominent IL1 signaling and increased IL-1𝛂 protein secretion in females, as well as common findings in both sexes, such as reduced co-stimulatory *CD86* expression compared to recovered subjects.

### Widespread increases in *NFKB1* expression are associated with LC development and persistence

In addition to the role of *NF*-κ*B* in a pro-inflammatory monocyte phenotype in those with LC (Fig. 5B), sex-specific gene network analyses of a variety of cell types, such as CD16^+^ monocytes and naive B cells in males at 3 months, and CD8^+^ TEM in females at 12 months, revealed that *NFKB1* was upregulated in those with persistent LC (fig. S7A-C). Given this, we next analyzed transcriptional expression of NF-κB transcription factors across cell types and time points. Since we found that upregulation of *NF*-κ*B* constituents did not vary substantially by sex (fig. S8), the following analyses show the sexes combined.

*NFKB1*, which is a primary component of the canonical NF-κB signaling pathway and induces cell proliferation and immune response programs (*151*), was found to have higher expression during acute infection in those who will develop LC at 3 months, as well as higher expression at 3 and 12 months in those with LC persisting to 12 months (Fig 7D). A similar pattern was seen with *RELB*, an essential activator of transcription in the non-canonical NF-κB pathway (*152*), where upregulation was observed in all cell types other than CD8^+^ TEM and naive B cells during acute infection (Fig 7E). Lastly, *c-REL*, which is primarily expressed in B and T lymphocytes and has been implicated in multiple autoimmune conditions (*153*), also had higher expression in the acute samples of those who will develop LC and in those with persistent LC at 12 months (Fig 7F). Overall, widespread increases in several NF-κB transcription factor genes at multiple time points suggests a strong association between this pathway and LC pathogenesis.

**Fig. 7.**
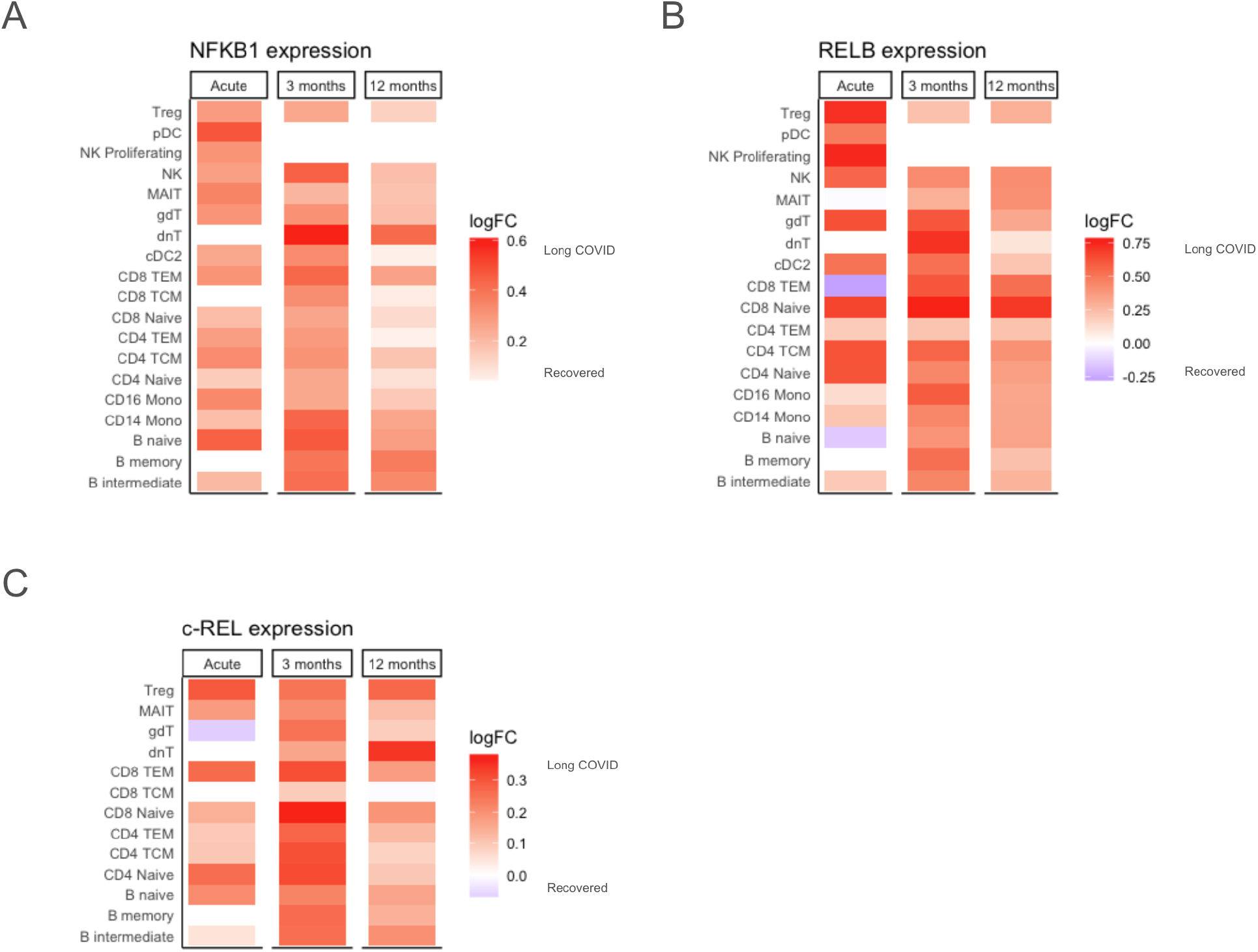
Increased expression of NF-κB transcription factors is associated with LC development and persistence. **(A-C)** Higher pseudobulk expression of **(A)** *NFkB1* and **(B)** *RELB* across many cell types is associated with LC development after acute infection and persistence for 12 months, with sexes combined. **(C)** Increased pseudobulk *c-REL* expression across T and B lymphocytes is associated with LC development after acute infection and persistence for 12 months, with sexes combined. Red tiles indicate higher expression (log2FC) in LC and blue tiles indicate lower expression.

### AP-1, *ETS1*, and T helper 2 immunity in LC

BTM analysis at 3 and 12 months after acute infection in subjects who had persistent LC at 12 months demonstrated increased expression of genes in the “signaling in T cells (I)” module (Fig. 5A-B). Inspection of this module revealed that four genes were implicated in a majority of these transcriptional differences: *FOS, FOSB, JUN,* and *JUNB* (Fig. S9). These genes are major components of the Activator Protein-1 (AP-1) transcription factor complex, which plays important roles in T cell activation, and transcriptional expression was increased in those with persistent LC across many cell types (fig. S9) (*154*). Several of these genes also demonstrated increased expression in those with LC in gene network analyses of various cell types (Fig. 7A-C).

Given these findings corresponding to altered T cell function in LC and known physical and functional associations between AP-1 and the *ETS1* transcription factor (*155*), we evaluated pseudobulk *ETS1* expression in lymphocyte subsets in which it is known to be expressed (*156*). During acute infection, *ETS1* expression was reduced in all B cell, T cell, and NK cell types in males who progressed to LC at 3 months post infection; however, it was increased in several lymphocyte subsets in females (Fig. 8A). Comparison of *ETS1* in those with LC versus those who recovered revealed lower expression at 3 months in those who would have persistent LC at 12 months, and it remained lower at 12 months in these individuals across most B and T cell subsets in both sexes (Fig. 8B).

**Fig. 8.**
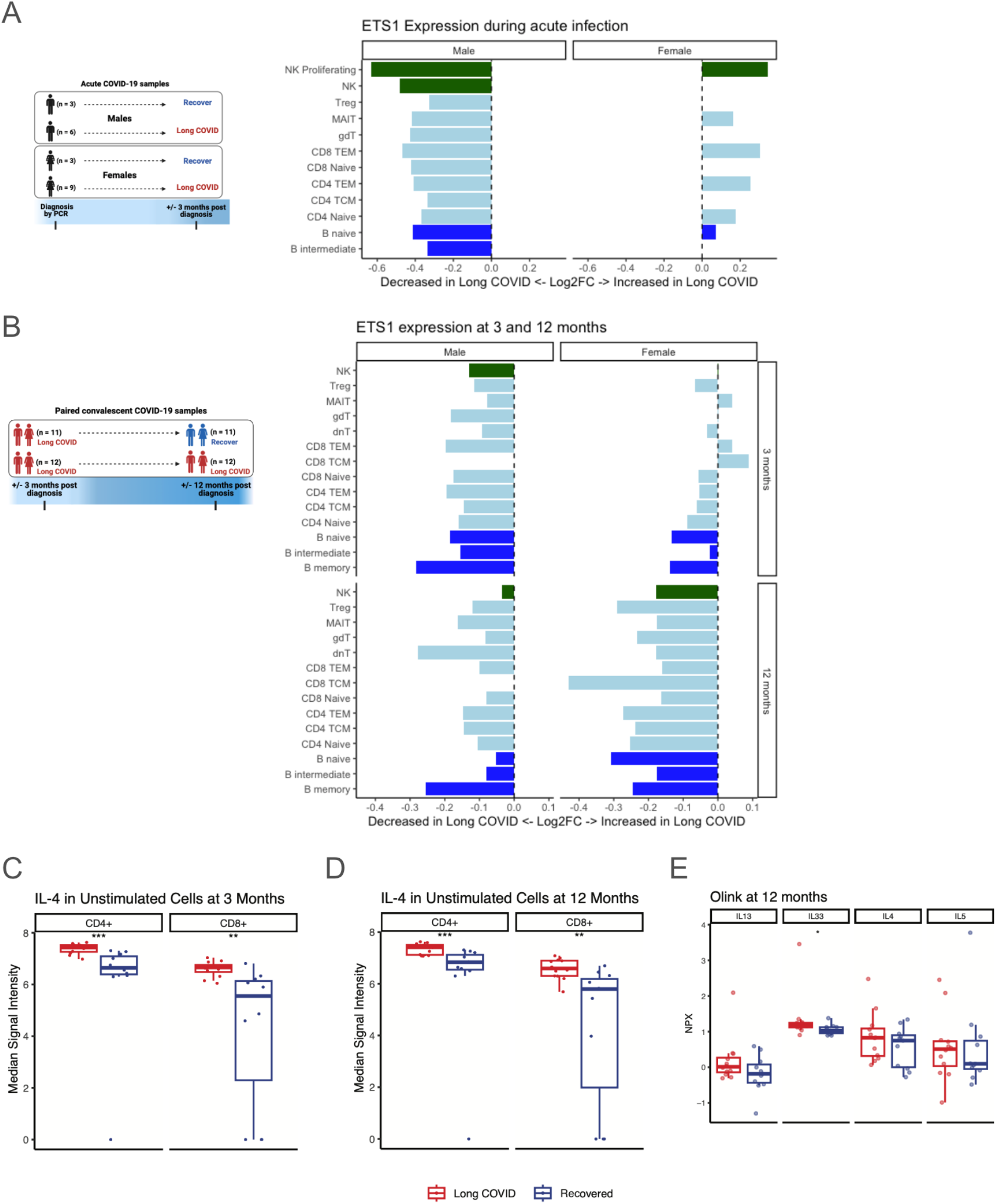
*ETS1* expression is reduced in LC, with a corresponding increase in Th2 polarization. **(A)** Log2FC of pseudobulk *ETS1* expression during acute infection, comparing those who develop LC at 3 months after acute infection versus those who recover by 3 months, separated by sex. Bar colors correspond to major cell types. **(B)** Log2FC of pseudobulk *ETS1* expression at 3 and 12 months post infection, comparing those with LC at 3 and 12 months versus those with LC at 3 months but recovery by 12 months, separated by sex. Bar colors correspond to major cell types. **(C-D)** Intracellular IL-4 as measured by CyTOF in CD4^+^ and CD8^+^ T cells at **(C)** 3 months and **(D)** 12 months post infection, comparing those with persistent LC at 3 and 12 months versus those with LC at 3 months but recovery by 12 months. **(E)** Plasma proteomic measurements of common Th2 immune cytokines IL-13, IL-33, IL-4, and IL-5, showing higher levels of all markers (only IL-33 meets unadjusted statistical significance) in those with persistent LC at 12 months versus those who recovered by 12 months. * indicates unadjusted p<0.05, ** indicates unadjusted p<0.01, and *** indicates unadjusted p<0.001.

As *ETS1* deficiency has previously been linked to increased T helper 2 (Th2) differentiation (*157*) we assessed proteomic correlates to the *ETS1* reduction seen in our transcriptomic data. Our CyTOF data revealed that IL-4, a major Th2 cytokine, was higher in CD4^+^ and CD8^+^ T cells at 3 and 12 months for subjects with persistent LC at 12 months (Fig. 8C-D). When separated by sex, this pattern persists across CD4^+^ naive, CD4^+^ memory, CD8^+^ naive, and CD8^+^ memory T cells (fig. S10A-B). Finally, plasma proteomics revealed higher IL-4, IL-5, IL-13, (though not reaching statistical significance) and higher IL-33 (p<0.05), all of which are known to induce Th2 signaling (*158*), in subjects with LC at 12 months (Fig. 8E). Together, these data suggest roles for increased AP-1 transcription factor, decreased *ETS1* transcription factor, and increased Th2 signaling in the pathogenesis of LC.

## DISCUSSION

This study uses multi-omic data to examine subjects during acute infection and 3 and 12 months after infection to discover abnormalities in innate and adaptive immunity of individuals with LC, many of which differ by sex. BTM, cell-communication, and gene network analyses uncovered cell-specific as well as more pervasive differences in those who developed LC, such as increased TGF-β signaling in males, NF-κB activation in both sexes, and reduced *ETS1* transcription factor expression in both sexes in numerous lymphocyte subsets in the months following acute infection. Some of these transcriptional differences were corroborated by corresponding changes in single-cell cytokine production and by plasma cytokine levels. Sex-specific immunologic differences were prominent across many cell types, indicating that mechanisms of LC may differ between sexes, potentially informing the search for tailored therapeutics.

Several genes with high specificity for biological sex were found to have expression differences that support roles in disease pathogenesis. In males, the *DDX3Y* gene was most reduced in those with LC at both 3 and 12 months, and the differences became most prominent in innate immune cells, such as CD56^bright^ NK cells, cDC2, and monocytes, at 12 months. *DDX3* genes have been shown to promote virus-induced interferon-β expression (*159*), therefore reduced *DDX3* expression may have implications for clearance of residual SARS-CoV-2 virus, suppression of other chronic viral infections, or response to new viral pathogens. In females, the *XIST* gene was noted to have higher expression during acute infection and at 3 and 12 months for those who developed LC. The RNA product of *XIST* does not encode a protein, but rather binds numerous other proteins and subsequently interacts with the X chromosome in females to silence gene transcription in one of the X chromosomes (*103*). This mechanism has been associated with multiple disease processes, and the Xist ribonucleoprotein complex has many sites of potential immunoreactivity that may contribute to multi-organ autoimmune pathology (*103*). An association between autoantibodies and LC has been suggested in some studies (*11, 55, 56*), but not in others (*12, 160*), necessitating further investigation that may benefit from considering sex-specific differences in titers, breadth, and association with LC symptoms.

In this cohort, there were notable perturbations in innate and early adaptive immunity during acute infection that correlated with development of LC. Proliferating NK cells in males showed bidirectional TGF-β signaling, which would be expected to reduce NK cell cytotoxicity, as previously observed in severe COVID-19 (*161*), which may limit the efficacy of the initial antiviral response. Increased *TGFβ1* expression was also found to be associated with LC development and persistence across many cell types, including naive and memory T cells, which may lead to reduced T helper 1 (Th1) differentiation (*162*) and impaired immunity against intracellular pathogens. Conversely, TGF-β signaling contributes to regulatory T cell differentiation, which may be important for avoiding self-reactivity (*163*). Interestingly, TGF-β has been previously suggested to play a role in myalgic encephalomyelitis/chronic fatigue syndrome (ME/CFS) (*164*) and was described to be increased in the plasma of males with LC in a pre-print study of a different LC patient cohort (*165*).

In convalescence, increased inflammatory signaling involving monocytes persisted to 12 months in those with LC, characterized by upregulation of major factors such as IL1 and NF-κB in transcriptomic data, IL-6 and CXCL10 in single-cell proteomics, and IL1 in plasma proteomics. Despite multiple proinflammatory markers, monocytes appeared to have dysfunctional interaction with T cells, as transcriptional analysis indicated reduced *CD86* co-stimulatory function, and plasma proteomics revealed multiple markers of T cell quiescence. Increased LAG3 was noted at 3 and 12 months in subjects with persistent LC, particularly in males. LAG3 is associated with suppressed/exhausted T cell phenotypes in the contexts of autoimmune disease, malignancy and infectious disease, and is under investigation as a target for novel cancer immunotherapies given its potential to reduce T cell cytotoxicity (149). In combination with our findings of reduced co-stimulatory CD86 expression in CD14^+^ monocytes, our data suggests a reduced capacity for appropriate innate and adaptive immune responses to existing infectious insults among subjects with persistent LC at 12 months, consistent with prior evidence showing reactivation of other viruses in those with LC (58). Potential impacts of the observed dysfunctional T cell signaling on host response to new pathogens deserves attention in future studies.

Single-cell transcriptomic analyses also revealed a broader pattern of reduced *ETS1* transcription factor expression in nearly all lymphocyte subsets to be associated with LC persistence to 12 months. *ETS1* deficiency in humans has also been characterized in the context of autoimmunity, where it was highly correlated with presence of autoimmune disease and disease severity (*157*). Additionally, *ETS1* deficiency was strongly associated with increased Th2 cell activity by biomarkers such as IL-4, IL-5, and IL-13 (*157*). *ETS1* deletion in mice also resulted in high IL-4 signaling activity, Th2 bias, and clinical features of autoimmunity (*157*). In our cohort, we found a significant increase in IL-4 protein expression across T cells in those with LC at 3 and 12 months, mirroring the reduced *ETS1* expression seen in these cells. While IL-4 signaling is classically associated with CD4^+^ T cell modulation, it also impacts CD8^+^ T cell function, where it induces pro-survival programs (*166*), but also reduces cytotoxicity in the setting of viral infection (*167*). Interestingly, IL-4 blockade has been shown to cause remission of autoimmune disease in *ETS1* deficient mice, and may offer a feasible intervention strategy in LC given the impressive efficacy and safety of anti-IL-4 receptor therapy in humans (*168*).

This study has several strengths. First, peripheral blood samples were collected during acute infection as well as convalescence, allowing for discovery of immune abnormalities during acute infection which correlate with LC symptom development, such as increased TGF-β signaling. Secondly, collection of samples at two time points after onset of LC revealed both features of LC and predictors of LC persistence, as demonstrated by *ETS1* and Th2 cytokine dysregulation. Third, the use of systems immunology approaches such as gene network analysis and cell-cell communication inference facilitated detection of central mediators of immune dysfunction that were cell- and sex-specific, as well as commonalities across cell types that may contribute to LC pathogenesis. Lastly, the use of single-cell and plasma proteomics allowed for confirmation that several pervasive transcriptional differences correspond to alterations in protein expression.

This study also has important limitations. The subjects enrolled in this study were first infected with SARS-CoV-2 early in the COVID-19 pandemic, prior to the widespread dissemination of mRNA vaccines which have greatly reduced the incidence of severe disease during acute infection. However, prevalence of LC has been relatively unchanged from 2022 to 2024 (*169*), suggesting that vaccination may not offer a comparable magnitude of protection against LC development as it does for severe disease during acute infection, and that ongoing evolution of the SARS-CoV-2 virus may not have significantly impacted post-acute sequelae. Additional studies of LC cohorts during subsequent SARS-CoV-2 variant epochs are needed to validate our findings. An additional limitation which applies to all LC studies is the lack of a consistent LC definition or objective measures that can be used for diagnosis. Since we prioritized performing transcriptomic analyses of multiple time points and analyzing the data by sex, this study was limited by small sample size. Additional studies with larger cohorts will be necessary to validate these findings. As this study was conducted on one patient cohort from a single institution, additional studies to validate these findings in multiple cohorts would be required prior to applying such findings to the clinical setting.

In summary, this study utilizes multiple modalities to uncover patterns of immune dysregulation that correlate with LC development, persistence, and resolution. Our discovery of sex-specific differences in immune response suggests that future mechanistic and therapeutic studies should consider stratification by sex, as supported by disproportionate impacts of severe acute disease in males (*80–86*) and LC in females (*17, 87–94*). We have also found several potential targets for intervention, most notably downregulation of *ETS1* transcription factor and increased expression of Th2 cytokines in those with LC. Lastly, persistent transcriptional aberrancies in T cell stimulation and T cell receptor signaling may have implications for infectious disease and immunity, an area which requires attention in future studies.

## MATERIALS AND METHODS

### Clinical cohort, patient samples, and study design

This study utilized samples collected from 45 participants enrolled in the prospective, observational patient cohort called “Infection Recovery in SARS-CoV-2” (IRIS). Adult patients were enrolled in this cohort approximately 3 months after symptomatic, polymerase chain reaction (PCR)-confirmed SARS-CoV-2 infection between August, 2020 - December, 2020 at Stanford University (*170*). Protocols were approved by the Stanford Institutional Review Board (IRB 57036), and participants provided written informed consent. Exclusion criteria for the overall IRIS cohort included pregnancy or active drug or alcohol abuse. For the participants in this study, those with immunocompromising conditions (e.g. malignancy, HIV, and solid organ transplant), on immunomodulatory medications (e.g. prednisone, mycophenolate mofetil, tacrolimus, and ibrutinib), or with medical conditions that may have accounted for their symptoms (e.g. cystic fibrosis and fibromyalgia) were also excluded. Participants were included in this study based on the availability of longitudinal blood samples and the prioritization of those with the clearest clinical phenotypes (e.g. asymptomatic throughout the entire convalescent period assessed for the recovered participants).

Study data were collected and managed using REDCap electronic data capture tools hosted at Stanford University (*171, 172*). PBMCs and plasma were collected at approximately 3 months and 12 months post infection. Clinical features including medical histories, physical exams, vital signs, and self-reported patient symptoms (both checklist survey and free text response) were evaluated at approximately 3, 6 and 12 months post SARS-CoV-2 infection.

Of the 45 participants in this study, 21 participants with banked PBMCs and plasma during acute SARS-CoV-2 infection were profiled in this work. Twelve of these patients had been previously enrolled in a phase 2, randomized, single-blind, placebo-controlled trial of single subcutaneous dose of Peginterferon Lambda-1a as potential treatment for outpatients with mild to moderate COVID-19 (NCT04331899) at Stanford University (*173*). Additionally, 9 of these patients had been previously enrolled in a phase 2, randomized, double-blind, placebo-controlled trial of 10-day course of oral Favipiravir as potential treatment for outpatient mild or asymptomatic COVID-19 (NCT04346628) at Stanford University (*174*). Notably, neither study found a difference in duration of SARS-CoV-2 viral shedding or in symptom duration between the treatment arm and the placebo arm (*173, 174*).

### Mass cytometry staining

PBMC samples were processed in a randomized fashion utilizing 20-plex barcoding. A consistent plate control sample was included for every CyTOF batch. A total of 26 batches were acquired. PBMC samples were individually thawed and added to pre-warmed media (RPMI 1640 (Thermo Scientific 21870092) supplemented with 10% fetal bovine serum (Corning Ref #35-016-CV), 2mM L-Glutamine (Thermo Scientific 25030081), and 1x Penicillin/Streptomycin/Amphotericin B (Gibco Cat #15240062 or Cytiva HyClone SV30079.01)) plus 20µL of benzonase (Millipore #70664) per 10mL total volume. Cells were washed, counted, and plated in 96-well round bottom plates (Thermo Scientific Cat #163320) at 1.5 million cells/200µL media per well such that each PBMC sample was plated into two wells (for an “unstimulated” and “stimulated” condition per sample). Cells were incubated overnight at 37°C, 5% CO2.

The following morning, cells in the wells designated for “stimulation” were resuspended in a combination of Resiquimod (R848) (4µg/mL; Invivogen Cat# tlrl-r848) and Polyinosinic:polycytidylic acid (Poly IC) (12.5µg/mL; Sigma P9582) for a total of 5 hours in the 37°C, 5% CO2 incubator, with the addition of phorbol 12-myristate 13-acetate and ionomycin (PMA/I) (0.0625x; eBioscience REF 00-4970-93) for the last two hours of stimulation. The cells that were designated to be “unstimulated” were mock-treated with media. A combination of 1x Brefeldin A (eBioscience #00-4506-51), 1x Monensin (eBioscience 00-4505-51), and 0.8µL CD107a-APC (BioLegend #328620) per 200µL of cells were added for the last 1.5 hours of the total 5 hour incubation for all cells. This stimulation protocol was previously optimized to induce activation and cytokine production of major PBMC cell subsets, including T cells, B cells, myeloid cells, and NK cells, while also minimizing cell loss.

After stimulation or mock incubation, cells were washed with CyFACS buffer (PBS, 0.1% BSA, 2nM EDTA, 0.05% sodium azide), barcoded in 20-plex format for 30 minutes at room temperature, thoroughly washed in CyFACS buffer, resuspended in PBS, and pooled into a round-bottom polypropylene tube (Falcon REF 352063). Pooled samples were then stained with PdCl viability reagent (final concentration of 500nM in PBS) for 5 minutes at room temperature and subsequently quenched with FBS. Following two CyFACS washes, cells were incubated with Human TruStain FcX (BioLegend 422302) at 5µL per million cells at room temperature for 10 minutes per manufacturer instructions. Next, cells were stained in 250µL of surface antibody cocktail (Table S5) for 30 minutes at 4°C. Following two CyFACS washes, cells were fixed in freshly diluted 2% paraformaldehyde (PFA) in PBS (16% PFA purchased from EMS Cat# 15710) for 20 minutes at room temperature in the dark. Cells were subsequently washed once in CyFACS, then twice in freshly prepared 1x eBioscience Permeabilization Buffer (Invitrogen Cat #00-8333-56) with spins at 800g for 5 minutes at 4°C. Cells were then stained with 250µl of intracellular antibody cocktail (Table S6) for 45 minutes at 4°C in the dark. Following 1 wash with Permeabilization Buffer and 2 washes with CyFACS, cells were resuspended in 35nM Cell ID Intercalator (191Ir, 193Ir) in 2% PFA and incubated overnight at 4°C in the dark. The following morning, cells were washed once with CyFACS and twice with Milli-Q® water. Cells were subsequently resuspended in Milli-Q® water and 0.1x EQ Four Element Calibration Beads (Fluidigm/Standard BioTools, Cat # 201078) and filtered through a 35µm nylon mesh into polystyrene tubes (BD Falcon #352235) for sample acquisition. Data were acquired on a Helios mass cytometer.

### Mass cytometry data preprocessing and data analysis

Flow Cytometry Standard (FCS) files were normalized and debarcoded using the Premessa package, as previously described (*175, 176*). FlowJo version 10.8.1 was used to manually gate out EQ beads and to gate on intact, single, live, CD45+ cells, which were subsequently exported as FCS files for further analysis using the open software R. Since no major batch effects were observed, batch normalization was not performed.

Mass cytometry data was arcsinh transformed with a cofactor of 0.001 prior to clustering. This transformation was chosen to optimize multimodality across markers (fig. S11), and thus facilitate cell clustering. Coarse cell clusters (monocytes/dendritic cells, natural killer cells, CD4+ T cells, CD8+ T cells, and B cells) were determined using mini-batch K-means clustering in R on a subset of 1,032,000 cells. Only CD3, CD4, CD8a, CD11c, CD14, CD16, CD19, CD20, CD56, and HLA-DR were utilized for coarse clustering. FlowSOM and ConsensusClusterPlus were utilized for clustering into fine cell types within the aforementioned coarse clusters using lineage markers (*177*). Random forest classification was used to apply these clustering rules to the entire dataset (*178*). Median signal intensity of functional markers with biological relevance to specific cell types mentioned in the text were calculated for each subject under unstimulated conditions.

### Olink® Proteomics

Plasma samples collected from IRIS study participants during acute SARS-CoV-2 infection, 3 months post infection, and 12 months post infection were analyzed using the Olink® Target 96 Inflammation panel and Olink® Target 96 Immune Response panel. Briefly, 80µl of plasma per sample was plated in a 96-well PCR plate (Sigma Aldrich #EP951020401), frozen, and shipped on dry ice to the Olink U.S. Service Lab in Waltham, MA where the assays were performed.

Proteomic data was obtained from the Olink biomarker platform. Since plasma samples from acutely infected patients were collected in two different tube types, PCA was performed to assess for batch effects between plasma samples from the Peginterferon Lambda-1a study (collected in sodium citrate tubes) and the Favipiravir study (collected in heparinized tubes). IRIS study plasma samples were collected in heparinized tubes. Given the presence of batch effect, the olink_normalization() function in the OlinkAnalayze R package was used for batch correction (*179*). Batch-corrected proteomic data was then utilized as confirmatory evidence for findings from scRNA-seq and mass cytometry data.

### Single-cell RNA sequencing

PBMC samples (up to 2 million cells per sample) were fixed using the Evercode Cell Fixation v2 (ECF2001) from Parse Biosciences per the manufacturer’s protocol, placed in a “Mr. Frosty,” and stored at -80°C for up to 6 months. After all samples were fixed, samples were processed using the Parse Biosciences Evercode WT Mega v2 (ECW02050) per the manufacturer’s protocol. A total of 1 million cells were processed, which included approximately 10,500 cells per sample from samples during acute SARS-CoV-infection as well as convalescent samples from 3 months and 12 months post infection. Briefly, the protocol included: 3 rounds of split-pool barcoding in 96-well plates, followed by cell lysis and sublibrary generation, amplification of barcoded cDNA, fragmentation, and size selection. Using the Agilent TapeStation 4150 to evaluate cDNA quality and concentration, one of the 16 sublibraries generated was deemed to be of suboptimal quality with lower cDNA concentration, so the remaining fifteen sublibraries were ultimately sequenced on a NovaSeq S4 instrument at approximately 30,000 reads per cell (Illumina; Chan Zuckerberg Biohub).

### Single-cell RNA sequencing data alignment, quality control, and reference mapping

The resulting sequencing data were processed using the Parse Biosciences Pipeline v1.0.3 per manufacturer guidelines. A custom genome containing the human genome (HG38) and the annotated Wuhan-Hu-1 SARS-CoV-2 reference sequence (NC_045512.2) was used as the reference genome for alignment. Further analysis was performed using the R package Seurat (*100*). To remove cell multiplets and low-quality or dying cells, cells with >7500 unique genes or >30,000 total RNA molecules or <100 genes expressed or >20% mitochondrial genes were removed from further analysis. Genes with expression in <100 cells, mitochondrial genes, ribosomal genes, and genes without annotation in Ensembl were also removed to facilitate interpretability.

The data was scaled and transformed using the SCTransform() function, and linear dimensional reduction was performed with principal component analysis (PCA). Using the first 50 principal components, UMAP was performed, a K-nearest neighbors graph was constructed using the FindNeighbors() function, and the cells were clustered using the FindClusters() function. One resulting cluster was identified as having about 22% mitochondrial or ribosomal RNA genes, so this cluster was removed from subsequent analyses. All remaining clusters had <5% mitochondrial or ribosomal RNA genes and were retained for downstream analyses.

For cell type annotation, we utilized Seurat v4 PBMC multimodal reference mapping, as previously described (*100*). Briefly, anchors were found between the reference and the dataset using the FindTransferAnchors() function with a precomputed supervised PCA transformation and 50 principal components. Using the MapQuery() function, cell type labels from “celltype.l2” were transferred from the reference to the dataset, and the dataset was projected onto the reference UMAP structure. To validate cell type annotation, canonical marker gene expression was evaluated (fig. S12) (*180*). Notably, “platelets” expressed genes expected for CD14^+^ monocytes in addition to genes expected for platelets, so “platelets” were removed from downstream analyses. Marker genes for innate lymphoid cells (“ILCs”) were not robustly expressed, so “ILCs” were also removed from downstream analyses.

### Single-cell RNA sequencing analysis of differentially expressed genes

RNA data was transformed by estimating the size factors using the computeLibraryFactors() and then logNormCounts() functions from the scater package (*181*). The modelGeneVar() function was used to model the mean-variance relationship of genes, and the top 1000 most variable genes for each cell type, sex, and time point relative to acute infection were selected for differential expression analysis using the getTopHVGs() function (*182*). Differentially expressed genes (DEGs) between conditions (i.e. LC or recovered) were determined using the Seurat FindMarkers() function and the MAST method (*183*), specifying the sample as a latent variable. Pseudobulk expression was assessed using edgeR, as implemented by the pbDS() function in the muscat R package (*184*).

Gene network analyses were conducted using the R package BioNet (*185*). To construct the gene network, the top genes (up to 400) by absolute log2 fold change for the cell type of interest were read into STRING version 12.0 (*186*) to produce a gene interaction network. Only genes with absolute log2 fold change >0.25 were included. This gene interaction data was then utilized as the interaction network for discovery of gene modules with BioNet, using unadjusted p values from the MAST DEG analysis as input. Modules were created with adjustment of false-discovery rates (all <0.05) to reduce network complexity and facilitate visualization of genes central to the module. The ggnetwork R package (*187*) was used to customize module visualization, and gene nodes colored by log2 fold change.

### Blood Transcription Module analysis

Immunologic pathway analysis was conducted using Blood Transcription Modules (BTMs), as previously described (*101*). Using output from the MAST DEG analysis, we included the top 500 genes by absolute log2 fold change which are present in annotated BTMs. Only modules with at least 10% of genes in the list of 500 were plotted, and for some figures we further reduced to only the top 50 BTMs to simplify visualization (as described in figure captions). BTM analyses were then used to select cell types for further analysis, and we prioritized the cell types which showed the greatest number of BTMs which differed between LC and recovered subjects.

### Cell-cell communication analysis

Cell-cell communication analyses were performed with MultiNicheNet (*109*). Selected parameters for our analysis included: minimum cell count per cell type and condition of 10; an absolute log2FC threshold of 0.5; an unadjusted p value threshold of 0.05; a non-zero expression value threshold of 0.05; and seeking the top 250 target genes predictions. Gene prioritization was performed with a differential expression weight of 1 for both ligand and receptor, a scaled ligand activity weight of 2, expression specificity weights of 2 for both ligand and receptor, a fraction of cells expressing ligands and receptors weight of 1, and a relative abundance weight of 0 for both sender and receiver cells. Resulting ligand-receptor pairs were manually curated using literature searches. Pairs that either did not have experimental evidence of interaction or the interaction was known to occur within the same cell (rather than as ligand-receptor pair on communicating cells) were not included in resulting figures (Table S7).

### Statistical analysis

Comparison of means was performed with the Wilcoxon rank sum test. All data analysis was performed in using R version 4.3.1 on a macOS when possible. For analyses which required additional memory or computer processing units, R version 4.2.0 was utilized on a Linux cluster.

## List of Supplementary Materials

Materials and Methods

Fig. S1 to S12

Tables S1 to S7

## Acknowledgments

We gratefully acknowledge all the participants in the “Infection Recovery in SARS-CoV-2” (IRIS) cohort and in the Peginterferon Lambda-1a and Favipiravir clinical trials. We are thankful to all the trial coordinators, lab personnel and staff who contributed to the success of these trials and cohort. We thank the Stanford ChEM-H and Stanford Innovative Medicines Accelerator Clinical Research Coordinator. We are grateful to Mark Davis and his laboratory for the use of their CyTOF machine. We thank Michael Leipold for CyTOF training and expertise. We thank Arjun Rustagi for his guidance with Parse Biosciences scRNAseq library preparation and data pre-processing. We appreciate the assistance of Norma Neff and Amanda Seng of the Chan Zuckerberg Biohub with sequencing. We also thank Kalani Ratnasiri for helpful discussions regarding scRNA-seq analysis and guidance in implementing BTM analysis. Figure schematics were created with BioRender.com.

## Funding

Stanford Innovative Medicines Accelerator IMA-1035 (CAB, AS, PG, HB, SPH).

National Institutes of Health grants OT2HL161847 (US, CAB), T32AI007502 (REH), 5T32AI007290-37 (LC), and U01 AI150741-01S1 (PJ).

Chan Zuckerberg Biohub Investigator Award (CAB).

Anonymous donors to Stanford University provided funding support for the Favipiravir and Peginterferon Lambda-1a clinical trials.

Peginterferon Lambda-1a was provided by Eiger BioPharmaceuticals.

National Institutes of Health, National Center for Research Resources, and the Stanford Clinical and Translational Science Award UL1TR001085 (REDCap platform services at Stanford).

## Author contributions

Conceptualization: REH, SMP, SPH, AS, PG, HB, CAB Methodology: SMP, REH, SPH, CAB

Investigation: REH, LC, MAS, KP Formal analysis: SMP, REH, SPH, CAB

Resources: PG, AS, HB, MH, US, KBJ, PJ, YM, MR, CAB

Visualization: SMP, REH, SPH, CAB

Funding acquisition: CAB, AS, PG, HB, US, PJ

Project administration: MAS, KP, MR Supervision: AS, CAB, SPH

Writing – original draft: REH, SMP, CAB

Writing – review & editing: REH, SMP, LC, MAS, KP, MR, PG, AS, HB, MH, US, KBJ, PJ, YM, SPH, CAB

## Data and materials availability

Mass cytometry FCS files with de-identified metadata supporting this publication will be made available in FlowRepository (http://flowrepository.org/). Data from scRNA-seq will be deposited with the Gene Expression Omnibus. Olink data with de-identified metadata will be made available. All original code for analysis and visualization will be available in a Github repository.

## References and Notes

1. WHO Coronavirus (COVID-19) Dashboard (available at https://covid19.who.int).

2. Coronavirus disease (COVID-19): Post COVID-19 condition (available at https://www.who.int/news-room/questions-and-answers/item/coronavirus-disease-(covid-19)-post-covid-19-condition).

3. A. S. for P. Affairs (ASPA), About Long COVID (2023) (available at https://www.covid.gov/be-informed/longcovid/about).

4. CDC, Healthcare WorkersCent. Dis. Control Prev. (2020) (available at https://www.cdc.gov/coronavirus/2019-ncov/hcp/clinical-care/post-covid-conditions.html).

5. H. E. Davis, L. McCorkell, J. M. Vogel, E. J. Topol, Long COVID: major findings, mechanisms and recommendations. Nat. Rev. Microbiol. 21, 133–146 (2023).

6. M. Merad, C. A. Blish, F. Sallusto, A. Iwasaki, The immunology and immunopathology of COVID-19. Science 375, 1122–1127 (2022).

7. J. T. Reese, H. Blau, E. Casiraghi, T. Bergquist, J. J. Loomba, T. J. Callahan, B. Laraway, C. Antonescu, B. Coleman, M. Gargano, K. J. Wilkins, L. Cappelletti, T. Fontana, N. Ammar, B. Antony, T. M. Murali, J. H. Caufield, G. Karlebach, J. A. McMurry, A. Williams, R. Moffitt, J. Banerjee, A. E. Solomonides, H. Davis, K. Kostka, G. Valentini, D. Sahner, C. G. Chute, C. Madlock-Brown, M. A. Haendel, P. N. Robinson, N3C Consortium, RECOVER Consortium, Generalisable long COVID subtypes: findings from the NIH N3C and RECOVER programmes. EBioMedicine 87, 104413 (2023).

8. T. Thaweethai, S. E. Jolley, E. W. Karlson, E. B. Levitan, B. Levy, G. A. McComsey, L. McCorkell, G. N. Nadkarni, S. Parthasarathy, U. Singh, T. A. Walker, C. A. Selvaggi, D. J. Shinnick, C. C. M. Schulte, R. Atchley-Challenner, G. A. Alba, R. Alicic, N. Altman, K. Anglin, U. Argueta, H. Ashktorab, G. Baslet, I. V. Bassett, L. Bateman, B. Bedi, S. Bhattacharyya, M.-A. Bind, A. L. Blomkalns, H. Bonilla, P. A. Bush, M. Castro, J. Chan, A. W. Charney, P. Chen, L. B. Chibnik, H. Y. Chu, R. G. Clifton, M. M. Costantine, S. K. Cribbs, S. I. Davila Nieves, S. G. Deeks, A. Duven, I. F. Emery, N. Erdmann, K. M. Erlandson, K. C. Ernst, R. Farah-Abraham, C. E. Farner, E. M. Feuerriegel, J. Fleurimont, V. Fonseca, N. Franko, V. Gainer, J. C. Gander, E. M. Gardner, L. N. Geng, K. S. Gibson, M. Go, J. D. Goldman, H. Grebe, F. L. Greenway, M. Habli, J. Hafner, J. E. Han, K. A. Hanson, J. Heath, C. Hernandez, R. Hess, S. L. Hodder, M. K. Hoffman, S. E. Hoover, B. Huang, B. L. Hughes, P. Jagannathan, J. John, M. R. Jordan, S. D. Katz, E. S. Kaufman, J. D. Kelly, S. W. Kelly, M. M. Kemp, J. P. Kirwan, J. D. Klein, K. S. Knox, J. A. Krishnan, A. Kumar, A. O. Laiyemo, A. A. Lambert, M. Lanca, J. K. Lee-Iannotti, B. P. Logarbo, M. T. Longo, C. A. Luciano, K. Lutrick, J. H. Maley, J. G. Marathe, V. Marconi, G. D. Marshall, C. F. Martin, Y. Matusov, A. Mehari, H. Mendez-Figueroa, R. Mermelstein, T. D. Metz, R. Morse, J. Mosier, C. Mouchati, J. Mullington, S. N. Murphy, R. B. Neuman, J. Z. Nikolich, I. Ofotokun, E. Ojemakinde, A. Palatnik, K. Palomares, T. Parimon, S. Parry, J. E. Patterson, T. F. Patterson, R. E. Patzer, M. J. Peluso, P. Pemu, C. M. Pettker, B. A. Plunkett, K. Pogreba-Brown, A. Poppas, J. G. Quigley, U. Reddy, R. Reece, H. Reeder, W. B. Reeves, E. M. Reiman, F. Rischard, J. Rosand, D. J. Rouse, A. Ruff, G. Saade, G. J. Sandoval, S. M. Schlater, F. Shepherd, Z. A. Sherif, H. Simhan, N. G. Singer, D. W. Skupski, A. Sowles, J. A. Sparks, F. I. Sukhera, B. S. Taylor, L. Teunis, R. J. Thomas, J. M. Thorp, P. Thuluvath, A. Ticotsky, A. T. Tita, K. R. Tuttle, A. E. Urdaneta, D. Valdivieso, T. M. VanWagoner, A. Vasey, M. Verduzco-Gutierrez, Z. S. Wallace, H. D. Ward, D. E. Warren, S. J. Weiner, S. Welch, S. W. Whiteheart, Z. Wiley, J. P. Wisnivesky, L. M. Yee, S. Zisis, L. I. Horwitz, A. S. Foulkes, RECOVER Consortium, Development of a Definition of Postacute Sequelae of SARS-CoV-2 Infection. JAMA 329, 1934–1946 (2023).

9. A. Ozonoff, N. D. Jayavelu, S. Liu, E. Melamed, C. E. Milliren, J. Qi, L. N. Geng, G. A. McComsey, C. B. Cairns, L. R. Baden, J. Schaenman, A. C. Shaw, H. Samaha, V. Seyfert-Margolis, F. Krammer, L. B. Rosen, H. Steen, C. Syphurs, R. Dandekar, C. P. Shannon, R. P. Sekaly, L. I. R. Ehrlich, D. B. Corry, F. Kheradmand, M. A. Atkinson, S. C. Brakenridge, N. I. Higuita, J. P. Metcalf, C. L. Hough, W. B. Messer, B. Pulendran, K. C. Nadeau, M. M. Davis, A. F. Sesma, V. Simon, H. van Bakel, S. Kim-Schulze, D. A. Hafler, O. Levy, M. Kraft, C. Bime, E. K. Haddad, C. S. Calfee, D. J. Erle, C. R. Langelier, W. Eckalbar, S. E. Bosinger, B. Peters, S. H. Kleinstein, E. F. Reed, A. D. Augustine, J. Diray-Arce, H. T. Maecker, M. C. Altman, R. R. Montgomery, P. M. Becker, N. Rouphael, Features of acute COVID-19 associated with post-acute sequelae of SARS-CoV-2 phenotypes: results from the IMPACC study. Nat. Commun. 15, 216 (2024).

10. G. Kenny, K. McCann, C. O’Brien, S. Savinelli, W. Tinago, O. Yousif, J. S. Lambert, C. O’Broin, E. R. Feeney, E. De Barra, P. Doran, P. W. G. Mallon, All-Ireland Infectious Diseases (AIID) Cohort Study Group, Identification of Distinct Long COVID Clinical Phenotypes Through Cluster Analysis of Self-Reported Symptoms. Open Forum Infect. Dis. 9, ofac060 (2022).

11. Y. Su, D. Yuan, D. G. Chen, R. H. Ng, K. Wang, J. Choi, S. Li, S. Hong, R. Zhang, J. Xie, S. Kornilov, K. Scherler, A. J. Pavlovitch-Bedzyk, S. Dong, C. Lausted, I. Lee, S. Fallen, C. L. Dai, P. Baloni, B. Smith, V. R. Duvvuri, K. G. Anderson, J. Li, F. Yang, C. J. Duncombe, D. J. McCulloch, C. Rostomily, P. Troisch, J. Zhou, S. Mackay, Q. DeGottardi, D. H. May, R. Taniguchi, R. M. Gittelman, M. Klinger, T. M. Snyder, R. Roper, G. Wojciechowska, K. Murray, R. Edmark, S. Evans, L. Jones, Y. Zhou, L. Rowen, R. Liu, W. Chour, H. A. Algren, W. R. Berrington, J. A. Wallick, R. A. Cochran, M. E. Micikas, ISB-Swedish COVID-19 Biobanking Unit, T. Wrin, C. J. Petropoulos, H. R. Cole, T. D. Fischer, W. Wei, D. S. B. Hoon, N. D. Price, N. Subramanian, J. A. Hill, J. Hadlock, A. T. Magis, A. Ribas, L. L. Lanier, S. D. Boyd, J. A. Bluestone, H. Chu, L. Hood, R. Gottardo, P. D. Greenberg, M. M. Davis, J. D. Goldman, J. R. Heath, Multiple early factors anticipate post-acute COVID-19 sequelae. Cell 185, 881–895.e20 (2022).

12. J. Klein, J. Wood, J. Jaycox, R. M. Dhodapkar, P. Lu, J. R. Gehlhausen, A. Tabachnikova, K. Greene, L. Tabacof, A. A. Malik, V. Silva Monteiro, J. Silva, K. Kamath, M. Zhang, A. Dhal, I. M. Ott, G. Valle, M. Peña-Hernandez, T. Mao, B. Bhattacharjee, T. Takahashi, C. Lucas, E. Song, D. Mccarthy, E. Breyman, J. Tosto-Mancuso, Y. Dai, E. Perotti, K. Akduman, T. J. Tzeng, Xu, A. C. Geraghty, M. Monje, I. Yildirim, J. Shon, R. Medzhitov, D. Lutchmansingh, J. D. Possick, N. Kaminski, S. B. Omer, H. M. Krumholz, L. Guan, C. S. Dela Cruz, D. van Dijk, A. Ring, D. Putrino, A. Iwasaki, Distinguishing features of Long COVID identified through immune profiling. Nature (2023), doi:10.1038/s41586-023-06651-y.

13. K. Yin, M. J. Peluso, R. Thomas, M.-G. Shin, J. Neidleman, X. Luo, R. Hoh, K. Anglin, B. Huang, U. Argueta, M. Lopez, D. Valdivieso, K. Asare, R. Ibrahim, L. Ständker, S. Lu, S. A. Goldberg, S. A. Lee, K. L. Lynch, J. D. Kelly, J. N. Martin, J. Münch, S. G. Deeks, T. J. Henrich, N. R. Roan, Long COVID manifests with T cell dysregulation, inflammation, and an uncoordinated adaptive immune response to SARS-CoV-2. BioRxiv Prepr. Serv. Biol. , 2023.02.09.527892 (2023).

14. C. Phetsouphanh, D. R. Darley, D. B. Wilson, A. Howe, C. M. L. Munier, S. K. Patel, J. A. Juno, L. M. Burrell, S. J. Kent, G. J. Dore, A. D. Kelleher, G. V. Matthews, Immunological dysfunction persists for 8 months following initial mild-to-moderate SARS-CoV-2 infection. Nat. Immunol. 23, 210–216 (2022).

15. M. J. Peluso, A. N. Deitchman, L. Torres, N. S. Iyer, S. E. Munter, C. C. Nixon, J. Donatelli, C. Thanh, S. Takahashi, J. Hakim, K. Turcios, O. Janson, R. Hoh, V. Tai, Y. Hernandez, E. A. Fehrman, M. A. Spinelli, M. Gandhi, L. Trinh, T. Wrin, C. J. Petropoulos, F. T. Aweeka, I. Rodriguez-Barraquer, J. D. Kelly, J. N. Martin, S. G. Deeks, B. Greenhouse, R. L. Rutishauser, T. J. Henrich, Long-term SARS-CoV-2-specific immune and inflammatory responses in individuals recovering from COVID-19 with and without post-acute symptoms. Cell Rep. 36, 109518 (2021).

16. M. J. Peluso, S. Lu, A. F. Tang, M. S. Durstenfeld, H.-E. Ho, S. A. Goldberg, C. A. Forman, E. Munter, R. Hoh, V. Tai, A. Chenna, B. C. Yee, J. W. Winslow, C. J. Petropoulos, B. Greenhouse, P. W. Hunt, P. Y. Hsue, J. N. Martin, J. Daniel Kelly, D. V. Glidden, S. G. Deeks, J. Henrich, Markers of Immune Activation and Inflammation in Individuals With Postacute Sequelae of Severe Acute Respiratory Syndrome Coronavirus 2 Infection. J. Infect. Dis. 224, 1839–1848 (2021).

17. PHOSP-COVID Collaborative Group, Clinical characteristics with inflammation profiling of long COVID and association with 1-year recovery following hospitalisation in the UK: a prospective observational study. Lancet Respir. Med. 10, 761–775 (2022).

18. K. M. Littlefield, R. O. Watson, J. M. Schneider, C. P. Neff, E. Yamada, M. Zhang, T. B. Campbell, M. T. Falta, S. E. Jolley, A. P. Fontenot, B. E. Palmer, SARS-CoV-2-specific T cells associate with inflammation and reduced lung function in pulmonary post-acute sequalae of SARS-CoV-2. PLOS Pathog. 18, e1010359 (2022).

19. C. Schultheiß, E. Willscher, L. Paschold, C. Gottschick, B. Klee, S.-S. Henkes, L. Bosurgi, J. Dutzmann, D. Sedding, T. Frese, M. Girndt, J. I. Höll, M. Gekle, R. Mikolajczyk, M. Binder, The IL-1β, IL-6, and TNF cytokine triad is associated with post-acute sequelae of COVID-19. Cell Rep. Med. 3, 100663 (2022).

20. R. C. Thompson, N. W. Simons, L. Wilkins, E. Cheng, D. M. Del Valle, G. E. Hoffman, C. Cervia, B. Fennessy, K. Mouskas, N. J. Francoeur, J. S. Johnson, L. Lepow, J. Le Berichel, C. Chang, A. G. Beckmann, Y.-C. Wang, K. Nie, N. Zaki, K. Tuballes, V. Barcessat, M. A. Cedillo, D. Yuan, L. Huckins, P. Roussos, T. U. Marron, Mount Sinai COVID-19 Biobank Team, B. S. Glicksberg, G. Nadkarni, J. R. Heath, E. Gonzalez-Kozlova, O. Boyman, S. Kim-Schulze, R. Sebra, M. Merad, S. Gnjatic, E. E. Schadt, A. W. Charney, N. D. Beckmann, Molecular states during acute COVID-19 reveal distinct etiologies of long-term sequelae. Nat. Med. 29, 236–246 (2023).

21. A. Fernández-Castañeda, P. Lu, A. C. Geraghty, E. Song, M.-H. Lee, J. Wood, M. R. O’Dea, S. Dutton, K. Shamardani, K. Nwangwu, R. Mancusi, B. Yalçın, K. R. Taylor, L. Acosta-Alvarez, K. Malacon, M. B. Keough, L. Ni, P. J. Woo, D. Contreras-Esquivel, A. M. S. Toland, J. R. Gehlhausen, J. Klein, T. Takahashi, J. Silva, B. Israelow, C. Lucas, T. Mao, M. A. Peña-Hernández, A. Tabachnikova, R. J. Homer, L. Tabacof, J. Tosto-Mancuso, E. Breyman, A. Kontorovich, D. McCarthy, M. Quezado, H. Vogel, M. M. Hefti, D. P. Perl, S. Liddelow, R. Folkerth, D. Putrino, A. Nath, A. Iwasaki, M. Monje, Mild respiratory COVID can cause multi-lineage neural cell and myelin dysregulation. Cell 185, 2452–2468.e16 (2022).

22. N. A. Scott, L. Pearmain, S. B. Knight, O. Brand, D. J. Morgan, C. Jagger, S. Harbach, S. Khan, H. A. Shuwa, M. Franklin, V. Kästele, T. Williams, I. Prise, F. A. McClure, P. Hackney, L. Smith, M. Menon, J. E. Konkel, C. Lawless, J. Wilson, A. G. Mathioudakis, S. C. Stanel, A. Ustianowski, G. Lindergard, S. Brij, N. Diar Bakerly, P. Dark, C. Brightling, P. Rivera-Ortega, G. M. Lord, A. Horsley, CIRCO, K. Piper Hanley, T. Felton, A. Simpson, J. R. Grainger, T. Hussell, E. R. Mann, Monocyte migration profiles define disease severity in acute COVID-19 and unique features of long COVID. Eur. Respir. J. 61, 2202226 (2023).

23. B. Vijayakumar, K. Boustani, P. P. Ogger, A. Papadaki, J. Tonkin, C. M. Orton, P. Ghai, K. Suveizdyte, R. J. Hewitt, S. R. Desai, A. Devaraj, R. J. Snelgrove, P. L. Molyneaux, J. L. Garner, J. E. Peters, P. L. Shah, C. M. Lloyd, J. A. Harker, Immuno-proteomic profiling reveals aberrant immune cell regulation in the airways of individuals with ongoing post-COVID-19 respiratory disease. Immunity 55, 542–556.e5 (2022).

24. L. Visvabharathy, B. A. Hanson, Z. S. Orban, P. H. Lim, N. M. Palacio, M. Jimenez, J. R. Clark, E. L. Graham, E. M. Liotta, G. Tachas, P. Penaloza-MacMaster, I. J. Koralnik, Neuro-PASC is characterized by enhanced CD4+ and diminished CD8+ T cell responses to SARS-CoV-2 Nucleocapsid protein. Front. Immunol. 14, 1155770 (2023).

25. H. A. Shuwa, T. N. Shaw, S. B. Knight, K. Wemyss, F. A. McClure, L. Pearmain, I. Prise, C. Jagger, D. J. Morgan, S. Khan, O. Brand, E. R. Mann, A. Ustianowski, N. D. Bakerly, P. Dark, C. E. Brightling, S. Brij, CIRCO, T. Felton, A. Simpson, J. R. Grainger, T. Hussell, J. E. Konkel, M. Menon, Alterations in T and B cell function persist in convalescent COVID-19 patients. Med N. Y. N 2, 720–735.e4 (2021).

26. H. Ruffieux, A. L. Hanson, S. Lodge, N. G. Lawler, L. Whiley, N. Gray, T. H. Nolan, L. Bergamaschi, F. Mescia, L. Turner, A. de Sa, V. S. Pelly, Cambridge Institute of Therapeutic Immunology and Infectious Disease-National Institute of Health Research (CITIID-NIHR) BioResource COVID-19 Collaboration, P. Kotagiri, N. Kingston, J. R. Bradley, E. Holmes, J. Wist, J. K. Nicholson, P. A. Lyons, K. G. C. Smith, S. Richardson, G. R. Bantug, C. Hess, A patient-centric modeling framework captures recovery from SARS-CoV-2 infection. Nat. Immunol. 24, 349–358 (2023).

27. A. Talla, S. V. Vasaikar, G. L. Szeto, M. P. Lemos, J. L. Czartoski, H. MacMillan, Z. Moodie, K. W. Cohen, L. B. Fleming, Z. Thomson, L. Okada, L. A. Becker, E. M. Coffey, S. C. De Rosa, E. W. Newell, P. J. Skene, X. Li, T. F. Bumol, M. Juliana McElrath, T. R. Torgerson, Persistent serum protein signatures define an inflammatory subcategory of long COVID. Nat. Commun. 14, 3417 (2023).

28. F. R. Hopkins, M. Govender, C. Svanberg, J. Nordgren, H. Waller, Å. Nilsdotter-Augustinsson, A. J. Henningsson, M. Hagbom, J. Sjöwall, S. Nyström, M. Larsson, Major alterations to monocyte and dendritic cell subsets lasting more than 6 months after hospitalization for COVID-19. Front. Immunol. 13, 1082912 (2022).

29. G. Captur, J. C. Moon, C.-C. Topriceanu, G. Joy, L. Swadling, J. Hallqvist, I. Doykov, N. Patel, J. Spiewak, T. Baldwin, M. Hamblin, K. Menacho, M. Fontana, T. A. Treibel, C. Manisty, B. O’Brien, J. M. Gibbons, C. Pade, T. Brooks, D. M. Altmann, R. J. Boyton, Á. McKnight, M. K. Maini, M. Noursadeghi, K. Mills, W. E. Heywood, UK COVIDsortium Investigators, Plasma proteomic signature predicts who will get persistent symptoms following SARS-CoV-2 infection. EBioMedicine 85, 104293 (2022).

30. R. E. Hamlin, C. A. Blish, Challenges and opportunities in long COVID research. Immunity 57, 1195–1214 (2024).

31. C. Gaebler, Z. Wang, J. C. C. Lorenzi, F. Muecksch, S. Finkin, M. Tokuyama, A. Cho, M. Jankovic, D. Schaefer-Babajew, T. Y. Oliveira, M. Cipolla, C. Viant, C. O. Barnes, Y. Bram, G. Breton, T. Hägglöf, P. Mendoza, A. Hurley, M. Turroja, K. Gordon, K. G. Millard, V. Ramos, F. Schmidt, Y. Weisblum, D. Jha, M. Tankelevich, G. Martinez-Delgado, J. Yee, R. Patel, J. Dizon, C. Unson-O’Brien, I. Shimeliovich, D. F. Robbiani, Z. Zhao, A. Gazumyan, R. E. Schwartz, T. Hatziioannou, P. J. Bjorkman, S. Mehandru, P. D. Bieniasz, M. Caskey, M. C. Nussenzweig, Evolution of antibody immunity to SARS-CoV-2. Nature 591, 639–644 (2021).

32. S. R. Stein, S. C. Ramelli, A. Grazioli, J.-Y. Chung, M. Singh, C. K. Yinda, C. W. Winkler, J. Sun, J. M. Dickey, K. Ylaya, S. H. Ko, A. P. Platt, P. D. Burbelo, M. Quezado, S. Pittaluga, M. Purcell, V. J. Munster, F. Belinky, M. J. Ramos-Benitez, E. A. Boritz, I. A. Lach, D. L. Herr, J. Rabin, K. K. Saharia, R. J. Madathil, A. Tabatabai, S. Soherwardi, M. T. McCurdy, NIH COVID-19 Autopsy Consortium, K. E. Peterson, J. I. Cohen, E. de Wit, K. M. Vannella, S. M. Hewitt, D. E. Kleiner, D. S. Chertow, SARS-CoV-2 infection and persistence in the human body and brain at autopsy. Nature 612, 758–763 (2022).

33. Z. Swank, Y. Senussi, Z. Manickas-Hill, X. G. Yu, J. Z. Li, G. Alter, D. R. Walt, Persistent Circulating Severe Acute Respiratory Syndrome Coronavirus 2 Spike Is Associated With Post-acute Coronavirus Disease 2019 Sequelae. Clin. Infect. Dis. 76, e487–e490 (2023).

34. V. Craddock, A. Mahajan, L. Spikes, B. Krishnamachary, A. K. Ram, A. Kumar, L. Chen, P. Chalise, N. K. Dhillon, Persistent circulation of soluble and extracellular vesicle-linked Spike protein in individuals with postacute sequelae of COVID-19. J. Med. Virol. 95, e28568 (2023).

35. C. Schultheiß, E. Willscher, L. Paschold, C. Gottschick, B. Klee, L. Bosurgi, J. Dutzmann, D. Sedding, T. Frese, M. Girndt, J. I. Höll, M. Gekle, R. Mikolajczyk, M. Binder, Liquid biomarkers of macrophage dysregulation and circulating spike protein illustrate the biological heterogeneity in patients with post-acute sequelae of COVID-19. J. Med. Virol. 95, e28364 (2023).

36. M. J. Peluso, S. G. Deeks, M. Mustapic, D. Kapogiannis, T. J. Henrich, S. Lu, S. A. Goldberg, R. Hoh, J. Y. Chen, E. O. Martinez, J. D. Kelly, J. N. Martin, E. J. Goetzl, SARS-CoV-2 and Mitochondrial Proteins in Neural-Derived Exosomes of COVID-19. Ann. Neurol. 91, 772–781 (2022).

37. M. Hany, A. Zidan, M. Gaballa, M. Ibrahim, A. S. S. Agayby, A. A. Abouelnasr, E. Sheta, B. Torensma, Lingering SARS-CoV-2 in Gastric and Gallbladder Tissues of Patients with Previous COVID-19 Infection Undergoing Bariatric Surgery. Obes. Surg. 33, 139–148 (2023).

38. C. C. L. Cheung, D. Goh, X. Lim, T. Z. Tien, J. C. T. Lim, J. N. Lee, B. Tan, Z. E. A. Tay, W. Y. Wan, E. X. Chen, S. N. Nerurkar, S. Loong, P. C. Cheow, C. Y. Chan, Y. X. Koh, T. T. Tan, S. Kalimuddin, W. M. D. Tai, J. L. Ng, J. G.-H. Low, J. Yeong, K. H. Lim, Residual SARS-CoV-2 viral antigens detected in GI and hepatic tissues from five recovered patients with COVID-19. Gut 71, 226–229 (2022).

39. Q. Xu, P. Milanez-Almeida, A. J. Martins, A. J. Radtke, K. B. Hoehn, C. Oguz, J. Chen, C. Liu, J. Tang, G. Grubbs, S. Stein, S. Ramelli, J. Kabat, H. Behzadpour, M. Karkanitsa, J. Spathies, H. Kalish, L. Kardava, M. Kirby, F. Cheung, S. Preite, P. C. Duncker, M. M. Kitakule, N. Romero, D. Preciado, L. Gitman, G. Koroleva, G. Smith, A. Shaffer, I. T. McBain, P. J. McGuire, S. Pittaluga, R. N. Germain, R. Apps, D. M. Schwartz, K. Sadtler, S. Moir, D. S. Chertow, S. H. Kleinstein, S. Khurana, J. S. Tsang, P. Mudd, P. L. Schwartzberg, K. Manthiram, Adaptive immune responses to SARS-CoV-2 persist in the pharyngeal lymphoid tissue of children. Nat. Immunol. 24, 186–199 (2023).

40. F. Tejerina, P. Catalan, C. Rodriguez-Grande, J. Adan, C. Rodriguez-Gonzalez, P. Muñoz, T. Aldamiz, C. Diez, L. Perez, C. Fanciulli, D. Garcia de Viedma, Gregorio Marañon Microbiology ID COVID 19 Study Group, Post-COVID-19 syndrome. SARS-CoV-2 RNA detection in plasma, stool, and urine in patients with persistent symptoms after COVID-19. BMC Infect. Dis. 22, 211 (2022).

41. D. Goh, J. C. T. Lim, S. B. Fernaíndez, C. R. Joseph, S. G. Edwards, Z. W. Neo, J. N. Lee, S. G. Caballero, M. C. Lau, J. P. S. Yeong, Case report: Persistence of residual antigen and RNA of the SARS-CoV-2 virus in tissues of two patients with long COVID. Front. Immunol. 13, 939989 (2022).

42. A. Zollner, R. Koch, A. Jukic, A. Pfister, M. Meyer, A. Rössler, J. Kimpel, T. E. Adolph, H. Tilg, Postacute COVID-19 is Characterized by Gut Viral Antigen Persistence in Inflammatory Bowel Diseases. Gastroenterology 163, 495–506.e8 (2022).

43. G. D. de Melo, F. Lazarini, S. Levallois, C. Hautefort, V. Michel, F. Larrous, B. Verillaud, C. Aparicio, S. Wagner, G. Gheusi, L. Kergoat, E. Kornobis, F. Donati, T. Cokelaer, R. Hervochon, Y. Madec, E. Roze, D. Salmon, H. Bourhy, M. Lecuit, P.-M. Lledo, COVID-19-related anosmia is associated with viral persistence and inflammation in human olfactory epithelium and brain infection in hamsters. Sci. Transl. Med. 13, eabf8396 (2021).

44. Q. Yao, M. E. Doyle, Q.-R. Liu, A. Appleton, J. F. O’Connell, N. Weng, J. M. Egan, Long-Term Dysfunction of Taste Papillae in SARS-CoV-2. NEJM Evid. 2, EVIDoa2300046 (2023).

45. T. M. de Lima, R. B. Martins, C. S. Miura, M. V. O. Souza, M. H. A. Cassiano, T. S. Rodrigues, F. P. Veras, J. de F. Sousa, R. Gomes, G. M. de Almeida, S. R. Melo, G. C. da Silva, M. Dias, C. F. Capato, M. L. Silva, V. E. D. de B. Luiz, L. R. Carenzi, D. S. Zamboni, D. M. De M. Jorge, F. de Q. Cunha, E. Tamashiro, W. T. Anselmo-Lima, F. C. P. Valera, E. Arruda, Tonsils are major sites of persistence of SARS-CoV-2 in children. Microbiol. Spectr. 11, e0134723 (2023).

46. A. Natarajan, S. Zlitni, E. F. Brooks, S. E. Vance, A. Dahlen, H. Hedlin, R. M. Park, A. Han, D. T. Schmidtke, R. Verma, K. B. Jacobson, J. Parsonnet, H. F. Bonilla, U. Singh, B. A. Pinsky, J. R. Andrews, P. Jagannathan, A. S. Bhatt, Gastrointestinal symptoms and fecal shedding of SARS-CoV-2 RNA suggest prolonged gastrointestinal infection. Med N. Y. N 3, 371–387.e9 (2022).

47. M. J. Peluso, Z. N. Swank, S. A. Goldberg, S. Lu, T. Dalhuisen, E. Borberg, Y. Senussi, M. Luna, C. Chang Song, A. Clark, A. Zamora, M. Lew, B. Viswanathan, B. Huang, K. Anglin, Hoh, P. Y. Hsue, M. S. Durstenfeld, M. A. Spinelli, D. V. Glidden, T. J. Henrich, J. D. Kelly, G. Deeks, D. R. Walt, J. N. Martin, Plasma-based antigen persistence in the post-acute phase of SARS-CoV-2 infection. MedRxiv Prepr. Serv. Health Sci. , 2023.10.24.23297114 (2023).

48. M. Peghin, A. Palese, M. Venturini, M. De Martino, V. Gerussi, E. Graziano, G. Bontempo, F. Marrella, A. Tommasini, M. Fabris, F. Curcio, M. Isola, C. Tascini, Post-COVID-19 symptoms 6 months after acute infection among hospitalized and non-hospitalized patients. Clin. Microbiol. Infect. Off. Publ. Eur. Soc. Clin. Microbiol. Infect. Dis. 27, 1507–1513 (2021).

49. J. García-Abellán, S. Padilla, M. Fernández-González, J. A. García, V. Agulló, M. Andreo, S. Ruiz, A. Galiana, F. Gutiérrez, M. Masiá, Antibody Response to SARS-CoV-2 is Associated with Long-term Clinical Outcome in Patients with COVID-19: a Longitudinal Study. J. Clin. Immunol. 41, 1490–1501 (2021).

50. J. García-Abellán, M. Fernández, S. Padilla, J. A. García, V. Agulló, V. Lozano, N. Ena, L. García-Sánchez, F. Gutiérrez, M. Masiá, Immunologic phenotype of patients with long-COVID syndrome of 1-year duration. Front. Immunol. 13, 920627 (2022).

51. K. Yin, M. J. Peluso, X. Luo, R. Thomas, M.-G. Shin, J. Neidleman, A. Andrew, K. C. Young, T. Ma, R. Hoh, K. Anglin, B. Huang, U. Argueta, M. Lopez, D. Valdivieso, K. Asare, T.-M. Deveau, S. E. Munter, R. Ibrahim, L. Ständker, S. Lu, S. A. Goldberg, S. A. Lee, K. L. Lynch, J. D. Kelly, J. N. Martin, J. Münch, S. G. Deeks, T. J. Henrich, N. R. Roan, Long COVID manifests with T cell dysregulation, inflammation and an uncoordinated adaptive immune response to SARS-CoV-2. Nat. Immunol. , 1–8 (2024).

52. E. Y. Wang, T. Mao, J. Klein, Y. Dai, J. D. Huck, J. R. Jaycox, F. Liu, T. Zhou, B. Israelow, P. Wong, A. Coppi, C. Lucas, J. Silva, J. E. Oh, E. Song, E. S. Perotti, N. S. Zheng, S. Fischer, Campbell, J. B. Fournier, A. L. Wyllie, C. B. F. Vogels, I. M. Ott, C. C. Kalinich, M. E. Petrone, A. E. Watkins, Yale IMPACT Team, C. Dela Cruz, S. F. Farhadian, W. L. Schulz, S. Ma, N. D. Grubaugh, A. I. Ko, A. Iwasaki, A. M. Ring, Diverse functional autoantibodies in patients with COVID-19. Nature 595, 283–288 (2021).

53. C. Cervia, Y. Zurbuchen, P. Taeschler, T. Ballouz, D. Menges, S. Hasler, S. Adamo, M. E. Raeber, E. Bächli, A. Rudiger, M. Stüssi-Helbling, L. C. Huber, J. Nilsson, U. Held, M. A. Puhan, O. Boyman, Immunoglobulin signature predicts risk of post-acute COVID-19 syndrome. Nat. Commun. 13, 446 (2022).

54. A. G. Richter, A. M. Shields, A. Karim, D. Birch, S. E. Faustini, L. Steadman, K. Ward, T. Plant, G. Reynolds, T. Veenith, A. F. Cunningham, M. T. Drayson, D. C. Wraith, Establishing the prevalence of common tissue-specific autoantibodies following severe acute respiratory syndrome coronavirus 2 infection. Clin. Exp. Immunol. 205, 99–105 (2021).

55. K. Son, R. Jamil, A. Chowdhury, M. Mukherjee, C. Venegas, K. Miyasaki, K. Zhang, Z. Patel, B. Salter, A. C. Y. Yuen, K. S.-K. Lau, B. Cowbrough, K. Radford, C. Huang, M. Kjarsgaard, A. Dvorkin-Gheva, J. Smith, Q.-Z. Li, S. Waserman, C. J. Ryerson, P. Nair, T. Ho, Balakrishnan, I. Nazy, D. M. E. Bowdish, S. Svenningsen, C. Carlsten, M. Mukherjee, Circulating anti-nuclear autoantibodies in COVID-19 survivors predict long COVID symptoms. Eur. Respir. J. 61, 2200970 (2023).

56. G. Wallukat, B. Hohberger, K. Wenzel, J. Fürst, S. Schulze-Rothe, A. Wallukat, A.-S. Hönicke, J. Müller, Functional autoantibodies against G-protein coupled receptors in patients with persistent Long-COVID-19 symptoms. J. Transl. Autoimmun. 4, 100100 (2021).

57. F. Tesch, F. Ehm, A. Vivirito, D. Wende, M. Batram, F. Loser, S. Menzer, J. Jacob, M. Roessler, M. Seifert, B. Kind, C. König, C. Schulte, T. Buschmann, D. Hertle, P. Ballesteros, S. Baßler, B. Bertele, T. Bitterer, C. Riederer, F. Sobik, L. Reitzle, C. Scheidt-Nave, J. Schmitt, Incident autoimmune diseases in association with SARS-CoV-2 infection: a matched cohort study. Clin. Rheumatol. 42, 2905–2914 (2023).

58. C. Szewczykowski, C. Mardin, M. Lucio, G. Wallukat, J. Hoffmanns, T. Schröder, F. Raith, L. Rogge, F. Heltmann, M. Moritz, L. Beitlich, J. Schottenhamml, M. Herrmann, T. Harrer, M. Ganslmayer, F. E. Kruse, M. Kräter, J. Guck, R. Lämmer, M. Zenkel, A. Gießl, B. Hohberger, Long COVID: Association of Functional Autoantibodies against G-Protein-Coupled Receptors with an Impaired Retinal Microcirculation. Int. J. Mol. Sci. 23, 7209 (2022).

59. M. J. Peluso, T.-M. Deveau, S. E. Munter, D. Ryder, A. Buck, G. Beck-Engeser, F. Chan, S. Lu, S. A. Goldberg, R. Hoh, V. Tai, L. Torres, N. S. Iyer, M. Deswal, L. H. Ngo, M. Buitrago, Rodriguez, J. Y. Chen, B. C. Yee, A. Chenna, J. W. Winslow, C. J. Petropoulos, A. N. Deitchman, J. Hellmuth, M. A. Spinelli, M. S. Durstenfeld, P. Y. Hsue, J. D. Kelly, J. N. Martin, S. G. Deeks, P. W. Hunt, T. J. Henrich, Chronic viral coinfections differentially affect the likelihood of developing long COVID. J. Clin. Invest. 133, e163669 (2023).

60. J. E. Gold, R. A. Okyay, W. E. Licht, D. J. Hurley, Investigation of Long COVID Prevalence and Its Relationship to Epstein-Barr Virus Reactivation. Pathog. Basel Switz. 10, 763 (2021).

61. S. Zubchenko, I. Kril, O. Nadizhko, O. Matsyura, V. Chopyak, Herpesvirus infections and post-COVID-19 manifestations: a pilot observational study. Rheumatol. Int. 42, 1523–1530 (2022).

62. C. Cervia-Hasler, S. C. Brüningk, T. Hoch, B. Fan, G. Muzio, R. C. Thompson, L. Ceglarek, R. Meledin, P. Westermann, M. Emmenegger, P. Taeschler, Y. Zurbuchen, M. Pons, D. Menges, T. Ballouz, S. Cervia-Hasler, S. Adamo, M. Merad, A. W. Charney, M. Puhan, P. Brodin, J. Nilsson, A. Aguzzi, M. E. Raeber, C. B. Messner, N. D. Beckmann, K. Borgwardt, O. Boyman, Persistent complement dysregulation with signs of thromboinflammation in active Long Covid. Science 383, eadg7942 (2024).

63. E. Pretorius, M. Vlok, C. Venter, J. A. Bezuidenhout, G. J. Laubscher, J. Steenkamp, D. B. Kell, Persistent clotting protein pathology in Long COVID/Post-Acute Sequelae of COVID-19 (PASC) is accompanied by increased levels of antiplasmin. Cardiovasc. Diabetol. 20, 172 (2021).

64. D. Buonsenso, D. Di Giuda, L. Sigfrid, D. A. Pizzuto, G. Di Sante, C. De Rose, I. Lazzareschi, M. Sali, F. Baldi, D. P. R. Chieffo, D. Munblit, P. Valentini, Evidence of lung perfusion defects and ongoing inflammation in an adolescent with post-acute sequelae of SARS-CoV-2 infection. *Lancet Child Adolesc*. Health 5, 677–680 (2021).

65. E. Pasini, G. Corsetti, C. Romano, T. M. Scarabelli, C. Chen-Scarabelli, L. Saravolatz, F. S. Dioguardi, Serum Metabolic Profile in Patients With Long-Covid (PASC) Syndrome: Clinical Implications. Front. Med. 8 (2021) (available at https://www.frontiersin.org/articles/10.3389/fmed.2021.714426).

66. H. Fogarty, L. Townsend, H. Morrin, A. Ahmad, C. Comerford, E. Karampini, H. Englert, M. Byrne, C. Bergin, J. M. O’Sullivan, I. Martin-Loeches, P. Nadarajan, C. Bannan, P. W. Mallon, G. F. Curley, R. J. S. Preston, A. M. Rehill, D. McGonagle, C. Ni Cheallaigh, R. I. Baker, T. Renné, S. E. Ward, J. S. O’Donnell, Irish COVID-19 Vasculopathy Study (iCVS) investigators, Persistent endotheliopathy in the pathogenesis of long COVID syndrome. J. Thromb. Haemost. JTH 19, 2546–2553 (2021).

67. N. Prasannan, M. Heightman, T. Hillman, E. Wall, R. Bell, A. Kessler, L. Neave, A. Doyle, Devaraj, D. Singh, H.-M. Dehbi, M. Scully, Impaired exercise capacity in post-COVID-19 syndrome: the role of VWF-ADAMTS13 axis. Blood Adv. 6, 4041–4048 (2022).

68. B. Appelman, B. T. Charlton, R. P. Goulding, T. J. Kerkhoff, E. A. Breedveld, W. Noort, C. Offringa, F. W. Bloemers, M. van Weeghel, B. V. Schomakers, P. Coelho, J. J. Posthuma, E. Aronica, W. Joost Wiersinga, M. van Vugt, R. C. I. Wüst, Muscle abnormalities worsen after post-exertional malaise in long COVID. Nat. Commun. 15, 17 (2024).

69. D. B. Kell, G. J. Laubscher, E. Pretorius, A central role for amyloid fibrin microclots in long COVID/PASC: origins and therapeutic implications. Biochem. J. 479, 537–559 (2022).

70. M. Charnley, S. Islam, G. K. Bindra, J. Engwirda, J. Ratcliffe, J. Zhou, R. Mezzenga, M. D. Hulett, K. Han, J. T. Berryman, N. P. Reynolds, Neurotoxic amyloidogenic peptides in the proteome of SARS-COV2: potential implications for neurological symptoms in COVID-19. Nat. Commun. 13, 3387 (2022).

71. L. Guo, B. Appelman, K. Mooij-Kalverda, R. H. Houtkooper, M. van Weeghel, F. M. Vaz, Dijkhuis, T. Dekker, B. S. Smids, J. W. Duitman, M. Bugiani, P. Brinkman, J. J. Sikkens, H. A. A. Lavell, R. C. I. Wüst, M. van Vugt, R. Lutter, M. A. van Agtmael, A. G. Algera, B. Appelman, F. E. H. P. van Baarle, M. Beudel, H. J. Bogaard, M. Bomers, P. I. Bonta, L. D. J. Bos, M. Botta, J. de Brabander, G. J. de Bree, S. de Bruin, M. Bugiani, E. B. Bulle, O. Chouchane, A. P. M. Cloherty, D. Buis, M. C. F. J. de Rotte, M. Dijkstra, D. A. Dongelmans, R. W. G. Dujardin, P. E. Elbers, L. M. Fleuren, S. E. Geerlings, T. B. H. Geijtenbeek, A. R. J. Girbes, A. Goorhuis, M. P. Grobusch, L. A. Hagens, J. Hamann, V. C. Harris, R. Hemke, S. M. Hermans, L. M. A. Heunks, M. W. Hollmann, J. Horn, J. W. Hovius, M. D. de Jong, R. Koing, E. H. T. Lim, N. van Mourik, J. F. Nellen, E. J. Nossent, F. Paulus, E. Peters, D. Piña-Fuentes, T. van der Poll, B. Preckel, J. M. Prins, S. J. Raasveld, T. D. Y. Reijnders, M. Schinkel, F. a. P. Schrauwen, M. J. Schultz, A. R. Schuurman, J. Schuurmans, K. Sigaloff, M. A. Slim, P. Smeele, M. R. Smit, C. Stijnis, W. Stilma, C. E. Teunissen, P. Thoral, A. M. Tsonas, P. R. Tuinman, M. van der Valk, D. P. Veelo, C. Volleman, H. de Vries, L. A. van Vught, M. van Vugt, D. Wouters, A. H. Zwinderman, M. C. Brouwer, W. J. Wiersinga, A. P. J. Vlaar, D. van de Beek, Prolonged indoleamine 2,3-dioxygenase-2 activity and associated cellular stress in post-acute sequelae of SARS-CoV-2 infection. eBioMedicine 94 (2023), doi:10.1016/j.ebiom.2023.104729.

72. A. V. W. Nunn, G. W. Guy, W. Brysch, J. D. Bell, Understanding Long COVID; Mitochondrial Health and Adaptation—Old Pathways, New Problems. Biomedicines 10, 3113 (2022).

73. Q. Liu, Q. Su, F. Zhang, H. M. Tun, J. W. Y. Mak, G. C.-Y. Lui, S. S. S. Ng, J. Y. L. Ching, Li, W. Lu, C. Liu, C. P. Cheung, D. S. C. Hui, P. K. S. Chan, F. K. L. Chan, S. C. Ng, Multi-kingdom gut microbiota analyses define COVID-19 severity and post-acute COVID-19 syndrome. Nat. Commun. 13, 6806 (2022).

74. Q. Liu, J. W. Y. Mak, Q. Su, Y. K. Yeoh, G. C.-Y. Lui, S. S. S. Ng, F. Zhang, A. Y. L. Li, W. Lu, D. S.-C. Hui, P. K. Chan, F. K. L. Chan, S. C. Ng, Gut microbiota dynamics in a prospective cohort of patients with post-acute COVID-19 syndrome. Gut 71, 544–552 (2022).

75. V. L. Carneiro, K. M. Littlefield, R. Watson, B. E. Palmer, C. Lozupone, Inflammation-associated gut microbiome in postacute sequelae of SARS-CoV-2 points towards new therapeutic targets. Gut 73, 376–378 (2024).

76. B. Vestad, T. Ueland, T. V. Lerum, T. B. Dahl, K. Holm, A. Barratt-Due, T. Kåsine, A. M. Dyrhol-Riise, B. Stiksrud, K. Tonby, H. Hoel, I. C. Olsen, K. N. Henriksen, A. Tveita, R. Manotheepan, M. Haugli, R. Eiken, Å. Berg, B. Halvorsen, T. Lekva, T. Ranheim, A. E. Michelsen, A. B. Kildal, A. Johannessen, L. Thoresen, H. Skudal, B. R. Kittang, R. B. Olsen, C. M. Ystrøm, N. V. Skei, R. Hannula, S. Aballi, R. Kvåle, O. H. Skjønsberg, P. Aukrust, J. R. Hov, M. Trøseid, NOR-Solidarity study group, Respiratory dysfunction three months after severe COVID-19 is associated with gut microbiota alterations. J. Intern. Med. 291, 801–812 (2022).

77. A. C. Wong, A. S. Devason, I. C. Umana, T. O. Cox, L. Dohnalová, L. Litichevskiy, J. Perla, P. Lundgren, Z. Etwebi, L. T. Izzo, J. Kim, M. Tetlak, H. C. Descamps, S. L. Park, S. Wisser, A. D. McKnight, R. D. Pardy, J. Kim, N. Blank, S. Patel, K. Thum, S. Mason, J.-C. Beltra, M. F. Michieletto, S. F. Ngiow, B. M. Miller, M. J. Liou, B. Madhu, O. Dmitrieva-Posocco, A. S. Huber, P. Hewins, C. Petucci, C. P. Chu, G. Baraniecki-Zwil, L. B. Giron, A. E. Baxter, A. R. Greenplate, C. Kearns, K. Montone, L. A. Litzky, M. Feldman, J. Henao-Mejia, B. Striepen, H. Ramage, K. A. Jurado, K. E. Wellen, U. O’Doherty, M. Abdel-Mohsen, A. L. Landay, A. Keshavarzian, T. J. Henrich, S. G. Deeks, M. J. Peluso, N. J. Meyer, E. J. Wherry, B. A. Abramoff, S. Cherry, C. A. Thaiss, M. Levy, Serotonin reduction in post-acute sequelae of viral infection. Cell 0 (2023), doi:10.1016/j.cell.2023.09.013.

78. C. Pereira, B. H. L. Harris, M. Di Giovannantonio, C. Rosadas, C.-E. Short, R. Quinlan, M. Sureda-Vives, N. Fernandez, I. Day-Weber, M. Khan, F. Marchesin, K. Katsanovskaja, E. Parker, G. P. Taylor, R. S. Tedder, M. O. McClure, M. Dani, M. Fertleman, The Association Between Antibody Response to Severe Acute Respiratory Syndrome Coronavirus 2 Infection and Post-COVID-19 Syndrome in Healthcare Workers. J. Infect. Dis. 223, 1671–1676 (2021).

79. S. W. X. Ong, S.-W. Fong, B. E. Young, Y.-H. Chan, B. Lee, S. N. Amrun, R. S.-L. Chee, N. K.-W. Yeo, P. Tambyah, S. Pada, S. Y. Tan, Y. Ding, L. Renia, Y.-S. Leo, L. F. P. Ng, D. C. Lye, Persistent Symptoms and Association With Inflammatory Cytokine Signatures in Recovered Coronavirus Disease 2019 Patients. Open Forum Infect. Dis. 8, ofab156 (2021).

80. M. Alkhouli, A. Nanjundappa, F. Annie, M. C. Bates, D. L. Bhatt, Sex Differences in Case Fatality Rate of COVID-19: Insights From a Multinational Registry. Mayo Clin. Proc. 95, 1613– 1620 (2020).

81. Y. Meng, P. Wu, W. Lu, K. Liu, K. Ma, L. Huang, J. Cai, H. Zhang, Y. Qin, H. Sun, W. Ding, L. Gui, P. Wu, Sex-specific clinical characteristics and prognosis of coronavirus disease-19 infection in Wuhan, China: A retrospective study of 168 severe patients. PLoS Pathog. 16, e1008520 (2020).

82. S. Richardson, J. S. Hirsch, M. Narasimhan, J. M. Crawford, T. McGinn, K. W. Davidson, the Northwell COVID-19 Research Consortium, D. P. Barnaby, L. B. Becker, J. D. Chelico, S. Cohen, J. Cookingham, K. Coppa, M. A. Diefenbach, A. J. Dominello, J. Duer-Hefele, L. Falzon, J. Gitlin, N. Hajizadeh, T. G. Harvin, D. A. Hirschwerk, E. J. Kim, Z. M. Kozel, L. M. Marrast, J. N. Mogavero, G. A. Osorio, M. Qiu, T. P. Zanos, Presenting Characteristics, Comorbidities, and Outcomes Among 5700 Patients Hospitalized With COVID-19 in the New York City Area. JAMA 323, 2052–2059 (2020).

83. G. Lippi, C. Mattiuzzi, F. Sanchis-Gomar, B. M. Henry, Clinical and demographic characteristics of patients dying from COVID-19 in Italy vs China. J. Med. Virol. 92, 1759–1760 (2020).

84. T. Chen, D. Wu, H. Chen, W. Yan, D. Yang, G. Chen, K. Ma, D. Xu, H. Yu, H. Wang, T. Wang, W. Guo, J. Chen, C. Ding, X. Zhang, J. Huang, M. Han, S. Li, X. Luo, J. Zhao, Q. Ning, Clinical characteristics of 113 deceased patients with coronavirus disease 2019: retrospective study. BMJ 368, m1091 (2020).

85. J. Fabião, B. Sassi, E. F. Pedrollo, F. Gerchman, C. K. Kramer, C. B. Leitão, L. C. Pinto, Why do men have worse COVID-19-related outcomes? A systematic review and meta-analysis with sex adjusted for age. Braz. J. Med. Biol. Res. 55, e11711 (2022).

86. A. Doerre, G. Doblhammer, The influence of gender on COVID-19 infections and mortality in Germany: Insights from age- and gender-specific modeling of contact rates, infections, and deaths in the early phase of the pandemic. PloS One 17, e0268119 (2022).

87. C. H. Sudre, B. Murray, T. Varsavsky, M. S. Graham, R. S. Penfold, R. C. Bowyer, J. C. Pujol, K. Klaser, M. Antonelli, L. S. Canas, E. Molteni, M. Modat, M. Jorge Cardoso, A. May, S. Ganesh, R. Davies, L. H. Nguyen, D. A. Drew, C. M. Astley, A. D. Joshi, J. Merino, N. Tsereteli, T. Fall, M. F. Gomez, E. L. Duncan, C. Menni, F. M. K. Williams, P. W. Franks, A. T. Chan, J. Wolf, S. Ourselin, T. Spector, C. J. Steves, Attributes and predictors of long COVID. Nat. Med. 27, 626–631 (2021).

88. C. Chen, S. R. Haupert, L. Zimmermann, X. Shi, L. G. Fritsche, B. Mukherjee, Global Prevalence of Post-Coronavirus Disease 2019 (COVID-19) Condition or Long COVID: A Meta-Analysis and Systematic Review. J. Infect. Dis. 226, 1593–1607 (2022).

89. F. Di Gennaro, A. Belati, O. Tulone, L. Diella, D. Fiore Bavaro, R. Bonica, V. Genna, L. Smith, M. Trott, O. Bruyere, L. Mirarchi, C. Cusumano, L. J. Dominguez, A. Saracino, N. Veronese, M. Barbagallo, Incidence of long COVID-19 in people with previous SARS-Cov2 infection: a systematic review and meta-analysis of 120,970 patients. Intern. Emerg. Med. 18, 1573–1581 (2023).

90. S. Wulf Hanson, C. Abbafati, J. G. Aerts, Z. Al-Aly, C. Ashbaugh, T. Ballouz, O. Blyuss, P. Bobkova, G. Bonsel, S. Borzakova, D. Buonsenso, D. Butnaru, A. Carter, H. Chu, C. De Rose, M. Diab, E. Ekbom, M. El Tantawi, V. Fomin, R. Frithiof, A. Gamirova, P. V. Glybochko, J. A. Haagsma, S. Haghjooy Javanmard, E. B. Hamilton, G. Harris, M. H. Heijenbrok-Kal, R. Helbok, M. E. Hellemons, D. Hillus, S. M. Huijts, M. Hultström, W. Jassat, F. Kurth, I.-M. Larsson, M. Lipcsey, C. Liu, C. D. Loflin, A. Malinovschi, W. Mao, L. Mazankova, D. McCulloch, D. Menges, N. Mohammadifard, D. Munblit, N. A. Nekliudov, O. Ogbuoji, I. M. Osmanov, J. L. Peñalvo, M. S. Petersen, M. A. Puhan, M. Rahman, V. Rass, N. Reinig, G. M. Ribbers, A. Ricchiuto, S. Rubertsson, E. Samitova, N. Sarrafzadegan, A. Shikhaleva, K. E. Simpson, D. Sinatti, J. B. Soriano, E. Spiridonova, F. Steinbeis, A. A. Svistunov, P. Valentini, B. J. van de Water, R. van den Berg-Emons, E. Wallin, M. Witzenrath, Y. Wu, H. Xu, T. Zoller, C. Adolph, J. Albright, J. O. Amlag, A. Y. Aravkin, B. L. Bang-Jensen, C. Bisignano, R. Castellano, E. Castro, S. Chakrabarti, J. K. Collins, X. Dai, F. Daoud, C. Dapper, A. Deen, B. B. Duncan, M. Erickson, S. B. Ewald, A. J. Ferrari, A. D. Flaxman, N. Fullman, A. Gamkrelidze, J. R. Giles, G. Guo, S. I. Hay, J. He, M. Helak, E. N. Hulland, M. Kereselidze, K. J. Krohn, A. Lazzar-Atwood, A. Lindstrom, R. Lozano, D. C. Malta, J. Månsson, A. M. Mantilla Herrera, A. H. Mokdad, L. Monasta, S. Nomura, M. Pasovic, D. M. Pigott, R. C. Reiner, G. Reinke, A. L. P. Ribeiro, D. F. Santomauro, A. Sholokhov, E. E. Spurlock, R. Walcott, A. Walker, C. S. Wiysonge, P. Zheng, J. P. Bettger, C. J. L. Murray, T. Vos, Estimated Global Proportions of Individuals With Persistent Fatigue, Cognitive, and Respiratory Symptom Clusters Following Symptomatic COVID-19 in 2020 and 2021. JAMA 328, 1604–1615 (2022).

91. A. Subramanian, K. Nirantharakumar, S. Hughes, P. Myles, T. Williams, K. M. Gokhale, T. Taverner, J. S. Chandan, K. Brown, N. Simms-Williams, A. D. Shah, M. Singh, F. Kidy, K. Okoth, R. Hotham, N. Bashir, N. Cockburn, S. I. Lee, G. M. Turner, G. V. Gkoutos, O. L. Aiyegbusi, C. McMullan, A. K. Denniston, E. Sapey, J. M. Lord, D. C. Wraith, E. Leggett, C. Iles, T. Marshall, M. J. Price, S. Marwaha, E. H. Davies, L. J. Jackson, K. L. Matthews, J. Camaradou, M. Calvert, S. Haroon, Symptoms and risk factors for long COVID in non-hospitalized adults. Nat. Med. 28, 1706–1714 (2022).

92. E. J. Thompson, D. M. Williams, A. J. Walker, R. E. Mitchell, C. L. Niedzwiedz, T. C. Yang, C. F. Huggins, A. S. F. Kwong, R. J. Silverwood, G. Di Gessa, R. C. E. Bowyer, K. Northstone, B. Hou, M. J. Green, B. Dodgeon, K. J. Doores, E. L. Duncan, F. M. K. Williams, OpenSAFELY Collaborative, A. Steptoe, D. J. Porteous, R. R. C. McEachan, L. Tomlinson, B. Goldacre, P. Patalay, G. B. Ploubidis, S. V. Katikireddi, K. Tilling, C. T. Rentsch, N. J. Timpson, N. Chaturvedi, C. J. Steves, Long COVID burden and risk factors in 10 UK longitudinal studies and electronic health records. Nat. Commun. 13, 3528 (2022).

93. S. V. Sylvester, R. Rusu, B. Chan, M. Bellows, C. O’Keefe, S. Nicholson, Sex differences in sequelae from COVID-19 infection and in long COVID syndrome: a review. Curr. Med. Res. Opin. 38, 1391–1399 (2022).

94. F. Bai, D. Tomasoni, C. Falcinella, D. Barbanotti, R. Castoldi, G. Mulè, M. Augello, D. Mondatore, M. Allegrini, A. Cona, D. Tesoro, G. Tagliaferri, O. Viganò, E. Suardi, C. Tincati, T. Beringheli, B. Varisco, C. L. Battistini, K. Piscopo, E. Vegni, A. Tavelli, S. Terzoni, G. Marchetti, A. d’Arminio Monforte, Female gender is associated with long COVID syndrome: a prospective cohort study. Clin. Microbiol. Infect. Off. Publ. Eur. Soc. Clin. Microbiol. Infect. Dis. 28, 611.e9–611.e16 (2022).

95. K. S. Forsyth, N. Jiwrajka, C. D. Lovell, N. E. Toothacre, M. C. Anguera, The conneXion between sex and immune responses. Nat. Rev. Immunol. , 1–16 (2024).

96. S. L. Klein, K. L. Flanagan, Sex differences in immune responses. Nat. Rev. Immunol. 16, 626–638 (2016).

97. B. Huang, Y. Cai, N. Li, K. Li, Z. Wang, L. Li, L. Wu, M. Zhu, J. Li, Z. Wang, M. Wu, W. Li, W. Wu, L. Zhang, X. Xia, S. Wang, H. Chen, Q. Wang, Sex-based clinical and immunological differences in COVID-19. BMC Infect. Dis. 21, 647 (2021).

98. A. C. Moser, J. Z. Li, J. J. Eron, E. Aga, E. S. Daar, D. A. Wohl, R. W. Coombs, A. C. Javan, R. Bender Ignacio, P. Jagannathan, J. Ritz, S. F. Sieg, U. M. Parikh, M. D. Hughes, J. S. Currier, D. M. Smith, K. W. Chew, ACTIV-2/A5401 Study Team, Predictors of SARS-CoV-2 RNA From Nasopharyngeal Swabs and Concordance With Other Compartments in Nonhospitalized Adults With Mild to Moderate COVID-19. Open Forum Infect. Dis. 9, ofac618 (2022).

99. J. Q. Ho, M. R. Sepand, B. Bigdelou, T. Shekarian, R. Esfandyarpour, P. Chauhan, V. Serpooshan, L. K. Beura, G. Hutter, S. Zanganeh, The immune response to COVID-19: Does sex matter? Immunology 166, 429–443 (2022).

100. Y. Hao, S. Hao, E. Andersen-Nissen, W. M. Mauck, S. Zheng, A. Butler, M. J. Lee, A. J. Wilk, C. Darby, M. Zager, P. Hoffman, M. Stoeckius, E. Papalexi, E. P. Mimitou, J. Jain, A. Srivastava, T. Stuart, L. M. Fleming, B. Yeung, A. J. Rogers, J. M. McElrath, C. A. Blish, R. Gottardo, P. Smibert, R. Satija, Integrated analysis of multimodal single-cell data. Cell 184, 3573–3587.e29 (2021).

101. S. Li, N. Rouphael, S. Duraisingham, S. Romero-Steiner, S. Presnell, C. Davis, D. S. Schmidt, S. E. Johnson, A. Milton, G. Rajam, S. Kasturi, G. M. Carlone, C. Quinn, D. Chaussabel, A. K. Palucka, M. J. Mulligan, R. Ahmed, D. S. Stephens, H. I. Nakaya, B. Pulendran, Molecular signatures of antibody responses derived from a systems biology study of five human vaccines. Nat. Immunol. 15, 195–204 (2014).

102. C. M. L. de Padilla, T. B. Niewold, The Type I Interferons: Basic Concepts and Clinical Relevance in Immune-mediated Inflammatory Diseases. Gene 576, 14–21 (2016).

103. D. R. Dou, Y. Zhao, J. A. Belk, Y. Zhao, K. M. Casey, D. C. Chen, R. Li, B. Yu, S. Srinivasan, B. T. Abe, K. Kraft, C. Hellström, R. Sjöberg, S. Chang, A. Feng, D. W. Goldman, A. A. Shah, M. Petri, L. S. Chung, D. F. Fiorentino, E. K. Lundberg, A. Wutz, P. J. Utz, H. Y. Chang, Xist ribonucleoproteins promote female sex-biased autoimmunity. Cell 187, 733–749.e16 (2024).

104. D. Szappanos, R. Tschismarov, T. Perlot, S. Westermayer, K. Fischer, E. Platanitis, F. Kallinger, M. Novatchkova, C. Lassnig, M. Müller, V. Sexl, K. L. Bennett, M. Foong-Sobis, J. M. Penninger, T. Decker, The RNA helicase DDX3X is an essential mediator of innate antimicrobial immunity. PLoS Pathog. 14, e1007397 (2018).

105. M. H. Vogt, E. Goulmy, F. M. Kloosterboer, E. Blokland, R. A. de Paus, R. Willemze, J. H. Falkenburg, UTY gene codes for an HLA-B60-restricted human male-specific minor histocompatibility antigen involved in stem cell graft rejection: characterization of the critical polymorphic amino acid residues for T-cell recognition. Blood 96, 3126–3132 (2000).

106. E. H. Warren, M. A. Gavin, E. Simpson, P. Chandler, D. C. Page, C. Disteche, K. A. Stankey, P. D. Greenberg, S. R. Riddell, The human UTY gene encodes a novel HLA-B8-restricted H-Y antigen. J. Immunol. Baltim. Md 1950 164, 2807–2814 (2000).

107. L. Wu, J. Cao, W. L. Cai, S. M. Lang, J. R. Horton, D. J. Jansen, Z. Z. Liu, J. F. Chen, M. Zhang, B. T. Mott, K. Pohida, G. Rai, S. C. Kales, M. J. Henderson, X. Hu, A. Jadhav, D. J. Maloney, A. Simeonov, S. Zhu, A. Iwasaki, M. D. Hall, X. Cheng, G. S. Shadel, Q. Yan, KDM5 histone demethylases repress immune response via suppression of STING. PLoS Biol. 16, e2006134 (2018).

108. L. Xiu, B. Ma, L. Ding, Antioncogenic roles of USP9Y and DDX3Y in lung cancer: USP9Y stabilizes DDX3Y by preventing its degradation through deubiquitination. Acta Histochem. 126, 152132 (2024).

109. R. Browaeys, J. Gilis, C. Sang-Aram, P. D. Bleser, L. Hoste, S. Tavernier, D. Lambrechts, R. Seurinck, Y. Saeys, MultiNicheNet: a flexible framework for differential cell-cell communication analysis from multi-sample multi-condition single-cell transcriptomics data, 2023.06.13.544751 (2023).

110. M. J. Robertson, Role of chemokines in the biology of natural killer cells. J. Leukoc. Biol. 71, 173–183 (2002).

111. Z. Guo, C. Zhou, L. Zhou, Z. Wang, X. Zhu, X. Mu, Overexpression of DAPK1-mediated inhibition of IKKβ/CSN5/PD-L1 axis enhances natural killer cell killing ability and inhibits tumor immune evasion in gastric cancer. Cell. Immunol. 372, 104469 (2022).

112. B. M. Maślikowski, L. Wang, Y. Wu, B. Fielding, P.-A. Bédard, JunD/AP-1 Antagonizes the Induction of DAPK1 To Promote the Survival of v-Src-Transformed Cells. J. Virol. 91, 10.1128/jvi.01925-16 (2016).

113. N. C. Reich, A Death-Promoting Role for ISG54/IFIT2. J. Interferon Cytokine Res. 33, 199–205 (2013).

114. L. Schroeder, C. Herwartz, D. Jordanovski, G. Steger, ZNF395 Is an Activator of a Subset of IFN-Stimulated Genes. Mediators Inflamm. 2017, 1248201 (2017).

115. C. M. Dubois, F. Blanchette, M. H. Laprise, R. Leduc, F. Grondin, N. G. Seidah, Evidence that furin is an authentic transforming growth factor-beta1-converting enzyme. Am. J. Pathol. 158, 305–316 (2001).

116. X.-Q. Li, Q.-Q. Zhang, H.-Y. Zhang, X.-H. Guo, H.-Q. Fan, L.-X. Liu, Interaction between insulin-like growth factor binding protein-related protein 1 and transforming growth factor beta 1 in primary hepatic stellate cells. Hepatobiliary Pancreat. Dis. Int. HBPD INT 16, 395–404 (2017).

117. G. Zhang, J. Lu, M. Yang, Y. Wang, H. Liu, C. Xu, Elevated GALNT10 expression identifies immunosuppressive microenvironment and dismal prognosis of patients with high grade serous ovarian cancer. Cancer Immunol. Immunother. CII 69, 175–187 (2020).

118. S. A. Cain, S. Woods, M. Singh, S. J. Kimber, C. Baldock, ADAMTS6 cleaves the large latent TGFβ complex and increases the mechanotension of cells to activate TGFβ. Matrix Biol. 114, 18–34 (2022).

119. K. Yanagisawa, H. Osada, A. Masuda, M. Kondo, T. Saito, Y. Yatabe, K. Takagi, T. Takahashi, T. Takahashi, Induction of apoptosis by Smad3 and down-regulation of Smad3 expression in response to TGF-β in human normal lung epithelial cells. Oncogene 17, 1743– 1747 (1998).

120. D. S. Matassa, M. R. Amoroso, F. Maddalena, M. Landriscina, F. Esposito, New insights into TRAP1 pathway. Am. J. Cancer Res. 2, 235–248 (2012).

121. G. Hua, Q. Zhang, Z. Fan, Heat shock protein 75 (TRAP1) antagonizes reactive oxygen species generation and protects cells from granzyme M-mediated apoptosis. J. Biol. Chem. 282, 20553–20560 (2007).

122. S. Taveirne, S. Wahlen, W. Van Loocke, L. Kiekens, E. Persyn, E. Van Ammel, K. De Mulder, J. Roels, L. Tilleman, M. Aumercier, P. Matthys, F. Van Nieuwerburgh, T. C. C. Kerre, T. Taghon, P. Van Vlierberghe, B. Vandekerckhove, G. Leclercq, The transcription factor ETS1 is an important regulator of human NK cell development and terminal differentiation. Blood 136, 288–298 (2020).

123. D. Chen, T.-X. Tang, H. Deng, X.-P. Yang, Z.-H. Tang, Interleukin-7 Biology and Its Effects on Immune Cells: Mediator of Generation, Differentiation, Survival, and Homeostasis. Front. Immunol. 12 (2021) (available at https://www.frontiersin.org/articles/10.3389/fimmu.2021.747324).

124. J.-H. Park, Q. Yu, B. Erman, J. S. Appelbaum, D. Montoya-Durango, H. L. Grimes, A. Singer, Suppression of IL7Rα Transcription by IL-7 and Other Prosurvival Cytokines: A Novel Mechanism for Maximizing IL-7-Dependent T Cell Survival. Immunity 21, 289–302 (2004).

125. Y. K. Takada, J. Yu, M. Shimoda, Y. Takada, Integrin Binding to the Trimeric Interface of CD40L Plays a Critical Role in CD40/CD40L Signaling. J. Immunol. Baltim. Md 1950 203, 1383–1391 (2019).

126. Y. K. Takada, M. Shimoda, E. Maverakis, B. H. Felding, R. H. Cheng, Y. Takada, Soluble CD40L activates soluble and cell-surface integrin αvβ3, α5β1, and α4β1 by binding to the allosteric ligand-binding site (site 2). J. Biol. Chem. 296, 100399 (2021).

127. Y. Wang, D. Gao, K. E. Lunsford, W. L. Frankel, G. L. Bumgardner, Targeting LFA-1 synergizes with CD40/CD40L blockade for suppression of both CD4-dependent and CD8-dependent rejection. Am. J. Transplant. Off. J. Am. Soc. Transplant. Am. Soc. Transpl. Surg. 3, 1251–1258 (2003).

128. A. Sinistro, C. Almerighi, C. Ciaprini, S. Natoli, E. Sussarello, S. Di Fino, F. Calò-Carducci, G. Rocchi, A. Bergamini, Downregulation of CD40 ligand response in monocytes from sepsis patients. Clin. Vaccine Immunol. CVI 15, 1851–1858 (2008).

129. R. Elgueta, M. J. Benson, V. C. de Vries, A. Wasiuk, Y. Guo, R. J. Noelle, Molecular mechanism and function of CD40/CD40L engagement in the immune system. Immunol. Rev. 229, 10.1111/j.1600-065X.2009.00782.x (2009).

130. P.-Y. Perera, J. H. Lichy, T. A. Waldmann, L. P. Perera, The role of Interleukin-15 in inflammation and immune responses to infection: implications for its therapeutic use. Microbes Infect. Inst. Pasteur 14, 247–261 (2012).

131. G. Tau, P. Rothman, Biologic functions of the IFN-γ receptors. Allergy 54, 1233–1251 (1999).

132. J. M. Pimentel, J.-Y. Zhou, G. S. Wu, The Role of TRAIL in Apoptosis and Immunosurveillance in Cancer. Cancers 15, 2752 (2023).

133. T. V. Zhao, Z. Hu, S. Ohtsuki, K. Jin, B. Wu, G. J. Berry, R. L. Frye, J. J. Goronzy, C. M. Weyand, Hyperactivity of the CD155 immune checkpoint suppresses anti-viral immunity in patients with coronary artery disease. *Nat*. Cardiovasc. Res. 1, 634–648 (2022).

134. N. Reymond, A.-M. Imbert, E. Devilard, S. Fabre, C. Chabannon, L. Xerri, C. Farnarier, C. Cantoni, C. Bottino, A. Moretta, P. Dubreuil, M. Lopez, DNAM-1 and PVR regulate monocyte migration through endothelial junctions. J. Exp. Med. 199, 1331–1341 (2004).

135. B. Dobosh, K. Zandi, D. M. Giraldo, S. L. Goh, K. Musall, M. Aldeco, J. LeCher, V. D. Giacalone, J. Yang, D. J. Eddins, M. Bhasin, E. Ghosn, V. Sukhatme, R. F. Schinazi, R. Tirouvanziam, Baricitinib attenuates the proinflammatory phase of COVID-19 driven by lung-infiltrating monocytes. Cell Rep. 39, 110945 (2022).

136. B. S. Stikker, G. Stik, A. F. van Ouwerkerk, L. Trap, S. Spicuglia, R. W. Hendriks, R. Stadhouders, Severe COVID-19-associated variants linked to chemokine receptor gene control in monocytes and macrophages. Genome Biol. 23, 96 (2022).

137. J. W. Schoggins, C. M. Rice, Interferon-stimulated genes and their antiviral effector functions. Curr. Opin. Virol. 1, 519–525 (2011).

138. O. Grünvogel, K. Esser-Nobis, A. Reustle, P. Schult, B. Müller, P. Metz, M. Trippler, M. P. Windisch, M. Frese, M. Binder, O. Fackler, R. Bartenschlager, A. Ruggieri, V. Lohmann, DDX60L Is an Interferon-Stimulated Gene Product Restricting Hepatitis C Virus Replication in Cell Culture. J. Virol. 89, 10548–10568 (2015).

139. X. Meng, D. Yang, R. Yu, H. Zhu, EPSTI1 Is Involved in IL-28A-Mediated Inhibition of HCV Infection. Mediators Inflamm. 2015, 716315 (2015).

140. S. Parthasarathy, A. R. Fehr, PARP14: A key ADP-ribosylating protein in host-virus interactions? PLoS Pathog. 18, e1010535 (2022).

141. P. Hubel, C. Urban, V. Bergant, W. M. Schneider, B. Knauer, A. Stukalov, P. Scaturro, A. Mann, L. Brunotte, H. H. Hoffmann, J. W. Schoggins, M. Schwemmle, M. Mann, C. M. Rice, A. Pichlmair, A protein-interaction network of interferon-stimulated genes extends the innate immune system landscape. Nat. Immunol. 20, 493–502 (2019).

142. J. Verhelst, E. Parthoens, B. Schepens, W. Fiers, X. Saelens, Interferon-inducible protein Mx1 inhibits influenza virus by interfering with functional viral ribonucleoprotein complex assembly. J. Virol. 86, 13445–13455 (2012).

143. A. Di Pietro, A. Kajaste-Rudnitski, A. Oteiza, L. Nicora, G. J. Towers, N. Mechti, E. Vicenzi, TRIM22 inhibits influenza A virus infection by targeting the viral nucleoprotein for degradation. J. Virol. 87, 4523–4533 (2013).

144. Z. Dong, Q. Yan, W. Cao, Z. Liu, X. Wang, Identification of key molecules in COVID-19 patients significantly correlated with clinical outcomes by analyzing transcriptomic data. Front. Immunol. 13, 930866 (2022).

145. C. Mei, F. Meng, X. Wang, S. Yan, Q. Zheng, X. Zhang, W. Fu, J. Xue, S. Wang, Y. He, X. Sun, X. Jiang, Y. Wang, CD30L is involved in the regulation of the inflammatory response through inducing homing and differentiation of monocytes via CCL2/CCR2 axis and NF-κB pathway in mice with colitis. Int. Immunopharmacol. 110, 108934 (2022).

146. K. Baillie, H. E. Davies, S. B. K. Keat, K. Ladell, K. L. Miners, S. A. Jones, E. Mellou, E. J. M. Toonen, D. A. Price, B. P. Morgan, W. M. Zelek, Complement dysregulation is a prevalent and therapeutically amenable feature of long COVID. Med N. Y. N 5, 239–253.e5 (2024).

147. M. Miyashita, H. Oshiumi, M. Matsumoto, T. Seya, DDX60, a DEXD/H box helicase, is a novel antiviral factor promoting RIG-I-like receptor-mediated signaling. Mol. Cell. Biol. 31, 3802–3819 (2011).

148. M. L. DeDiego, L. Martinez-Sobrido, D. J. Topham, Novel Functions of IFI44L as a Feedback Regulator of Host Antiviral Responses. J. Virol. 93, e01159–19 (2019).

149. Q. Zheng, D. Wang, R. Lin, Q. Lv, W. Wang, IFI44 is an immune evasion biomarker for SARS-CoV-2 and Staphylococcus aureus infection in patients with RA. Front. Immunol. 13, 1013322 (2022).

150. I. Aksentijevich, S. L. Masters, P. J. Ferguson, P. Dancey, J. Frenkel, A. van Royen-Kerkhoff, R. Laxer, U. Tedgård, E. W. Cowen, T.-H. Pham, M. Booty, J. D. Estes, N. G. Sandler, N. Plass, D. L. Stone, M. L. Turner, S. Hill, J. A. Butman, R. Schneider, P. Babyn, H. I. El-Shanti, E. Pope, K. Barron, X. Bing, A. Laurence, C.-C. R. Lee, D. Chapelle, G. I. Clarke, K. Ohson, M. Nicholson, M. Gadina, B. Yang, B. D. Korman, P. K. Gregersen, P. M. van Hagen, A. E. Hak, M. Huizing, P. Rahman, D. C. Douek, E. F. Remmers, D. L. Kastner, R. Goldbach-Mansky, An autoinflammatory disease with deficiency of the interleukin-1-receptor antagonist. N. Engl. J. Med. 360, 2426–2437 (2009).

151. Q. Guo, Y. Jin, X. Chen, X. Ye, X. Shen, M. Lin, C. Zeng, T. Zhou, J. Zhang, NF-κB in biology and targeted therapy: new insights and translational implications. Signal Transduct. Target. Ther. 9, 53 (2024).

152. A. Oeckinghaus, S. Ghosh, The NF-κB Family of Transcription Factors and Its Regulation. Cold Spring Harb. Perspect. Biol. 1, a000034 (2009).

153. T. D. Gilmore, S. Gerondakis, The c-Rel Transcription Factor in Development and Disease. Genes Cancer 2, 695–711 (2011).

154. V. Atsaves, V. Leventaki, G. Z. Rassidakis, F. X. Claret, AP-1 Transcription Factors as Regulators of Immune Responses in Cancer. Cancers 11, 1037 (2019).

155. A. G. Bassuk, J. M. Leiden, A direct physical association between ETS and AP-1 transcription factors in normal human T cells. Immunity 3, 223–237 (1995).

156. L. A. Garrett-Sinha, Review of Ets1 structure, function, and roles in immunity. Cell. Mol. Life Sci. CMLS 70, 3375–3390 (2013).

157. C. J. Kim, C.-G. Lee, J.-Y. Jung, A. Ghosh, S. N. Hasan, S.-M. Hwang, H. Kang, C. Lee, G.-C. Kim, D. Rudra, C.-H. Suh, S.-H. Im, The Transcription Factor Ets1 Suppresses T Follicular Helper Type 2 Cell Differentiation to Halt the Onset of Systemic Lupus Erythematosus. Immunity 49, 1034–1048.e8 (2018).

158. A. M. Miller, Role of IL-33 in inflammation and disease. J. Inflamm. Lond. Engl. 8, 22 (2011).

159. W. Saikruang, L. Ang Yan Ping, H. Abe, D. M. Kasumba, H. Kato, T. Fujita, The RNA helicase DDX3 promotes IFNB transcription via enhancing IRF-3/p300 holocomplex binding to the IFNB promoter. Sci. Rep. 12, 3967 (2022).

160. A. Bodansky, C.-Y. Wang, A. Saxena, A. Mitchell, A. F. Kung, S. Takahashi, K. Anglin, B. Huang, R. Hoh, S. Lu, S. A. Goldberg, J. Romero, B. Tran, R. Kirtikar, H. Grebe, M. So, B. Greenhouse, M. S. Durstenfeld, P. Y. Hsue, J. Hellmuth, J. D. Kelly, J. N. Martin, M. S. Anderson, S. G. Deeks, T. J. Henrich, J. L. DeRisi, M. J. Peluso, Autoantigen profiling reveals a shared post-COVID signature in fully recovered and long COVID patients. JCI Insight 8, e169515 (2023).

161. M. Witkowski, C. Tizian, M. Ferreira-Gomes, D. Niemeyer, T. C. Jones, F. Heinrich, S. Frischbutter, S. Angermair, T. Hohnstein, I. Mattiola, P. Nawrath, S. McEwen, S. Zocche, E. Viviano, G. A. Heinz, M. Maurer, U. Kölsch, R. L. Chua, T. Aschman, C. Meisel, J. Radke, B. Sawitzki, J. Roehmel, K. Allers, V. Moos, T. Schneider, L. Hanitsch, M. A. Mall, C. Conrad, H. Radbruch, C. U. Duerr, J. A. Trapani, E. Marcenaro, T. Kallinich, V. M. Corman, F. Kurth, L. E. Sander, C. Drosten, S. Treskatsch, P. Durek, A. Kruglov, A. Radbruch, M.-F. Mashreghi, A. Diefenbach, Untimely TGFβ responses in COVID-19 limit antiviral functions of NK cells. Nature 600, 295–301 (2021).

162. L. Gorelik, S. Constant, R. A. Flavell, Mechanism of transforming growth factor beta-induced inhibition of T helper type 1 differentiation. J. Exp. Med. 195, 1499–1505 (2002).

163. L. Gorelik, R. A. Flavell, Transforming growth factor-beta in T-cell biology. Nat. Rev. Immunol. 2, 46–53 (2002).

164. S. Blundell, K. K. Ray, M. Buckland, P. D. White, Chronic fatigue syndrome and circulating cytokines: A systematic review. Brain. Behav. Immun. 50, 186–195 (2015).

165. Julio Silva, Takehiro Takahashi, Jamie Wood, Peiwen Lu, Alexandra Tabachnikova, Jeff R. Gehlhausen, Kerrie Greene, Bornali Bhattacharjee, Valter Silva Monteiro, Carolina Lucas, Rahul M. Dhodapkar, Laura Tabacof, Mario Peña-Hernandez, Kathy Kamath, Tianyang Mao, Dayna Mccarthy, Ruslan Medzhitov, David van Dijk, Harlan M. Krumholz, Leying Guan, David Putrino, Akiko Iwasaki, Sex differences in symptomatology and immune profiles of Long COVID. medRxiv , 2024.02.29.24303568 (2024).

166. C. Riou, A. R. Dumont, B. Yassine-Diab, E. K. Haddad, R.-P. Sekaly, IL-4 influences the differentiation and the susceptibility to activation-induced cell death of human naive CD8+ T cells. Int. Immunol. 18, 827–835 (2006).

167. D. K. Wijesundara, D. C. Tscharke, R. J. Jackson, C. Ranasinghe, Reduced interleukin-4 receptor α expression on CD8+ T cells correlates with higher quality anti-viral immunity. PloS One 8, e55788 (2013).

168. M. Castro, J. Corren, I. D. Pavord, J. Maspero, S. Wenzel, K. F. Rabe, W. W. Busse, L. Ford, L. Sher, J. M. FitzGerald, C. Katelaris, Y. Tohda, B. Zhang, H. Staudinger, G. Pirozzi, N. Amin, M. Ruddy, B. Akinlade, A. Khan, J. Chao, R. Martincova, N. M. H. Graham, J. D. Hamilton, B. N. Swanson, N. Stahl, G. D. Yancopoulos, A. Teper, Dupilumab Efficacy and Safety in Moderate-to-Severe Uncontrolled Asthma. N. Engl. J. Med. 378, 2486–2496 (2018).

169. Long COVID - Household Pulse Survey - COVID-19 (2024) (available at https://www.cdc.gov/nchs/covid19/pulse/long-covid.htm).

170. K. B. Jacobson, M. Rao, H. Bonilla, A. Subramanian, I. Hack, M. Madrigal, U. Singh, P. Jagannathan, P. Grant, Patients With Uncomplicated Coronavirus Disease 2019 (COVID-19) Have Long-Term Persistent Symptoms and Functional Impairment Similar to Patients with Severe COVID-19: A Cautionary Tale During a Global Pandemic. Clin. Infect. Dis. Off. Publ. Infect. Dis. Soc. Am. 73, e826–e829 (2021).

171. P. A. Harris, R. Taylor, R. Thielke, J. Payne, N. Gonzalez, J. G. Conde, Research electronic data capture (REDCap)--a metadata-driven methodology and workflow process for providing translational research informatics support. J. Biomed. Inform. 42, 377–381 (2009).

172. P. A. Harris, R. Taylor, B. L. Minor, V. Elliott, M. Fernandez, L. O’Neal, L. McLeod, G. Delacqua, F. Delacqua, J. Kirby, S. N. Duda, REDCap Consortium, The REDCap consortium: Building an international community of software platform partners. J. Biomed. Inform. 95, 103208 (2019).

173. P. Jagannathan, J. R. Andrews, H. Bonilla, H. Hedlin, K. B. Jacobson, V. Balasubramanian, Purington, S. Kamble, C. R. de Vries, O. Quintero, K. Feng, C. Ley, D. Winslow, J. Newberry, K. Edwards, C. Hislop, I. Choong, Y. Maldonado, J. Glenn, A. Bhatt, C. Blish, T. Wang, C. Khosla, B. A. Pinsky, M. Desai, J. Parsonnet, U. Singh, Peginterferon Lambda-1a for treatment of outpatients with uncomplicated COVID-19: a randomized placebo-controlled trial. Nat. Commun. 12, 1967 (2021).

174. M. Holubar, A. Subramanian, N. Purington, H. Hedlin, B. Bunning, K. S. Walter, H. Bonilla, A. Boumis, M. Chen, K. Clinton, L. Dewhurst, C. Epstein, P. Jagannathan, R. H. Kaszynski, L. Panu, J. Parsonnet, E. L. Ponder, O. Quintero, E. Sefton, U. Singh, L. Soberanis, H. Truong, J. R. Andrews, M. Desai, C. Khosla, Y. Maldonado, Favipiravir for Treatment of Outpatients With Asymptomatic or Uncomplicated Coronavirus Disease 2019: A Double-Blind, Randomized, Placebo-Controlled, Phase 2 Trial. Clin. Infect. Dis. Off. Publ. Infect. Dis. Soc. Am. 75, 1883–1892 (2022).

175. E. R. Zunder, R. Finck, G. K. Behbehani, E. D. Amir, S. Krishnaswamy, V. D. Gonzalez, C. G. Lorang, Z. Bjornson, M. H. Spitzer, B. Bodenmiller, W. J. Fantl, D. Pe’er, G. P. Nolan, Palladium-based mass tag cell barcoding with a doublet-filtering scheme and single-cell deconvolution algorithm. Nat. Protoc. 10, 316–333 (2015).

176. R. Finck, E. F. Simonds, A. Jager, S. Krishnaswamy, K. Sachs, W. Fantl, D. Pe’er, G. P. Nolan, S. C. Bendall, Normalization of mass cytometry data with bead standards. Cytometry A **83A**, 483–494 (2013).

177. H. Crowell, V. Zanotelli, S. Chevrier, M. Robinson, CATALYST: Cytometry dATa anALYSis ToolsBioconductor (available at http://bioconductor.org/packages/CATALYST/).

178. A. Liaw, M. Wiener, Classification and regression by randomForest. *R News* **2**, 18–22 (2002).

179. K. Nevola (kathy-nevola), M. Sandin (marisand), J. Guess (jrguess), S. Forsberg (simfor), C. Cambronero (Orbmac), P. Pucholt (AskPascal), B. Zhang (boxizhang), M. Sheikhi (MasoumehSheikhi), K. Diamanti (klevdiamanti), A. Kar (amrita-kar), L. Conze (leiliuC), K. Hodén (kristianHoden), P. Eriksson (b_watcher), N. Moloney, B. Lötstedt, E. Sprecher, J. Barbagallo (jbarbagallo), O. Mansson (olofmansson), O. Caster (OlaCaster), Olink, OlinkAnalyze: Facilitate Analysis of Proteomic Data from Olink (2024) (available at https://cran.r-project.org/web/packages/OlinkAnalyze/index.html).

180. Azimuth (available at https://azimuth.hubmapconsortium.org/references/human_pbmc/).

181. D. J. McCarthy, K. R. Campbell, A. T. L. Lun, Q. F. Wills, Scater: pre-processing, quality control, normalization and visualization of single-cell RNA-seq data in R. Bioinformatics 33, 1179–1186 (2017).

182. A. T. L. Lun, D. J. McCarthy, J. C. Marioni, A step-by-step workflow for low-level analysis of single-cell RNA-seq data with Bioconductor. F1000R*esearch* **5**, 2122 (2016).

183. M. I. Love, W. Huber, S. Anders, Moderated estimation of fold change and dispersion for RNA-seq data with DESeq2. Genome Biol. 15, 550 (2014).

184. H. L. Crowell, P. Germain, C. Soneson, A. Sonrel, M. D. Robinson, muscat: Multi-sample multi-group scRNA-seq data analysis tools. (available at https://github.com/HelenaLC/muscat).

185. D. Beisser, G. W. Klau, T. Dandekar, T. Müller, M. T. Dittrich, BioNet: an R-Package for the functional analysis of biological networks. Bioinforma. Oxf. Engl. 26, 1129–1130 (2010).

186. D. Szklarczyk, R. Kirsch, M. Koutrouli, K. Nastou, F. Mehryary, R. Hachilif, A. L. Gable, T. Fang, N. T. Doncheva, S. Pyysalo, P. Bork, L. J. Jensen, C. von Mering, The STRING database in 2023: protein-protein association networks and functional enrichment analyses for any sequenced genome of interest. Nucleic Acids Res. 51, D638–D646 (2023).

187. F. Briatte, M. Bojanowski, M. Canouil, Z. Charlop-Powers, J. C. Fisher, K. Johnson, T. Rinker, ggnetwork: Geometries to Plot Networks with “ggplot2” (2024) (available at https://cran.r-project.org/web/packages/ggnetwork/index.html).

188. F. J. Hartmann, E. F. Simonds, S. C. Bendall, A Universal Live Cell Barcoding-Platform for Multiplexed Human Single Cell Analysis. Sci. Rep. 8, 10770 (2018).

