## Supplementary Materials for "Sex differences and immune correlates of Long COVID development, persistence, and resolution"

### MATERIALS AND METHODS

#### Mass cytometry reagents

PBMC surface and intracellular antibodies were either purchased pre-conjugated to metals from Fluidigm/Standard BioTools or purchased from Biolegend or R&D Systems and conjugated using Fluidigm Maxpar® X8 labeling kits or IONpath MIBItag conjugation kit (for 157Gd only). Antibody-metal conjugations were performed according to manufacturers' instructions with the following deviations for the Maxpar® X8 labeling kits: 30kDa filters were used instead of 50kDa filters; after the final washes, metal-conjugated antibody was recovered in W buffer, the concentration was determined using Qubit 4 Fluorometer (ThermoFisher) per manufacturer instructions, and conjugated antibodies were stored at 4°C for up to 1 month without antibody stabilization buffer. Conjugated antibodies were then titrated, pre-combined into either surface or intracellular staining cocktails (Table S5-6) to ensure staining consistency, filtered through Millex®-VV 0.1µm filter (Millipore #SLVV033RS), aliquoted for single use, and frozen at -80°C for long-term storage.

Live cell barcoding was adapted from Hartmann et al. *(188)*. Briefly, anti-b2-microglobulin mouse monoclonal antibody (clone: 2M2; Biolegend) was conjugated to 113In and 115In via IONpath MIBItag conjugation kits per manufacturer instructions, as well as to Cell-ID cisplatin 194Pt, 195Pt, 196Pt, and 198Pt (Fluidigm/Standard BioTools) according to the Fluidigm Maxpar® X8 labeling kit protocol. Rather than using a polymer with lanthanide metal per manufacturer protocol, a working dilution of each Cell-ID cisplatin (194Pt, 195Pt, 196Pt, and 198Pt) was made by diluting 20µL of the 1mM Cell-ID cisplatin stock solution to 1mL of C-buffer. These cisplatin working solutions were each conjugated to anti-b2-microglobulin antibody adding 400µL of cisplatin working solution to the 30kDa filter with the purified partially reduced antibody, resuspending the antibody, combining this resuspended antibody with the remaining 600µL of cisplatin working solution, incubating the 1mL antibody-cisplatin solution in a 37°C water bath for 90 minutes, adding half of the solution to 30kDa filter, centrifuging this solution at 12,000 xg for 15 minutes at room temperature, discarding the flow-through, adding the remaining half of the solution into the same 30 kDa filter, repeating centrifugation, and discarding the flow through. Metal conjugated antibody was washed with W-buffer as in Fluidigm Maxpar® X8 labeling kit protocol. The metal conjugated antibodies were recovered, the concentrations were quantified, and the optimal staining concentrations were determined, as described above. For staining consistency, barcodes were pre-combined in a six-choose-three scheme, aliquoted for single use, and frozen at -80°C for long-term storage.

PdCl viability staining was made as previously described *(188)* by re-suspending (Ethylenediamine)palladium(II) chloride (PdCl) (Sigma-Aldrich #574902, CAS #15020-99-2) in DMSO to a final concentration of 100mM, which was then pre-conditioned in a 37°C water bath for 48 hours, subsequently aliquoted into 10µL aliquots, and frozen at -20°C for long-term storage. Cell ID Intercalator (191Ir, 193Ir) at 500µM was purchased from Fluidigm/Standard BioTools, aliquoted upon arrival, and stored at -20°C. Intercalator was titrated for optimal signal intensity between 300 and 1000 dual counts in the 191Ir channel per manufacturer recommendations.

### SUPPLEMENTARY FIGURES

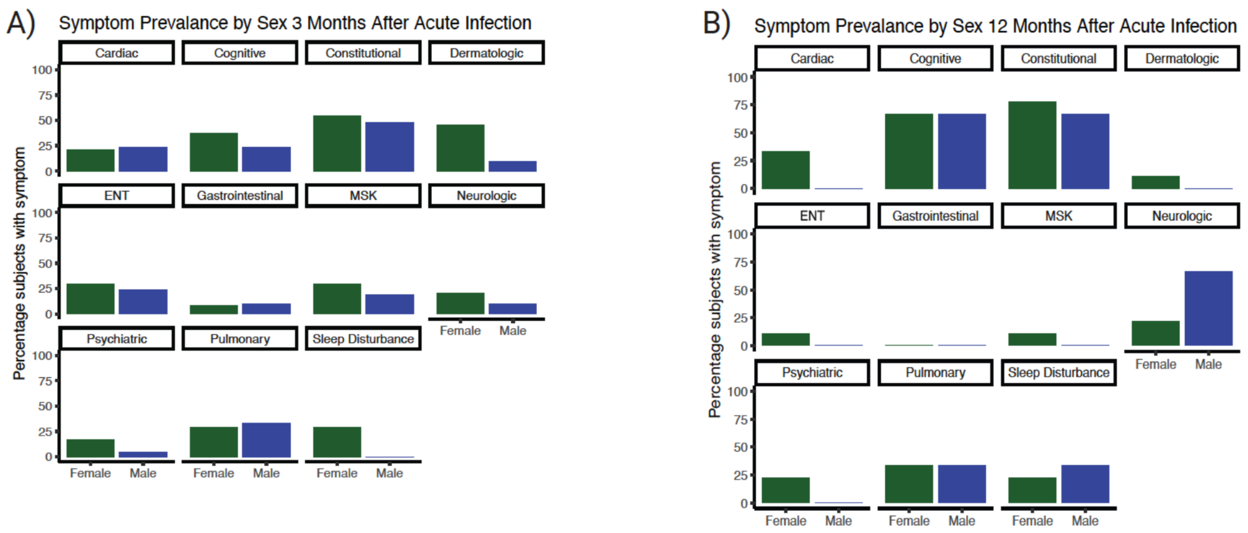


**Fig. S1. Symptom prevalence of subjects with LC at (A)** 3 months after acute infection (n=36), and **(B)** 12 months after acute infection (n=12). ENT = ear, nose, throat; MSK = musculoskeletal.

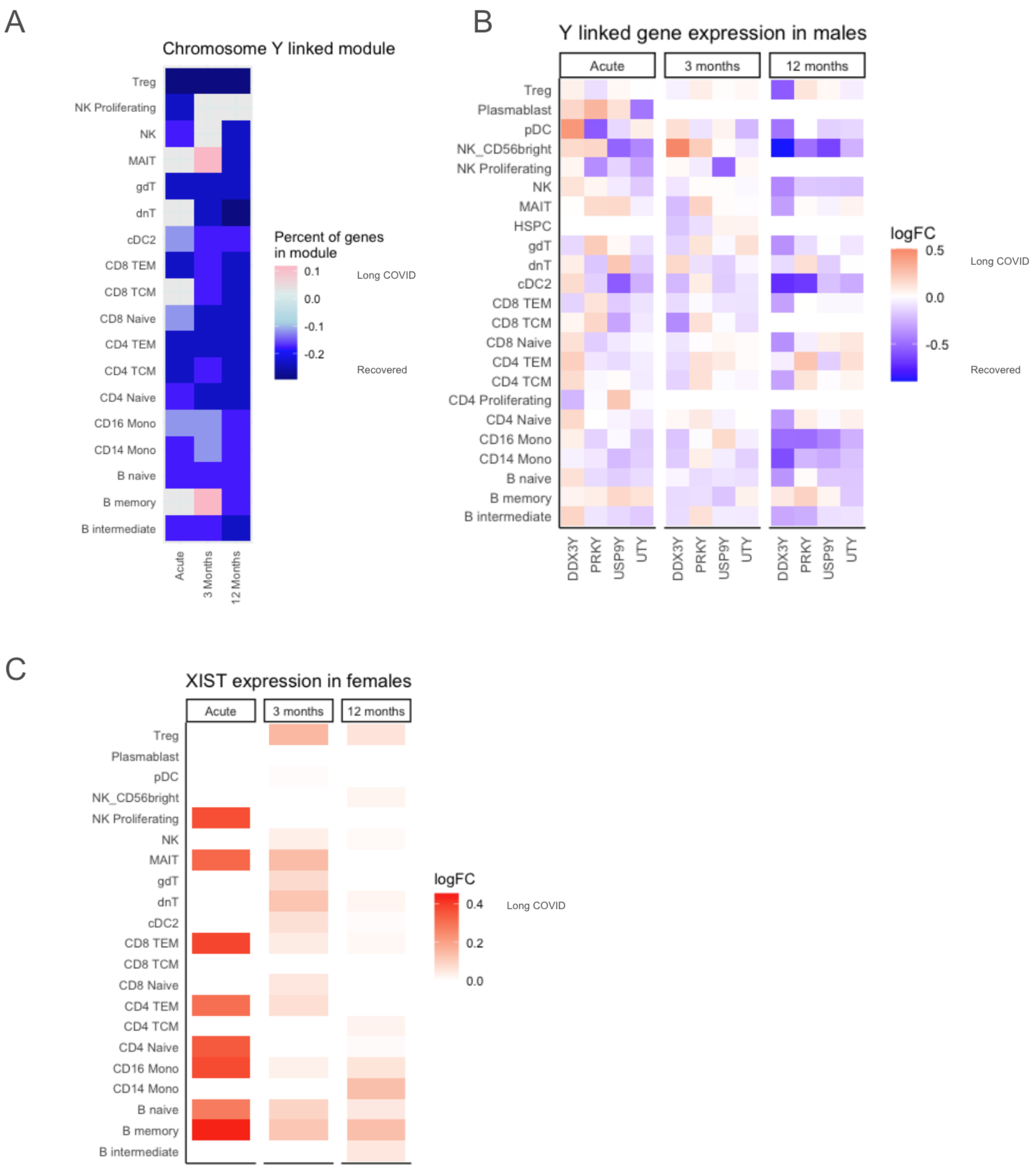


**Fig. S2. "Chromosome Y linked" module during acute infection and convalescence. (A)** In both sexes combined, the "chromosome Y linked" BTM was downregulated in almost all cell types in those with LC. Percentage of genes in the module with absolute log2FC >0.25 is indicated by tile color in the heatmap, with red indicating higher expression in LC and blue indicating lower expression. **(B)** Pseudobulk expression of the four most variable Y linked genes in males at acute infection as well as 3 and 12 months post infection. Red indicates higher expression and blue indicates lower expression in LC. **(C)** Pseudobulk expression of the *XIST* gene in females at acute infection as well as 3 and 12 months post infection. Log2FC is indicated by the color bar on the right, with red indicating higher expression in LC.

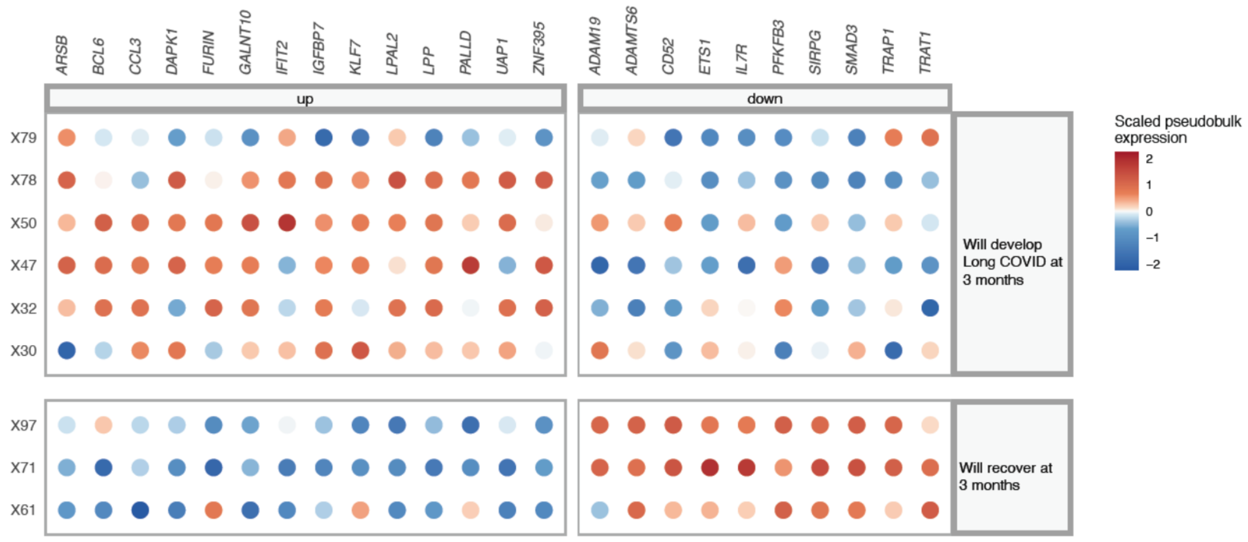


**Fig. S3. Patient-level pseudobulk expression of target genes for proliferating NK cells in males during acute infection, divided by those who will develop LC vs. recover at 3 months post infection.** Individual subjects are shown on the y-axis, and dot color indicates scaled pseudobulk expression of the target gene.

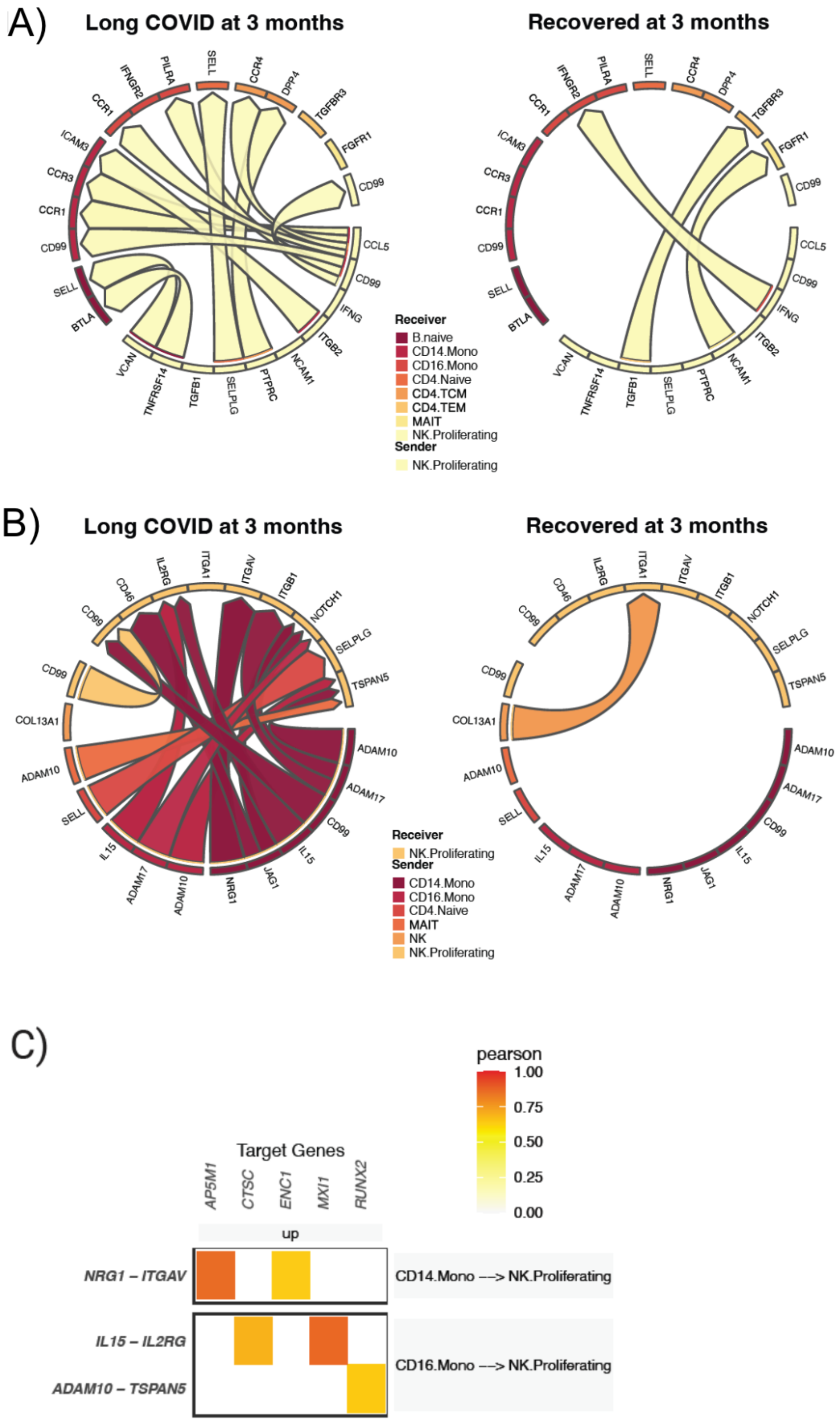


**Fig. S4. Proliferating NK cells in females during acute infection. (A-B)** Predicted ligand-receptor interactions **(A)** originating from proliferating NK cells, and **(B)** going toward proliferating NK cells, separated by presence of LC symptoms 3 months after acute infection versus resolution of symptoms by 3 months. **(C)** Target genes with increased expression in proliferating NK cells of acutely infected female subjects who will develop LC at 3 months post infection, with associated ligand-receptor interactions. Tile color in the heatmap indicates the pearson correlation coefficient of expression of the ligand-receptor pair and the target gene.

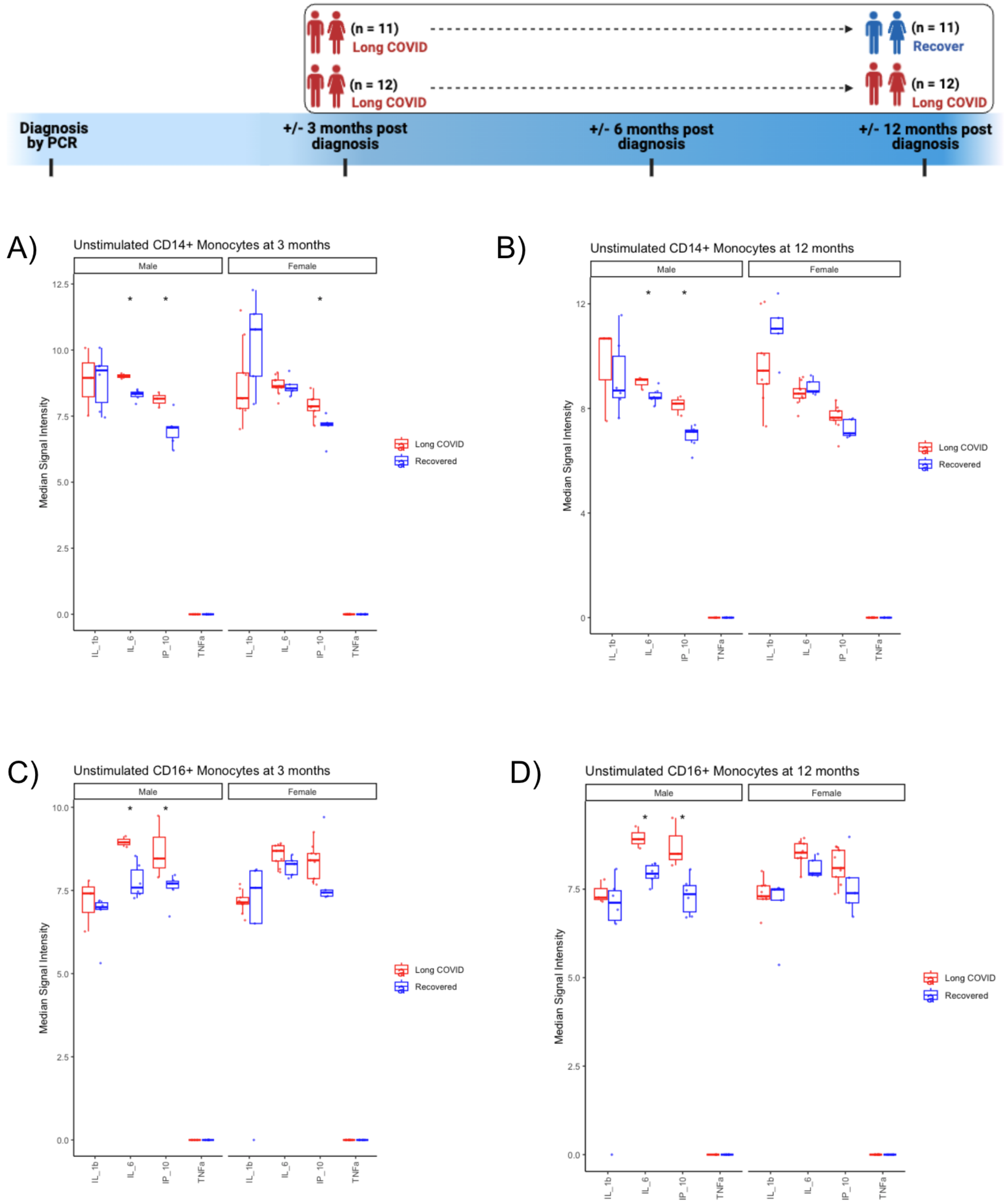


**Fig. S5. Increased inflammatory markers in monocytes in those with persistent LC. (A-B)** Median signal intensity by CyTOF of intracellular IL-1β, IL-6, IP-10 (aka CXCL10), and TNF**α** in unstimulated samples of $CD14^{+}$ monocytes at **(A)** 3 and **(B)** 12 months after acute infection, separated by sex. **(C-D)** Median signal intensity by CyTOF of IL-1β, IL-6, IP-10 (aka CXCL10), and TNF**α** in unsimulated samples of $CD16^+$ monocytes at **(C)** 3 and **(D)** 12 months after acute infection, separated by sex. * indicates unadjusted $p<0.05$ between those with persistent LC and recovery at 12 months.

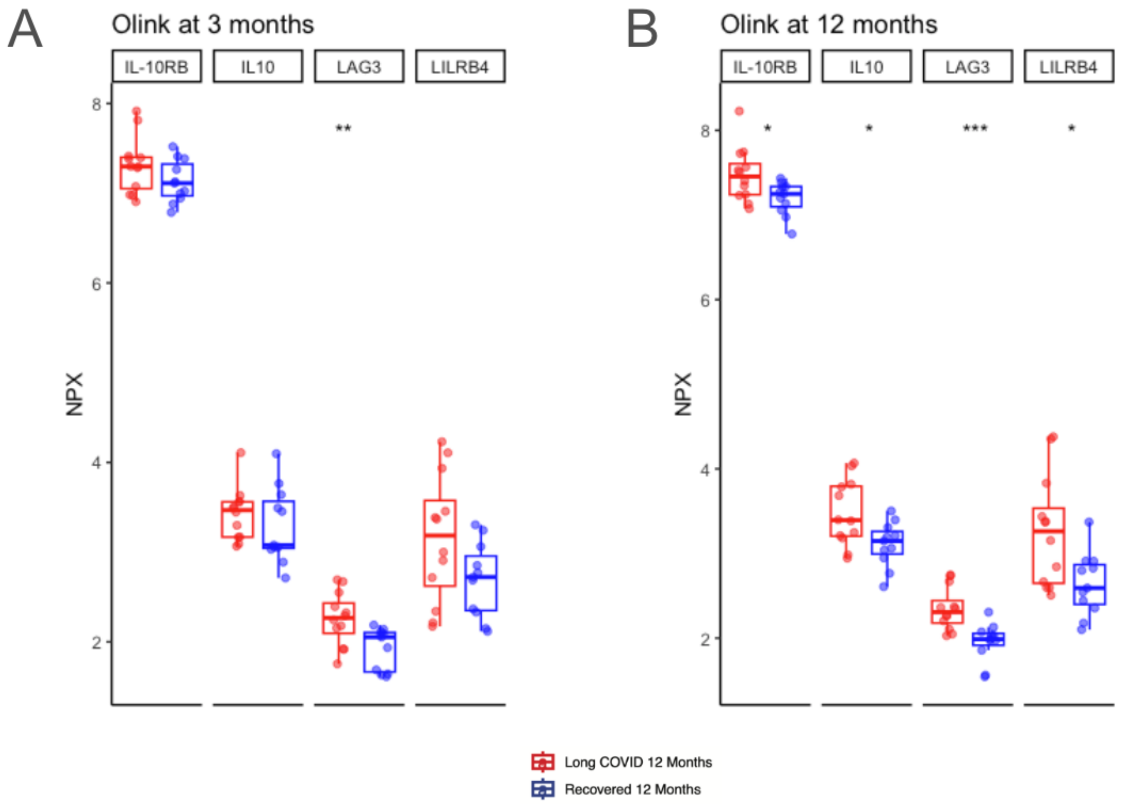


**Fig. S6. Plasma proteomic correlates of T cell exhaustion in sexes combined at (A) 3 months and (B) 12 months in those with persistent LC to 12 months compared to those with LC at 3 months but recovery by 12 months post infection.** * indicates unadjusted p<0.05, ** indicates unadjusted p<0.01, and *** indicates unadjusted p<0.001.

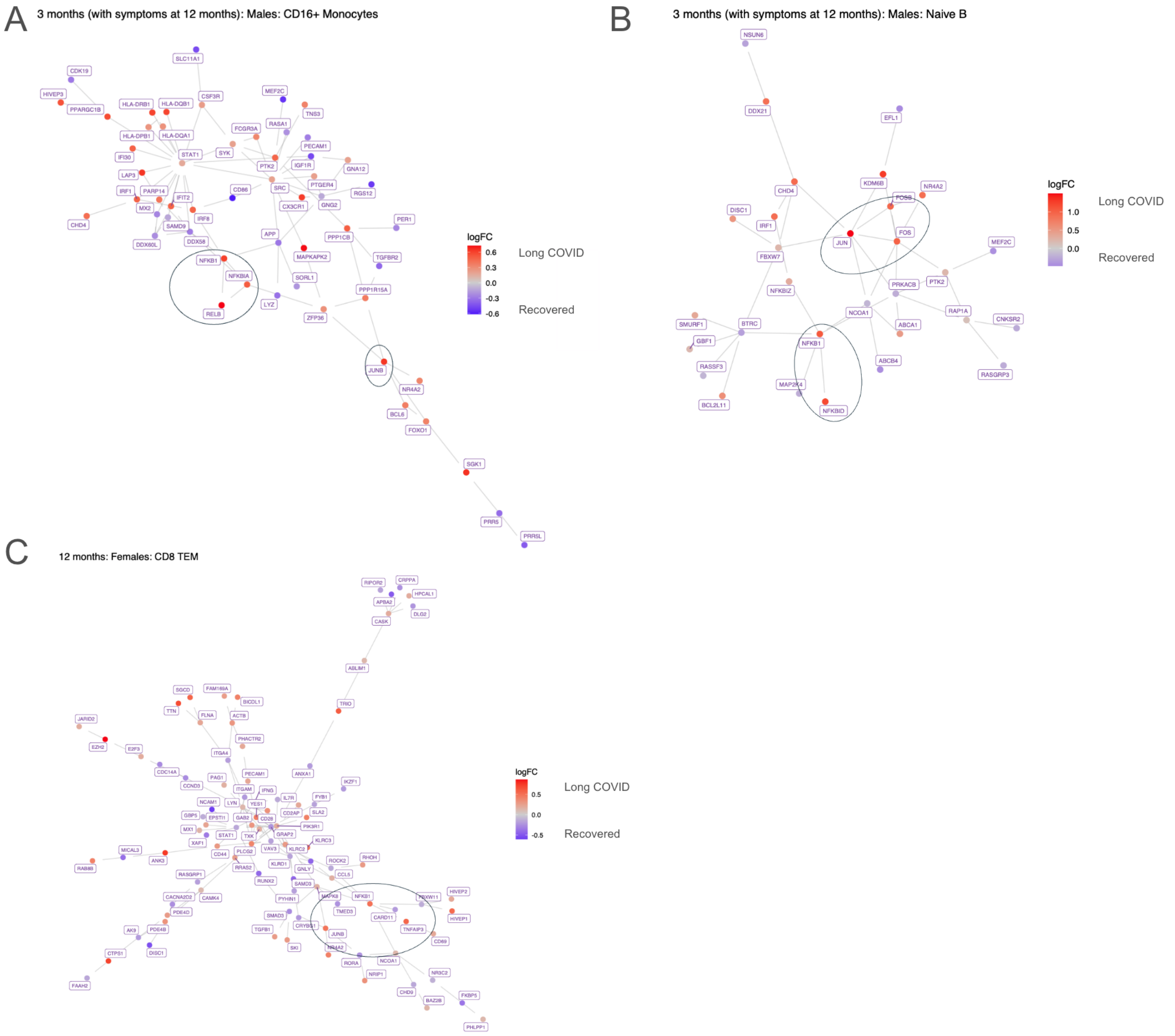


**Fig. S7. Increased NF-κB signaling in LC development and persistence. (A)** Gene network of CD16⁺ monocytes in males at 3 months post infection, comparing those with persistent LC to 12 months versus those who recover between 3 and 12 months. Red nodes indicate higher expression (log2FC) in those with persistent LC at 12 months, and blue nodes indicate lower expression. Increased expression of *NFkB1, RELB,* and *NFkBIA* are circled. **(B)** Gene network of naive B cells in males at 3 months post infection, comparing those with persistent LC to 12 months versus those who recover between 3 and 12 months. Red nodes indicate higher expression (log2FC) in those with persistent LC at 12 months, and blue nodes indicate lower expression. Increased expression of *NFkB1,* and *NFkBID* are circled. **(C)** Gene network of $CD8^+$ T effector memory (TEM) cells in females at 12 months post infection, comparing those with persistent LC at 12 months versus those who recovered by 12 months. Red nodes indicate higher expression (log2FC) in those with persistent LC at 12 months, and blue nodes indicate lower expression. Increased expression of *NFkB1* is circled.

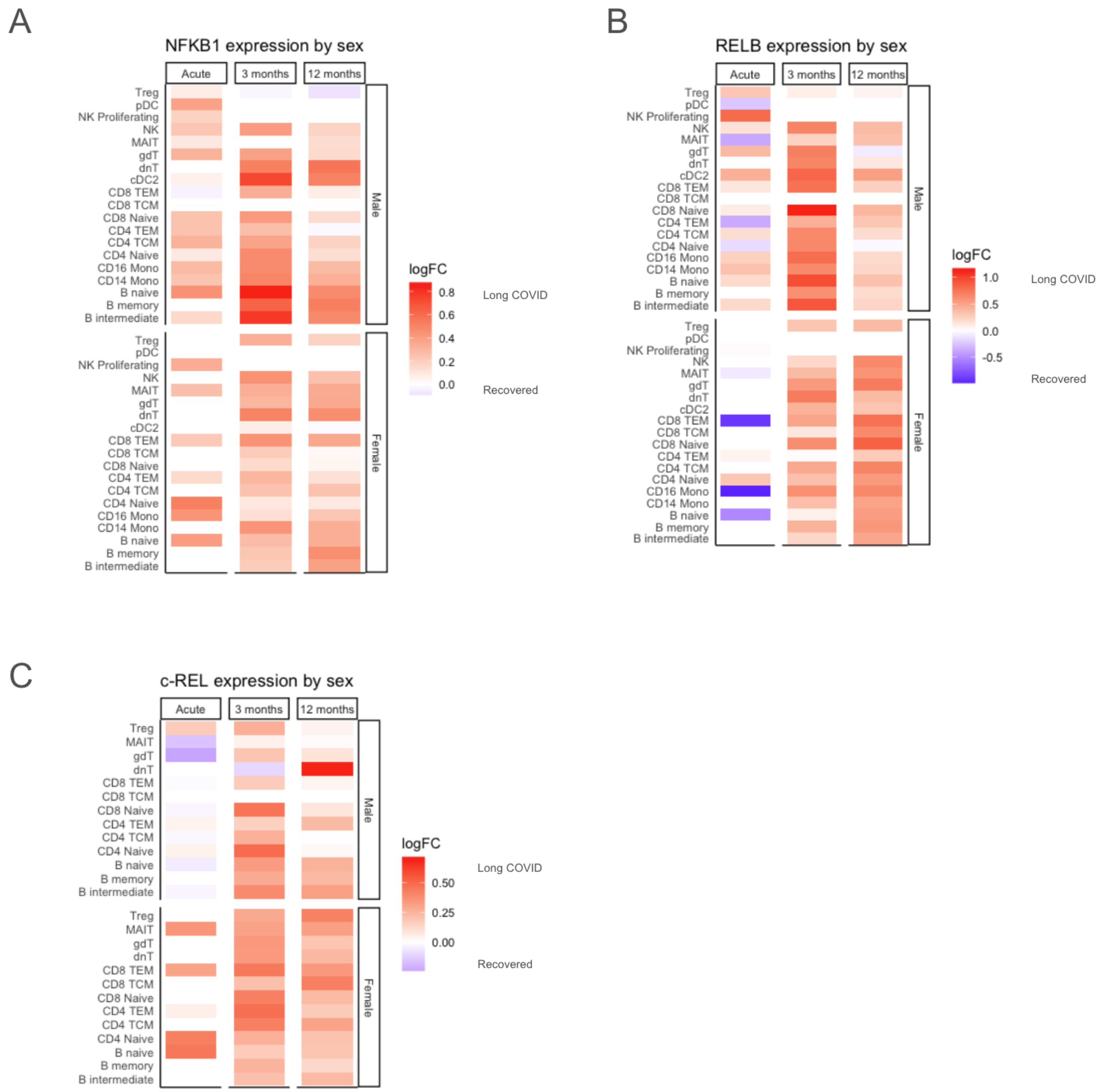


**Fig. S8. Increased NF-κB signaling in LC development and persistence is similar between sexes. (A-B)** Higher pseudobulk expression of **(A)** *NFkB1* and **(B)** *RELB* across many cell types is associated with LC development after acute infection and persistence for 12 months, separated by sex. **(C)** Increased pseudobulk *c-REL* expression across T and B lymphocytes is associated with LC development after acute infection and persistence for 12 months, separated by sex. Red tiles indicate higher expression (log2FC) in LC and blue tiles indicate lower expression.

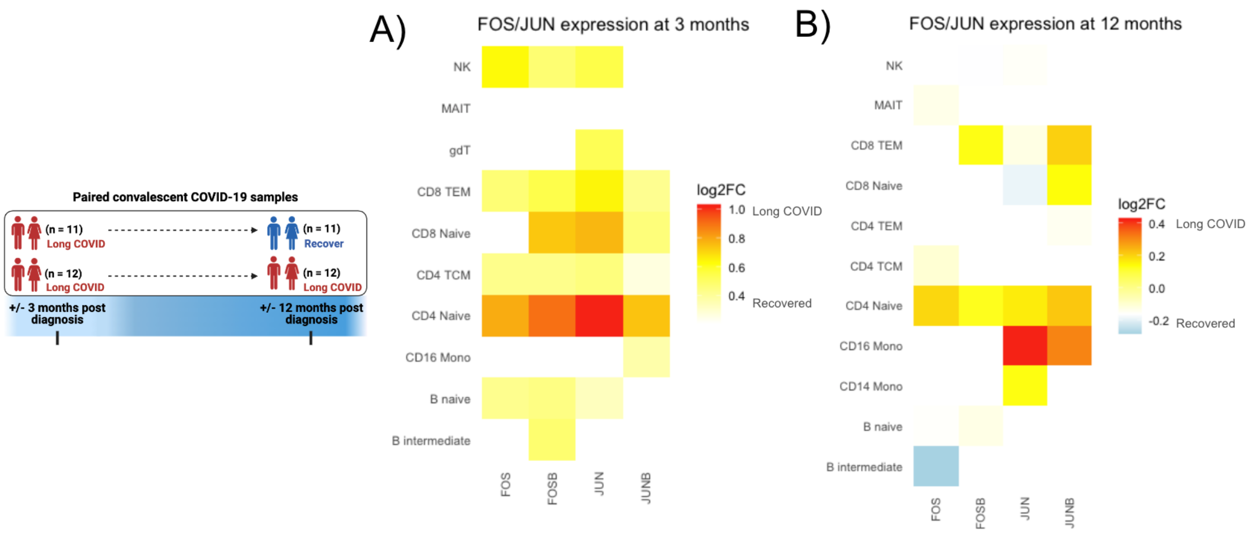


**Fig. S9. Higher pseudobulk expression of AP-1 transcription factor genes is associated with LC persistence. (A-B)** *FOS, FOSB, JUN,* and *JUNB* expression in lymphocytes at **(A)** 3 months and **(B)** 12 months, comparing those with persistent LC to 12 months and those who recover between 3 and 12 months. Yellow/red tiles indicate higher expression (log2FC) in persistent LC and blue tiles indicate lower expression.

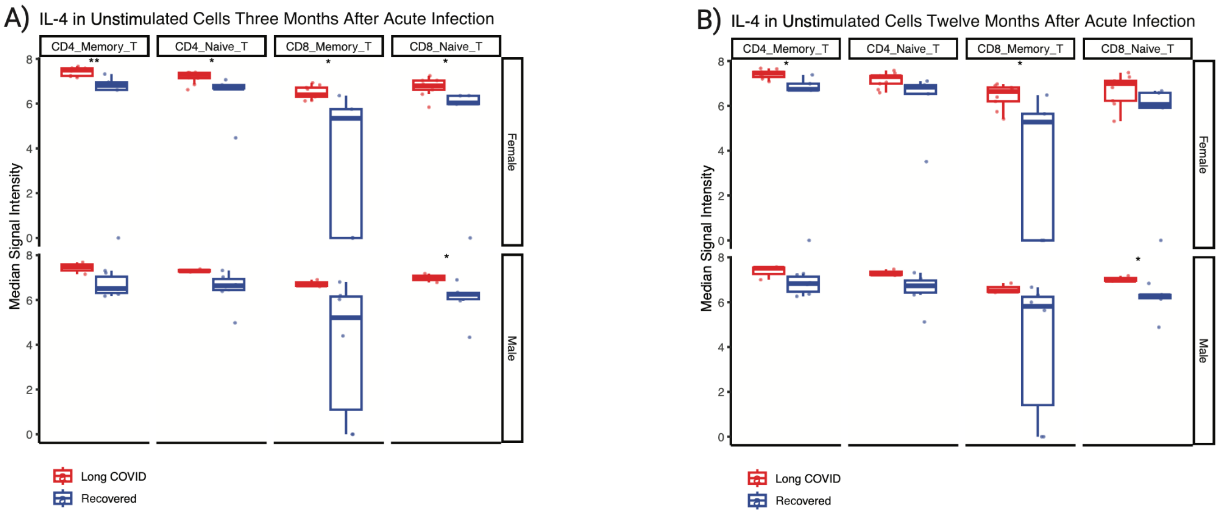


**Fig. S10. Intracellular IL-4 as measured by CyTOF is increased across T cell subsets in persistent LC.** IL-4 measured at **(A)** 3 months and **(B)** 12 months post infection, comparing those with LC at 3 and 12 months versus those with LC at 3 months but recovery by 12 months, separated by sex. * indicates p<0.05, ** indicates p<0.01.

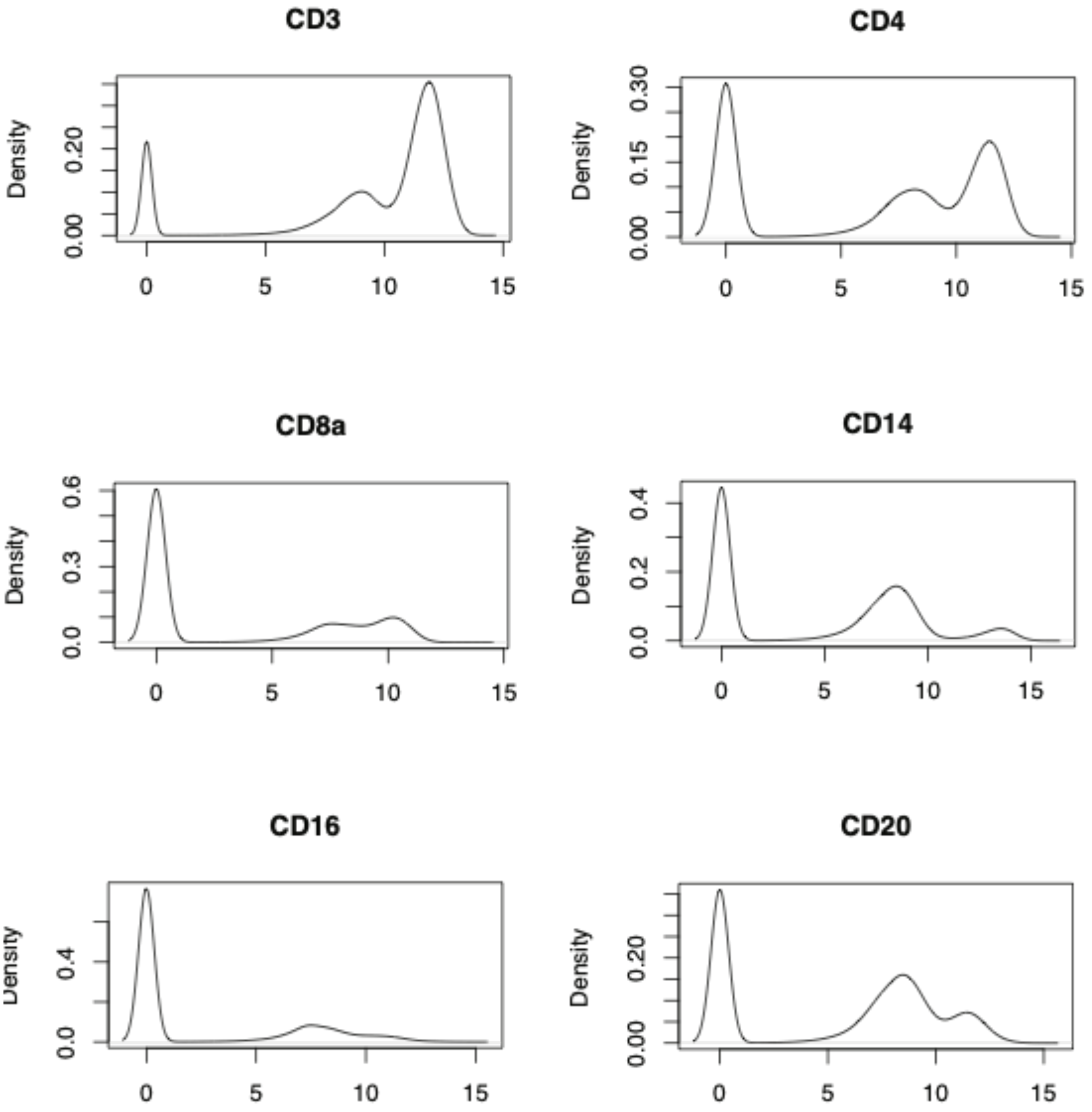


**Fig. S11. Density plots of example CyTOF markers, demonstrating multimodality with arsinch transformation cofactor of 0.001.**

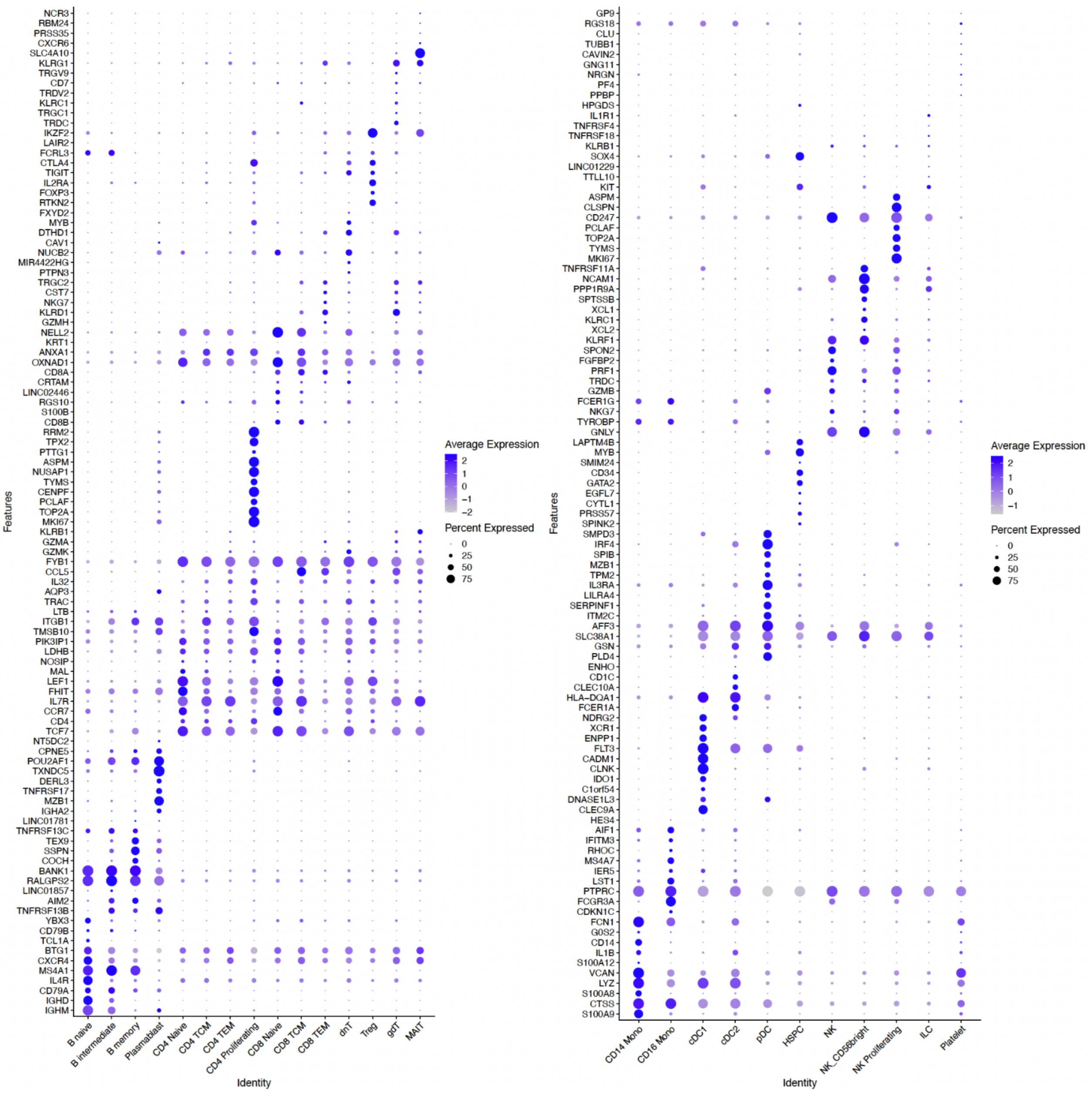


**Fig. S12. Canonical gene expression for cell type annotation in scRNAseq data.** Dot plot shows average and percent expression of annotation markers by cell type.

### SUPPLEMENTARY TABLES

| | Total (N=45) | Recovered (N=9) | Long COVID (N=36) | Female Recovered (N=4) | Female Long COVID (N=20) | Male Recovered (N=5) | Male Long COVID (N=16) |
|---|---|---|---|---|---|---|---|
| **Sex** | | | | | | | |
| Female | 24 (53.3%) | 4 (44.4%) | 20 (55.6%) | 4 (100%) | 20 (100%) | 0 (0%) | 0 (0%) |
| Male | 21 (46.7%) | 5 (55.6%) | 16 (44.4%) | 0 (0%) | 0 (0%) | 5 (100%) | 16 (100%) |
| **Age (years)** | | | | | | | |
| Mean (SD) | 42.8 (12.3) | 41.7 (9.45) | 43.0 (13.0) | 41.8 (10.4) | 41.2 (12.2) | 41.6 (9.89) | 45.3 (14.1) |
| Median [Min, Max] | 43.0 [20.0, 67.0] | 42.0 [27.0, 53.0] | 43.0 [20.0, 67.0] | 45.5 [27.0, 49.0] | 40.0 [20.0, 63.0] | 41.0 [27.0, 53.0] | 43.0 [26.0, 67.0] |
| **Race** | | | | | | | |
| White | 36 (80.0%) | 5 (55.6%) | 31 (86.1%) | 1 (25.0%) | 19 (95.0%) | 4 (80.0%) | 12 (75.0%) |
| Asian | 7 (15.6%) | 3 (33.3%) | 4 (11.1%) | 2 (50.0%) | 1 (5.0%) | 1 (20.0%) | 3 (18.8%) |
| Black or African American | 1 (2.2%) | 0 (0%) | 1 (2.8%) | 0 (0%) | 0 (0%) | 0 (0%) | 1 (6.3%) |
| American Indian | 0 (0%) | 0 (0%) | 0 (0%) | 0 (0%) | 0 (0%) | 0 (0%) | 0 (0%) |
| Pacific Islander | 0 (0%) | 0 (0%) | 0 (0%) | 0 (0%) | 0 (0%) | 0 (0%) | 0 (0%) |
| Other | 0 (0%) | 0 (0%) | 0 (0%) | 0 (0%) | 0 (0%) | 0 (0%) | 0 (0%) |
| Missing | 1 (2.2%) | 1 (11.1%) | 0 (0%) | 1 (25.0%) | 0 (0%) | 0 (0%) | 0 (0%) |
| **Ethnicity** | | | | | | | |
| Not Hispanic or Latino | 27 (60.0%) | 8 (88.9%) | 19 (52.8%) | 4 (100%) | 8 (40.0%) | 4 (80.0%) | 11 (68.8%) |
| Hispanic or Latino | 18 (40.0%) | 1 (11.1%) | 17 (47.2%) | 0 (0%) | 12 (60.0%) | 1 (20.0%) | 5 (31.3%) |
| **Body Mass Index** | | | | | | | |
| Mean (SD) | 30.2 (5.87) | 31.3 (7.86) | 30.0 (5.37) | 34.0 (10.1) | 31.1 (6.12) | 29.0 (5.78) | 28.7 (4.04) |
| Median [Min, Max] | 29.0 [19.6, 45.1] | 27.8 [22.4, 44.6] | 29.4 [19.6, 45.1] | 34.1 [23.3, 44.6] | 30.4 [19.6, 45.1] | 27.2 [22.4, 38.0] | 28.3 [22.9, 36.9] |
| **Hospitalized** | | | | | | | |
| No | 36 (80.0%) | 8 (88.9%) | 28 (77.8%) | 4 (100%) | 17 (85.0%) | 4 (80.0%) | 11 (68.8%) |
| Yes | 9 (20.0%) | 1 (11.1%) | 8 (22.2%) | 0 (0%) | 3 (15.0%) | 1 (20.0%) | 5 (31.3%) |
| **Intensive Care Unit** | | | | | | | |
| No | 41 (91.1%) | 8 (88.9%) | 33 (91.7%) | 4 (100%) | 19 (95.0%) | 4 (80.0%) | 14 (87.5%) |
| Yes | 4 (8.9%) | 1 (11.1%) | 3 (8.3%) | 0 (0%) | 1 (5.0%) | 1 (20.0%) | 2 (12.5%) |
| **Intubated** | | | | | | | |
| No | 44 (97.8%) | 9 (100%) | 35 (97.2%) | 4 (100%) | 19 (95.0%) | 5 (100%) | 16 (100%) |
| Yes | 1 (2.2%) | 0 (0%) | 1 (2.8%) | 0 (0%) | 1 (5.0%) | 0 (0%) | 0 (0%) |
| **Treatment during acute COVID-19** | | | | | | | |
| Remdesivir | 8 (17.8%) | 1 (11.1%) | 7 (19.4%) | 0 (0%) | 2 (10.0%) | 1 (20.0%) | 5 (31.3%) |
| Peginterferon Lambda-1a | 13 (28.9%) | 2 (22.2%) | 11 (30.6%) | 1 (25.0%) | 5 (25.0%) | 1 (20.0%) | 6 (37.5%) |
| Favipiravir | 8 (17.8%) | 3 (33.3%) | 5 (13.9%) | 3 (75.0%) | 3 (15.0%) | 0 (0%) | 2 (12.5%) |
| None or placebo | 16 (35.6%) | 3 (33.3%) | 13 (36.1%) | 0 (0%) | 10 (50.0%) | 3 (60.0%) | 3 (18.8%) |

**Table S1. Demographics of overall cohort at 3 months post SARS-CoV-2 infection, separated by LC versus recovered symptom status at 3 months after acute infection.**

| | Total | Recovered | Long COVID | Female | | Male | |
|---|---|---|---|---|---|---|---|
| | (N=21) | (N=6) | (N=15) | Recovered (N=3) | Long COVID (N=9) | Recovered (N=3) | Long COVID (N=6) |
| **Sex** | | | | | | | |
| Female | 12 (57.1%) | 3 (50.0%) | 9 (60.0%) | 3 (100%) | 9 (100%) | 0 (0%) | 0 (0%) |
| Male | 9 (42.9%) | 3 (50.0%) | 6 (40.0%) | 0 (0%) | 0 (0%) | 3 (100%) | 6 (100%) |
| **Age (years)** | | | | | | | |
| Mean (SD) | 44.7 (12.5) | 41.8 (9.00) | 45.8 (13.7) | 39.3 (11.2) | 42.1 (13.4) | 44.3 (7.57) | 51.3 (13.3) |
| Median [Min, Max] | 42.0 [26.0, 65.0] | 41.5 [27.0, 53.0] | 47.0 [26.0, 65.0] | 42.0 [27.0, 49.0] | 36.0 [26.0, 63.0] | 41.0 [39.0, 53.0] | 54.0 [34.0, 65.0] |
| **Race** | | | | | | | |
| White | 19 (90.5%) | 4 (66.7%) | 15 (100%) | 1 (33.3%) | 9 (100%) | 3 (100%) | 6 (100%) |
| Asian | 2 (9.5%) | 2 (33.3%) | 0 (0%) | 2 (66.7%) | 0 (0%) | 0 (0%) | 0 (0%) |
| Black or African American | 0 (0%) | 0 (0%) | 0 (0%) | 0 (0%) | 0 (0%) | 0 (0%) | 0 (0%) |
| American Indian | 0 (0%) | 0 (0%) | 0 (0%) | 0 (0%) | 0 (0%) | 0 (0%) | 0 (0%) |
| Pacific Islander | 0 (0%) | 0 (0%) | 0 (0%) | 0 (0%) | 0 (0%) | 0 (0%) | 0 (0%) |
| Other | 0 (0%) | 0 (0%) | 0 (0%) | 0 (0%) | 0 (0%) | 0 (0%) | 0 (0%) |
| **Ethnicity** | | | | | | | |
| Not Hispanic or Latino | 16 (76.2%) | 6 (100%) | 10 (66.7%) | 3 (100%) | 4 (44.4%) | 3 (100%) | 6 (100%) |
| Hispanic or Latino | 5 (23.8%) | 0 (0%) | 5 (33.3%) | 0 (0%) | 5 (55.6%) | 0 (0%) | 0 (0%) |
| **Body Mass Index** | | | | | | | |
| Mean (SD) | 30.8 (6.22) | 31.2 (6.64) | 30.6 (6.28) | 30.5 (8.86) | 31.5 (7.55) | 31.9 (5.48) | 29.4 (4.04) |
| Median [Min, Max] | 29.8 [19.6, 45.1] | 29.2 [23.3, 40.4] | 29.8 [19.6, 45.1] | 27.8 [23.3, 40.4] | 29.8 [19.6, 45.1] | 30.6 [27.2, 38.0] | 29.4 [23.8, 35.8] |
| **Hospitalized** | | | | | | | |
| No | 20 (95.2%) | 6 (100%) | 14 (93.3%) | 3 (100%) | 8 (88.9%) | 3 (100%) | 6 (100%) |
| Yes | 1 (4.8%) | 0 (0%) | 1 (6.7%) | 0 (0%) | 1 (11.1%) | 0 (0%) | 0 (0%) |
| **Intensive Care Unit** | | | | | | | |
| No | 20 (95.2%) | 6 (100%) | 14 (93.3%) | 3 (100%) | 8 (88.9%) | 3 (100%) | 6 (100%) |
| Yes | 1 (4.8%) | 0 (0%) | 1 (6.7%) | 0 (0%) | 1 (11.1%) | 0 (0%) | 0 (0%) |
| **Intubated** | | | | | | | |
| No | 20 (95.2%) | 6 (100%) | 14 (93.3%) | 3 (100%) | 8 (88.9%) | 3 (100%) | 6 (100%) |
| Yes | 1 (4.8%) | 0 (0%) | 1 (6.7%) | 0 (0%) | 1 (11.1%) | 0 (0%) | 0 (0%) |
| **Treatment during acute COVID-19** | | | | | | | |
| Remdesivir | 1 (4.8%) | 0 (0%) | 1 (6.7%) | 0 (0%) | 1 (11.1%) | 0 (0%) | 0 (0%) |
| Peginterferon Lambda-1a | 11 (52.4%) | 2 (33.3%) | 9 (60.0%) | 1 (33.3%) | 4 (44.4%) | 1 (33.3%) | 5 (83.3%) |
| Favipiravir | 5 (23.8%) | 2 (33.3%) | 3 (20.0%) | 2 (66.7%) | 2 (22.2%) | 0 (0%) | 1 (16.7%) |
| None or placebo | 4 (19.0%) | 2 (33.3%) | 2 (13.3%) | 0 (0%) | 2 (22.2%) | 2 (66.7%) | 0 (0%) |

**Table S2. Demographics of study participants with acute COVID-19 samples based on LC vs. recovery status at 3 months post infection.**

| Differentially expressed gene (DEG) | Long COVID (3 mo) | Cell Type of DEG | Function of Encoded Protein |
|---|---|---|---|
| XIST | ↑ | B intermediate, B naïve, CD4 naïve, CD4 TCM, CD4 TEM, CD8 naïve, CD8 TEM, gdT, MAIT, Treg, NK, cDC2, pDC, CD14 mono, CD16 mono | Involved in X chromosome inactivation; regulated by TGF-β/BMP signaling |
| DDX3Y | ↓ | B intermediate, B naïve, CD4 naïve, CD4 TCM, CD4 TEM, CD8 naïve, CD8 TEM, gdT, MAIT, Treg, NK, cDC2, pDC, CD14 mono, CD16 mono | RNA helicase; may enhance IFNB1 expression via IRF3/IRF7 pathway |
| UTY/KDM6C | ↓ | B intermediate, B naïve, CD4 naïve, CD4 TCM, CD4 TEM, CD8 naïve, CD8 TEM, gdT, MAIT, Treg, NK, pDC, CD14 mono, CD16 mono | minor histocompatibility antigen important for B and T cell development/maturation |
| KDM5D | ↓ | Treg, pDC | minor histocompatibility antigen; suppresses stimulator of interferon genes (STING) |
| USP9Y | ↓ | B intermediate, B naïve, CD4 naïve, CD4 TCM, CD4 TEM, CD8 naïve, CD8 TEM, gdT, MAIT, Treg, NK, pDC, CD14 mono | ubiquitin-specific peptidase; essential component of TGF-β/BMP signaling |
| PRKY | ↓ | CD4 naïve, CD4 TCM, CD4 TEM, CD8 naïve, CD8 TEM, gdT, MAIT, Treg, NK, CD16 mono | pseudogene |

**Table S3. Characteristics of the differentially expressed genes driving the "chromosome Y linked" BTM.**

| | Total (N=23) | Recovered (N=11) | Long COVID (N=12) | Female Recovered (N=5) | Female Long COVID (N=9) | Male Recovered (N=6) | Male Long COVID (N=3) |
|---|---|---|---|---|---|---|---|
| **Sex** | | | | | | | |
| Female | 14 (60.9%) | 5 (45.5%) | 9 (75.0%) | 5 (100%) | 9 (100%) | 0 (0%) | 0 (0%) |
| Male | 9 (39.1%) | 6 (54.5%) | 3 (25.0%) | 0 (0%) | 0 (0%) | 6 (100%) | 3 (100%) |
| **Age (years)** | | | | | | | |
| Mean (SD) | 41.2 (12.7) | 37.3 (9.90) | 44.8 (14.3) | 38.8 (12.5) | 42.1 (13.4) | 36.0 (8.20) | 53.0 (16.6) |
| Median [Min, Max] | 43.0 [20.0, 65.0] | 43.0 [20.0, 49.0] | 41.5 [26.0, 65.0] | 46.0 [20.0, 49.0] | 36.0 [26.0, 63.0] | 37.0 [26.0, 44.0] | 60.0 [34.0, 65.0] |
| **Race** | | | | | | | |
| White | 20 (87.0%) | 8 (72.7%) | 12 (100%) | 4 (80.0%) | 9 (100%) | 4 (66.7%) | 3 (100%) |
| Asian | 3 (13.0%) | 3 (27.3%) | 0 (0%) | 1 (20.0%) | 0 (0%) | 2 (33.3%) | 0 (0%) |
| Black or African American | 0 (0%) | 0 (0%) | 0 (0%) | 0 (0%) | 0 (0%) | 0 (0%) | 0 (0%) |
| American Indian | 0 (0%) | 0 (0%) | 0 (0%) | 0 (0%) | 0 (0%) | 0 (0%) | 0 (0%) |
| Pacific Islander | 0 (0%) | 0 (0%) | 0 (0%) | 0 (0%) | 0 (0%) | 0 (0%) | 0 (0%) |
| Other | 0 (0%) | 0 (0%) | 0 (0%) | 0 (0%) | 0 (0%) | 0 (0%) | 0 (0%) |
| **Ethnicity** | | | | | | | |
| Not Hispanic or Latino | 12 (52.2%) | 5 (45.5%) | 7 (58.3%) | 2 (40.0%) | 4 (44.4%) | 3 (50.0%) | 3 (100%) |
| Hispanic or Latino | 11 (47.8%) | 6 (54.5%) | 5 (41.7%) | 3 (60.0%) | 5 (55.6%) | 3 (50.0%) | 0 (0%) |
| **Body Mass Index** | | | | | | | |
| Mean (SD) | 29.6 (5.97) | 28.4 (4.94) | 30.7 (6.80) | 29.9 (6.27) | 31.5 (7.55) | 27.2 (3.65) | 28.4 (4.01) |
| Median [Min, Max] | 29.8 [19.6, 45.1] | 28.2 [21.9, 38.9] | 30.1 [19.6, 45.1] | 29.8 [21.9, 38.9] | 29.8 [19.6, 45.1] | 27.6 [22.9, 32.0] | 30.4 [23.8, 31.0] |
| **Hospitalized** | | | | | | | |
| No | 19 (82.6%) | 8 (72.7%) | 11 (91.7%) | 5 (100%) | 8 (88.9%) | 3 (50.0%) | 3 (100%) |
| Yes | 4 (17.4%) | 3 (27.3%) | 1 (8.3%) | 0 (0%) | 1 (11.1%) | 3 (50.0%) | 0 (0%) |
| **Intensive Care Unit** | | | | | | | |
| No | 21 (91.3%) | 10 (90.9%) | 11 (91.7%) | 5 (100%) | 8 (88.9%) | 5 (83.3%) | 3 (100%) |
| Yes | 2 (8.7%) | 1 (9.1%) | 1 (8.3%) | 0 (0%) | 1 (11.1%) | 1 (16.7%) | 0 (0%) |
| **Intubated** | | | | | | | |
| No | 22 (95.7%) | 11 (100%) | 11 (91.7%) | 5 (100%) | 8 (88.9%) | 6 (100%) | 3 (100%) |
| Yes | 1 (4.3%) | 0 (0%) | 1 (8.3%) | 0 (0%) | 1 (11.1%) | 0 (0%) | 0 (0%) |
| **Treatment during acute COVID-19** | | | | | | | |
| Remdesivir | 4 (17.4%) | 3 (27.3%) | 1 (8.3%) | 0 (0%) | 1 (11.1%) | 3 (50.0%) | 0 (0%) |
| Peginterferon Lambda-1a | 8 (34.8%) | 1 (9.1%) | 7 (58.3%) | 0 (0%) | 4 (44.4%) | 1 (16.7%) | 3 (100%) |
| Favipiravir | 4 (17.4%) | 2 (18.2%) | 2 (16.7%) | 1 (20.0%) | 2 (22.2%) | 1 (16.7%) | 0 (0%) |
| None or placebo | 7 (30.4%) | 5 (45.5%) | 2 (16.7%) | 4 (80.0%) | 2 (22.2%) | 1 (16.7%) | 0 (0%) |

**Table S4. Demographics of study participants with paired 3 and 12 month samples based on persistent Long COVID vs. recovery status at 12 months post infection.**

| Isotope | Antibody | Clone | Company |
|---|---|---|---|
| 89Y | CD45 | HI30 | Fluidigm/Standard BioTools (pre-conjugated) |
| 141Pr | CD57 | HNK-1 | BioLegend |
| 142Nd | CD19 | HIB19 | Fluidigm/Standard BioTools (pre-conjugated) |
| 143Nd | CD45RA | HI100 | Fluidigm/Standard BioTools (pre-conjugated) |
| 144Nd | CD38 | HIT2 | Fluidigm/Standard BioTools (pre-conjugated) |
| 145Nd | CD4 | RPA-T4 | Fluidigm/Standard BioTools (pre-conjugated) |
| 146Nd | NKB1 (KIR3DL1) | DX9 | R&D Systems |
| 147Sm | CD8a | RPA-T8 | BioLegend |
| 148Nd | CD14 | RMO52 | Fluidigm/Standard BioTools (pre-conjugated) |
| 150Nd | CD28 | CD28.2 | BioLegend |
| 151Eu | CD123 | 6H6 | Fluidigm/Standard BioTools (pre-conjugated) |
| 152Sm | TCRγδ | 11F2 | Fluidigm/Standard BioTools (pre-conjugated) |
| 153Eu | CD20 | 2H7 | BioLegend |
| 154Sm | CD3 | UCHT1 | Fluidigm/Standard BioTools (pre-conjugated) |
| 155Gd | CD158b (KIR2DL2/L3) | DX27 | BioLegend |
| 157Gd | CD294/CRTH2 | BM16 | BioLegend |
| 159Tb | CD11c | Bu15 | Fluidigm/Standard BioTools (pre-conjugated) |
| 162Dy | CD25 | M-A251 | BioLegend |
| 163Dy | CD183/CXCR3 | G025H7 | Fluidigm/Standard BioTools (pre-conjugated) |
| 164Dy | CD161 | HP-3G10 | Fluidigm/Standard BioTools (pre-conjugated) |
| 166Er | CD197/CCR7 | G043H7 | BioLegend |
| 167Er | CD27 | O323 | Fluidigm/Standard BioTools (pre-conjugated) |

| | | | |
|---|---|---|---|
| 168Er | CD196/CCR6 | G034E3 | BioLegend |
| 171Yb | CD185/CXCR5 | RF8B2 | Fluidigm/Standard BioTools (pre-conjugated) |
| 172Yb | CD194/CCR4 | L291H4 | BioLegend |
| 173Yb | HLA-DR | L243 | Fluidigm/Standard BioTools (pre-conjugated) |
| 174Yb | CD127 | A019D5 | BioLegend |
| 176Yb | CD56 | NCAM16.2 | Fluidigm/Standard BioTools (pre-conjugated) |
| 209Bi | CD16 | 3G8 | Fluidigm/Standard BioTools (pre-conjugated) |

**Table S5. CyTOF surface staining panel.**

| Isotope | Antibody | Clone | Company |
|---|---|---|---|
| 149Sm | APC** | APC003 | BioLegend |
| 156Gd | IL-6 | MQ2-12A5 | Fluidigm/Standard BioTools (pre-conjugated) |
| 158Gd | IFNg | B27 | Fluidigm/Standard BioTools (pre-conjugated) |
| 160Gd | IL-4 | MP4-25D2 | BioLegend |
| 161Dy | IL-1b | JK1B-1 | BioLegend |
| 165Ho | CXCL10 | J034D6 | BioLegend |
| 169Tm | IL-17A | BL168 | Fluidigm/Standard BioTools (pre-conjugated) |
| 170Er | TNFa | MAb11 | BioLegend |
| 175Lu | Perforin | B-D48 | Fluidigm/Standard BioTools (pre-conjugated) |
| ***NOTE: Anti-APC-149Sm is to target anti-human APC - CD107a (Biolegend, clone H4A3)* | | | |

**Table S6. CyTOF intracellular staining panel.**

| Time Point | Sex | Cell type | Sender or Receiver | Ligand | Receptor |
|---|---|---|---|---|---|
| Acute infection | Male | Proliferating NK | receiver | CRTAM | ILDR2 |
| Acute infection | Male | Proliferating NK | receiver | COL4A4 | ITGB1 |
| Acute infection | Male | Proliferating NK | sender | TGFB1 | APP |
| Acute infection | Male | Proliferating NK | sender | NETO2 | IL18R1 |
| Acute infection | Female | Proliferating NK | receiver | TGFB1 | ITGA5 |
| Acute infection | Female | Proliferating NK | receiver | SPON2 | ITGA5 |
| Acute infection | Female | Proliferating NK | receiver | ENG | ITGAV |
| Acute infection | Female | Proliferating NK | receiver | FAM3C | ADGRG5 |
| Acute infection | Female | Proliferating NK | receiver | COL4A4 | ITGA1 |
| Acute infection | Female | Proliferating NK | receiver | COL4A4 | ITGA2 |
| Acute infection | Female | Proliferating NK | receiver | COL4A4 | CD47 |
| Acute infection | Female | Proliferating NK | receiver | COL4A4 | ITGB1 |
| Acute infection | Female | Proliferating NK | receiver | COL4A4 | ITGAV |
| Acute infection | Female | Proliferating NK | sender | PTPRC | CD247 |
| Acute infection | Female | Proliferating NK | sender | PTPRC | CD4 |
| Acute infection | Female | Proliferating NK | sender | ITGB2 | CD82 |
| Acute infection | Female | Proliferating NK | sender | FAM3C | ADGRG5 |
| Acute infection | Female | Proliferating NK | sender | CLEC2B | KLRB1 |
| Acute infection | Female | Proliferating NK | sender | CALR | TAP1 |
| Acute infection | Female | Proliferating NK | sender | CALR | TAP2 |
| Acute infection | Female | Proliferating NK | sender | CALM3 | PDE1B |
| Acute infection | Female | Proliferating NK | sender | ITGAL | CD226 |
| Acute infection | Female | Proliferating NK | sender | CALR | ITGAV |
| Acute infection | Female | Proliferating NK | sender | PAM | DPP4 |
| Acute infection | Male | CD14 mono | receiver | TGFB1 | SDC2 |
| Acute infection | Male | CD14 mono | receiver | CD55 | ADGRE2 |

| | | | | | |
|---|---|---|---|---|---|
| Acute infection | Male | CD14 mono | receiver | CD55 | CR1 |
| Acute infection | Male | CD14 mono | receiver | HBEGF | CD9 |
| Acute infection | Male | CD14 mono | receiver | CD55 | ADGRE2 |
| Acute infection | Male | CD14 mono | receiver | SPON2 | ITGAM |
| Acute infection | Male | CD14 mono | receiver | BMP8B | BMPR2 |
| Acute infection | Male | CD14 mono | receiver | BMP8B | PLAUR |
| Acute infection | Male | CD14 mono | receiver | LILRA6 | CD300E |
| Acute infection | Male | CD16 mono | receiver | CD55 | ADGRE2 |
| Acute infection | Male | CD16 mono | receiver | LRP1B | PLAUR |
| Acute infection | Male | CD16 mono | receiver | CD55 | CR1 |
| Acute infection | Male | CD16 mono | receiver | NECTIN2 | PVR |
| Acute infection | Male | CD16 mono | receiver | JAM3 | ITGB2 |
| Acute infection | Male | CD16 mono | receiver | JAM3 | ITGB2 |
| Acute infection | Male | CD16 mono | receiver | S100A8 | ITGB2 |
| Acute infection | Male | CD16 mono | receiver | TGFB2 | ENG |
| Acute infection | Male | CD16 mono | receiver | EMC1 | PRLR |
| Acute infection | Male | CD16 mono | receiver | ANGPT1 | ITGA5 |
| Acute infection | Male | CD16 mono | receiver | JAM3 | ITGB2 |
| Acute infection | Female | CD14 mono | receiver | ITGB2 | CD82 |
| Acute infection | Female | CD14 mono | receiver | HLA.DRA | CD53 |
| Acute infection | Female | CD14 mono | receiver | TSPAN3 | ADAM10 |
| Acute infection | Female | CD14 mono | receiver | HLA.DRA | CD82 |
| Acute infection | Female | CD14 mono | receiver | CD300C | PTPRO |
| Acute infection | Female | CD14 mono | receiver | HLA.DRA | CD82 |
| Acute infection | Female | CD14 mono | receiver | HLA.DRB5 | CD4 |
| Acute infection | Female | CD14 mono | receiver | HLA.DRA | CD63 |
| Acute infection | Female | CD14 mono | receiver | HLA.C | LILRB1 |

| | | | | | |
|---|---|---|---|---|---|
| Acute infection | Female | CD16 mono | receiver | FCN1 | LRP1 |
| Acute infection | Female | CD16 mono | receiver | SEMA4A | PLXNA2 |
| Acute infection | Female | CD16 mono | receiver | ADAM10 | TSPAN14 |
| Acute infection | Female | CD16 mono | receiver | HLA.DQA2 | CD4 |
| Acute infection | Female | CD16 mono | receiver | COL4A4 | ITGAV |
| Acute infection | Female | CD16 mono | receiver | TSPAN33 | ADAM10 |
| Acute infection | Female | CD16 mono | receiver | NCAM1 | NPTN |
| Acute infection | Female | CD16 mono | receiver | HLA.C | LILRB1 |
| Acute infection | Female | CD16 mono | receiver | LAMB3 | CD151 |
| Acute infection | Female | CD16 mono | receiver | SIGLEC1 | PTPRC |
| Acute infection | Female | CD16 mono | receiver | COL4A4 | CD93 |
| Acute infection | Female | CD16 mono | receiver | ADAM17 | NOTCH2 |

**Table S7. MultiNicheNet ligand-receptor pair curation.** Pairs that were excluded due to lack of experimental evidence or the interaction is known to occur within the same cell.
